# Characterization of Host Cell Cytosol-filled Vesicles in the Human Malaria Parasite *Plasmodium falciparum*

**DOI:** 10.64898/2026.09.10.750675

**Authors:** Anna-Lena Roßmann, Jana Dröge, Hanyu Wang, Charlotte Duda, James Zhen, Ricarda Sabitzki, Katharina Höhn, Guilherme B. Farias, Andres Guillen-Samander, Tim-Wolf Gilberger, Chi-Minh Ho, Tobias Spielmann

## Abstract

Endocytosis of host cell cytosol is a key process in malaria blood stages. The endocytosed material consists mostly of haemoglobin (Hb) and is transported to the parasite’s digestive vacuole (DV), where it is degraded. However, the endosomal transport pathway of the parasite is not well defined. A number of proteins are known that – when inactivated – lead to the appearance of Hb-filled vesicles (HbVs) in the parasite cytoplasm and prevent Hb from arriving in the DV, indicating they play a role in endosomal transport. However, despite the prominence of these HbVs, their morphology, molecular composition, and relationship to the endosomal transport pathway remain poorly understood.

Here we carried out a morphological and surface proteome study of HbVs. CryoET showed HbVs are coatless, double membraned vesicles with a very narrow inter-membrane space devoid of larger protein densities. HbV surface proteomes generated by BioIDs most prominently detected DV proteases. Halo-based tracking of one of these DV protease confirmed it as a bona fide HbV cargo. The BioID also identified a number of vesicle trafficking proteins, including PfTBC8 and Rab11 proteins. Functional analysis of PfTBC8 showed its importance for the transport of Hb to the DV. This was due to a novel phenotype characterized by accumulation of diverse Hb-filled structures in the parasite cytosol and gradual decline of DV protease trafficking. PfTBC8 DiQ-BioID detected Rab11a and PfTBC8 inactivation altered Rab11a localization, suggesting it may be a Rab11a effector and that the new endosomal transport phenotype may result from a recycling defect. Together with the detection of VPS35 in the BioID and evidence from the CryoET of a disrupted inner HbV membrane in a proportion of vesicles, these findings support a model in which HbVs undergo maturation prior to fusion with the DV. Overall, our findings define morphological and molecular features of HbVs, identify protein cargo transported through this pathway, and reveal a previously unrecognized Rab-regulatory component required for endosomal trafficking in blood-stage *P. falciparum* parasites.

## Introduction

The protozoan parasite *Plasmodium falciparum* is responsible for the majority of malaria-related deaths worldwide and the major driver of the global malaria burden (WHO, 2025). *P. falciparum* parasites multiply in the human blood where they develop in red blood cells (RBCs). During this cycle the parasites develop within 48 h from the small ring form to the trophozoite stage and finally to the multinucleated schizont stage after which the host RBC ruptures, releasing short-lived merozoite stage parasites that rapidly invade new RBCs to initiate the next round of multiplication. The intracellular RBC stages endocytose host cell cytosol (HCC), a process essential for their development, providing both nutrients and physical space for parasite growth (Dalal and Klemba, 2015; Hanssen et al., 2012; Krugliak et al., 2002; Lew et al., 2003). The endocytosed HCC consists almost exclusively of hemoglobin (Hb). Reduced Hb uptake underlies the mechanism by which the parasite becomes partially resistant to the first line drug artemisinin (Behrens et al., 2021; Birnbaum et al, 2020; Yang et al) and digestion of the internalized hemoglobin is the target of different antimalarial drugs (Klonis et al., 2013; Nsanzabana, 2019), highlighting the importance of this pathway.

In model organisms, the architecture of the endosomal pathway and the molecular mechanisms of endosomal transport have been extensively characterized (Grant and Donaldson, 2009; Huotari and Helenius, 2011; Ungermann and Moeller, 2025; Gopaldass et al., 2024; Klumperman and Raposo, 2014). Following endocytosis, internalized cargo is delivered to the early endosome (EE), which functions as the primary sorting station of the endocytic system (Huotari and Helenius, 2011). From there, cargo is either recycled back to the plasma membrane or targeted for degradation via the late endosome (LE) and lysosome (Grant and Donaldson, 2009; Schmidt and Teis, 2012). Endosomal maturation, cargo sorting and vesicular fusion are coordinated by several conserved processes, including ESCRT-mediated membrane remodeling, HOPS/CORVET-mediated vesicle fusion, Rab GTPase switching, changes in the phosphoinositide composition, and retromer-dependent cargo sorting (Borchers et al., 2023; Sbrissa et al., 1999; Schleinitz et al., 2023; Schmidt and Teis, 2012; Seaman, 2012).

In mammalian cells, EEs - characterized by the presence of Rab5 and phosphatidylinositol 3-phosphate [PI(3)P] and - receive incoming cargo and serve as major sorting stations (Grant and Donaldson, 2009; Jovic et al., 2010; Naslavsky and Caplan, 2018). Endocytic vesicle fusion with EEs involves the CORVET tethering complex, which is recruited through Rab5 and promotes tethering and fusion with EE membranes (Perini et al., 2014). Recycling of Cargo at the EE either occurs by fast recycling via Rab4 or by slow recycling via the recycling endosome (RE) by Rab11 (Sönnichsen et al., 2000; Ullrich et al., 1996). Ubiquitinated cargo destined for degradation is concentrated on EEs and internalized into intraluminal vesicles by ESCRT-mediated membrane invagination, committing it to lysosomal degradation (Babst et al., 2002; Jovic et al., 2010; Saftig and Klumperman, 2009). During endosomal maturation, Rab5 is replaced by Rab7 (Kanai et al., 2001) through the Mon1–Ccz1 complex (Borchers et al., 2023; Rink et al., 2005). In parallel, PI(3)P on the early endosomal membrane is converted to PI(3,5)P₂ by the PI(3)P 5-kinase PIKfyve (Huotari and Helenius, 2011; Marat and Haucke, 2016). Together, these activities establish a molecular switch that drives progression from EEs to LEs that then fuse with the lysosome (Huotari and Helenius, 2011; Rink et al., 2005).

In contrast, the organization and dynamics of the endocytic pathway in *P. falciparum* remain poorly understood (Spielmann et al., 2020). Blood-stage parasites internalize HCC through a specialized structure termed the cytostome, an invagination of both parasite plasma membrane (PPM) and parasitophorous vacuolar membrane (PVM) (Aikawa, 1971; Aikawa et al., 1966; Langreth et al., 1978; Spielmann et al., 2020). The cytostome contains a constricted neck surrounded by proteins needed for endocytosis, including for instance KIC7 and Kelch13 (Birnbaum et al., 2020; Liffner et al., 2023; Spielmann et al., 2020; Tutor et al., 2023). How the cytostome internalises HCC and how the endocytosed material is transported to the parasite’s digestive vacuole (DV) where it is digested, is not well defined. Previous studies identified several proteins that cause an accumulation of HCC-filled vesicles (here referred to as Hb filled vesicles, HbVs) in the parasite cytoplasm when they are inactivated, including for instance a Rabenosyn5-like protein (RBNS5L), indicating they are involved in the transport of HbVs to the DV (Jonscher et al., 2019; Mukherjee et al., 2022; Sabitzki et al., 2024; Schmidt et al., 2023; Schmitz et al., 2026). Depolymerization of actin using cytochalasin D (CytD) equally leads to HbV accumulation (Lazarus et al., 2008; Milani et al., 2015; Smythe et al., 2007).

If cytostome endocytosis is blocked, HbVs are not formed, indicating these vesicles or their content originate from the cytostome (Sabitzki et al., 2024). Transmission electron microscopy indicated that at least some of the HbVs were double-membraned (Sabitzki et al., 2024), in line with their cytostomal origin which is an invagination of two membranes (Spielmann et al., 2020). Furthermore HbVs can harbor intraluminal vesicles (Jonscher et al., 2019; Sabitzki et al., 2024; Schmitz et al., 2026), potentially indicating similarities to ESCRT generated intraluminal vesicles in model organisms. However, apart from their Hb content, little is known about their protein composition and characteristics. PI(3)P and Rab5b were detected at HbVs (Jonscher et al., 2019; Sabitzki et al., 2024) but as PI(3)P typically is found at the DV of the parasite (Tawk et al., 2010), it is not clear whether this indicates an early endosomal characteristic. Overall, it is unclear whether HbVs are endosomes (i.e., sorting stations such as EEs in other organisms), or are pure transport vesicles, and their position in the endosomal pathway and the characteristics of that pathway in the parasite are not well understood.

Here, studied HbV morphology and surface protein composition using cryo-electron tomography (cryo-ET) and BioID. Our findings provide high resolution characterization of HbVs, identify cargo of HbVs and identifies PfTBC8 as a Rab effector with an important function in endosomal trafficking in *P. falciparum* blood-stage parasites.

## RESULTS

### HbVs are double membrane, coatless vesicles containing diverse intraluminal structures

To better understand the morphology of these hemoglobin-filled vesicles, we used in-situ cryo-electron tomography (cryoET) to image the parasites in a fully hydrated, vitrified state, yielding artifact-free, cryo-preserved images of the accumulated HbVs within parasite-infected RBCs at high resolution in 3D (Fig. 1A, Fig. S1A, Table 1). Analysis of 126 individual HbVs across 101 cryoET tomograms and 61 parasite-infected RBCs revealed that these HbVs are double-membraned, with a mean maximum diameter of 618.88 nm (median = 630.76 nm, SD = 166.58 nm) (Fig. 1D). This double-membrane architecture together with the density of their content confirms their identity as vesicles of endocytic origin. Surprisingly, we observed no dense protein coat on the outer surface of the vesicles (Fig. 1). We observed structural variations along the bounding membranes within individual HbVs: 19.9% of the quantified HbVs contained regions where the inner lipid bilayer was partially degraded or fragmented (Fig. 1A, black arrow), leaving a single, intact outer membrane (Fig. 1A, pink arrow). Additionally, 27% of the HbVs contained intraluminal bodies within the vesicular lumen. These intraluminal bodies exhibited diverse shapes, including snowman- (Fig. 1E) and horseshoe-shaped (Fig. 1F) architectures, separate hemoglobin-filled compartments trapped within the intermembrane space (Fig. 1G), and intraluminal bodies enclosed by one or more membrane layers within the vesicular lumen (Fig. 1H). The latter was the most frequent (50% of all HbVs with intraluminal bodies), whereas the snowman shape was the rarest (2.9% of HbVs with intraluminal bodies) (Fig. 1I). Finally, we observed that 4.8% of the HbVs exhibited both boundary membrane heterogeneity and the presence of these intraluminal bodies (Fig. 1B). Intraluminal bodies were found in similar proportion within vesicles with and without single-membrane regions. To determine whether these distinct morphologies correspond to differences in vesicle size, we compared the HbVs diameters of the four groups (HbVs, HbVs with single-membrane only, HbVs with intraluminal body only, and HbVs with both features). A one-way ANOVA revealed no overall statistically significant difference in vesicle diameter across the subpopulations (p = 0.474) (Fig. S1B).

**Figure 1.**
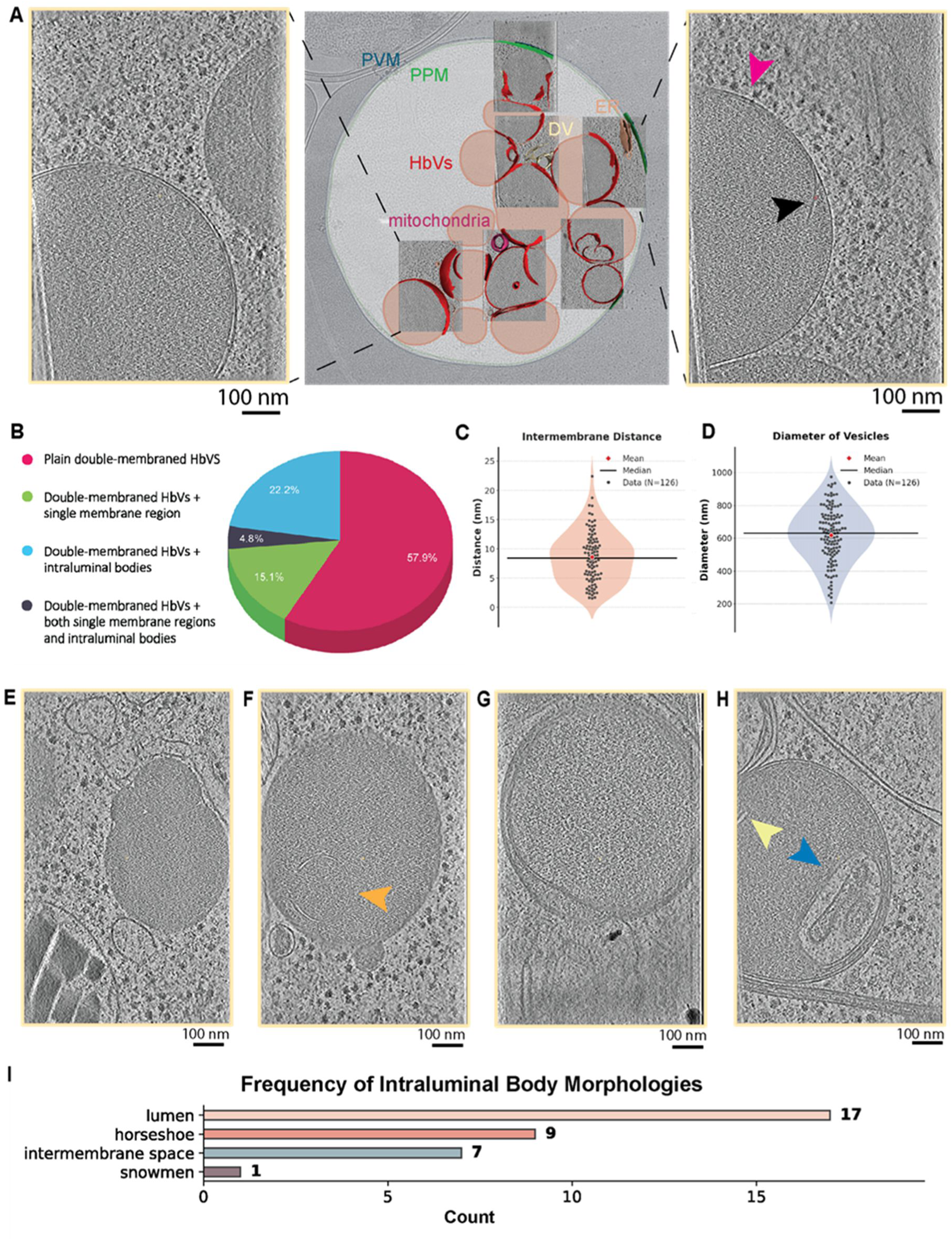
Structural characterization of HbVs. **A)** Averaged central slice from a tomogram (see also Fig. S1A) showing HbVs membranes (left panel). Averaged central slice from a tomogram showing HbVs morphological heterogeneity along the vesicle membranes (right panel). Arrowheads denote regions exhibiting standard double-membrane architecture (pink arrowhead) versus localized single-membrane morphology (black arrowhead). **B)** Proportion of HbVs structural features. **C)** Distribution of mean intermembrane distances per tomogram across the PfRbsn5L inactivation accumulated HbVs along double-membraned regions (y-axis: Distance in nm), with an average spacing of 8.44 nm (SD = 4.18 nm). **D)** Distribution of HbVs diameter (y-axis: Diameter in nm), with an average diameter of 618.88 nm (SD = 166.58 nm) **E-H)** Averaged central slice from a tomogram showing snowman-shaped compartmentalized HbVs (E), horseshoe shaped ILB (orange arrowhead) inside HbVs (F), intraluminal bodies (ILB) in HbVs intermembrane space (G), ILBs enclosed by one (yellow arrowhead) or more membrane layers (blue arrowhead) within the vesicular lumen (H). **I)** Histogram detailing the frequency of specific intraluminal body morphologies. Corresponding reference structures are depicted in (E-H).

**Table 1.**
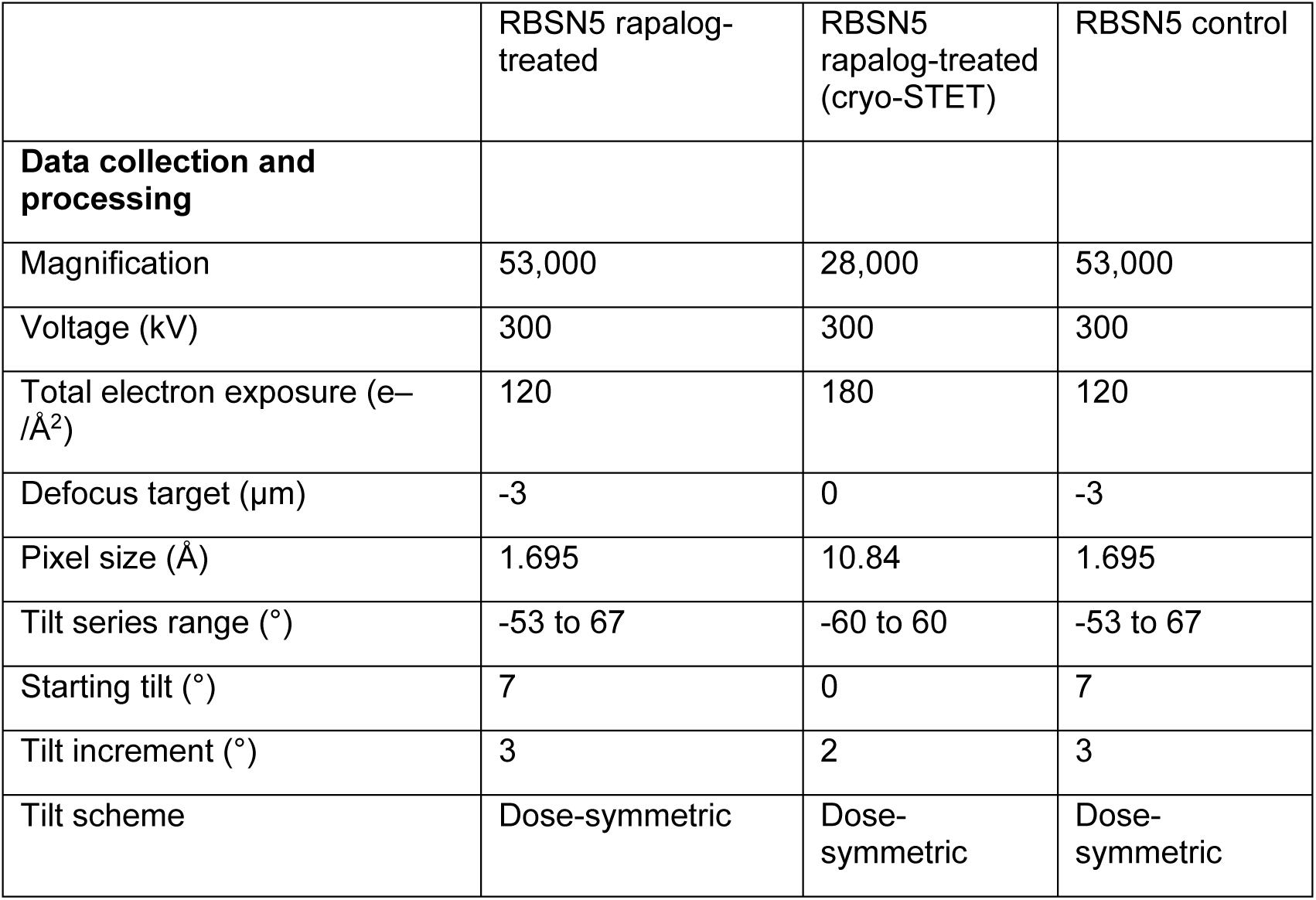
Cryo-ET data collection.

Next, we quantified the spacing between the two lipid bilayers in double-membraned regions, finding an average intermembrane distance of 8.44 nm (median = 8.33 nm, SD = 4.18 nm) (Fig. 1C, Fig. S2). This spacing remained consistent regardless of vesicle morphology; a one-way ANOVA revealed no statistically significant differences in intermembrane distance across the four HbV subpopulations (p = 0.262) (Fig. S1B). Because these bounding membranes originate from the PVM and PPM, the intermembrane space of the HbV derives directly from the parasitophorous vacuole, which is known to be abundantly populated with membrane-protein complexes and soluble proteins, including the PTEX translocon, the EPIC complex, and early transcribed membrane proteins (ETRAMPs) (Batinovic et al., 2017; Ho et al., 2018; Spielmann et al., 2012). As such, we were surprised to observe that the intermembrane space in the HbVs appeared largely devoid of such protein densities. This loss of intermembrane protein density suggests an exclusion event occurs during or after vesicle scission. Finally, we also observed 3 instances indicating fusion of HbVs with the DV (Fig. S1C).

### Surface proteome of HbVs using a PI(3)P sensor BioID detects endosomal poteins

Two thirds of HbVs are positive for PI(3)P at all tested times after their induction (Jonscher et al., 2019; Sabitzki et al., 2024) which can be detected with the probe Px-p40 (Kanai et al., 2001) while in the unperturbed state PI(3)P is predominantly found at the DV (Tawk et al., 2010). We took advantage of this to carry out a BioID (Roux et al., 2012) enriching for proteins on the HbV surface by fusing miniTurbo with Px-p40 (Fig. 2A) and then inducing HbVs either by inactivating RBSN5L or by adding CytD and comparing it to the control (Fig. 2B). The presence of Px-p40-miniTurbo at the induced HbVs and at the DV in controls was confirmed by microscopy (Fig. 2C). Quantitative BioID in trophozoites (Fig. 2D), detecting proteins enriched in the HbV condition over control, identified plasmepsins, DV proteases suspected to be transported indirectly via the PV followed by endocytic internalization (Klemba et al., 2004) as the top enriched proteins shared in both conditions (Fig. 2E,F, Fig. S3 and Table S1). A more mildly, but still significantly enriched group of proteins included two TBC domain proteins, Rab11a and Rab11b, the CORVET subunit VPS3, the retromer-complex component VPS35 and the endosomal transport and maturation organization protein FHIP1b (Fig. 2E,F; Fig. S3 and Table S1). While enriched proteins were generally shared in both conditions (Fig. 2F), the group of mildly enriched proteins that reached significance differed in the two conditions. In the RBSN5L condition Rab11a and a TBC domain protein were above the significance cut off while in the CytD condition this was the case for Rab11b, a different TBC domain protein and MyoF (previously linked to endocytosis (Schmidt et al., 2023)). Overall, these hits fit with an endosomal nature of the HbVs and suggest some degree of conservation to endosomal transport in better studied model organisms.

**Figure 2.**
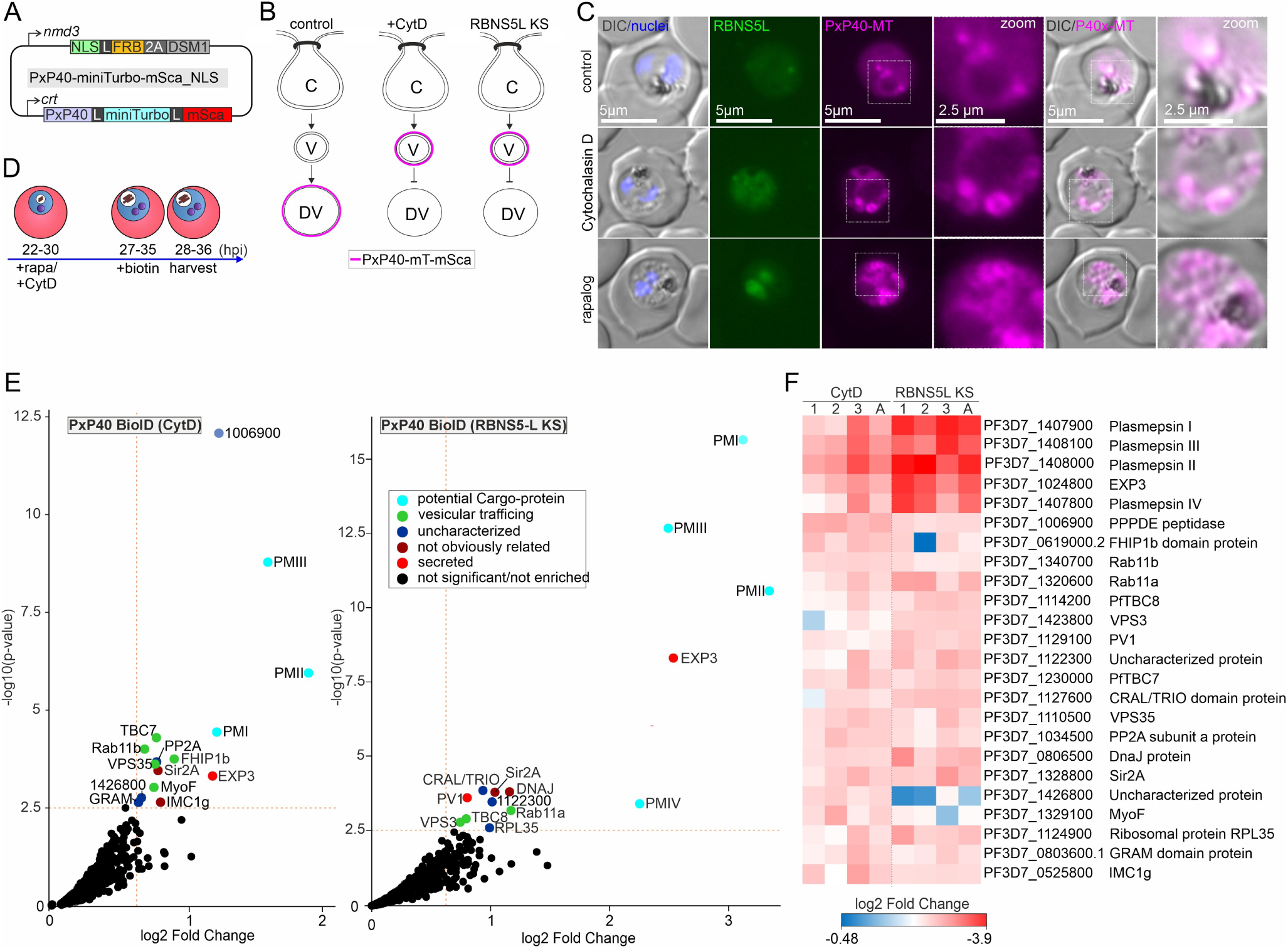
PI(3)P-based HbV surface proxiome. **A)** Schematic of the PxP40-miniTurbo-mSca_NLS plasmid used to generate the surface proteome of PI(3)P positive HCC-filled vesicles in the RBNS5L^endo^ parasites. **B)** Experimental setup showing the 3 BioID conditions (RBNS5L KS, Rabenosyn5-like KS). **C)** Representative live-cell fluorescence microscopy images (three independent experiments) showing Px-P40x signal as a probe for PI(3)P at the digestive vacuole (DV, control) or at induced vesicles induced as indicated (CytD or rapalog). Boxes, zoomed areas shown to the right (zoom). **D)** Scheme showing timing of experiment. **E)** Top-right quadrants of volcano plots for CytD and RBNS5-KS-induced vesicles (cut offs (dashed red lines): log2-normalised ratios rapalog over control ≥0.6, p-values ≤ 0.003 (-log10 ≥2.5) of three independent experiments). Significantly enriched proteins color-coded as indicated. Proteins designated by their annotated abbreviation if they have one; numbers indicate the respective PF3D7 PlasmoDB ID (see F for accessions). Full volcano plot shown in Figure S2. KS, knock sideways. **F)** Heatmap of significantly enriched proteins identified in at least one of the HbV-induction conditions in the PX-p40 BioIDs (log₂ fold enrichment) of experiments in (E).

### Halo-based pulse chase tracking shows that Plasmepsin2 is a bona fide cargo of the HbV pathway

The identification of plasmepsins and previous data (Klemba et al., 2004) supports the idea that these proteases are transported to the DV via endocytic re-internalisation. However, so far this conclusion is primarily based on immune-electron microscopy (EM) detection of PM2 at the cytostome (Klemba et al., 2004). Plasmepsins would be the first intramembrane cargo of the HbV pathway and the first non-host cell protein endocytosis cargo in the parasite. We therefore decided to assess the transport of PM2 in detail.

To analyze whether PM2 is indeed first transported to the PV and then internalized via the cytostome, we episomally expressed C-terminally Halo-tagged PM2 (Fig. 3A) in a cell line wherein we can conditionally prevent endocytosis at the cytostome by knock sideways (KS) of KIC7 (Birnbaum et al., 2020). The Halo tag permits time resolved tracking by labeling PM2-Halo populations with different fluorophore-conjugated Halo substrates (Koreny et al., 2023; Svendsen et al., 2008; von Knoerzer-Suckow et al., 2025). For tracking, we first stained (and thereby saturated off) previously expressed PM2-Halo with Halo substrate 1. After wash out of substrate 1, endocytosis was inhibited (KIC7 KS, induced by adding rapalog) followed by addition of Halo substrate 2 to track only PM2 produced after cytostome transport was blocked (Fig. 3B). Using this regimen, substrate 1-labelled PM2-Halo (the “old” PM2 pool) was almost exclusively found in the DV, consistent with this being the endpoint of PM2 trafficking. In contrast, the newly labelled PM2-Halo was detectable in the cell periphery in ∼20% of control and ∼80% of cells with inactivated cytostome function (Fig. 3C,D). While this was not formally tested here, the signal in the cell periphery likely constitutes PM2 embedded in the PPM, as processing to release the protease domain from the membrane-embedded pro-peptide part occurs only under acidic conditions (Francis et al., 1997). The detection of cell peripheral Halo2 signal in some of the control cells indicated that already the time resolved Halo labelling was sufficient to visualize PM2 transiently present at the PPM, but after blocking cytostome function, the accumulation in the PPM was substantially increased (Fig. 3D). These results indicate that PM2 is indeed trafficked via secretion to the PPM and re-internalization by the cytostome.

**Figure 3.**
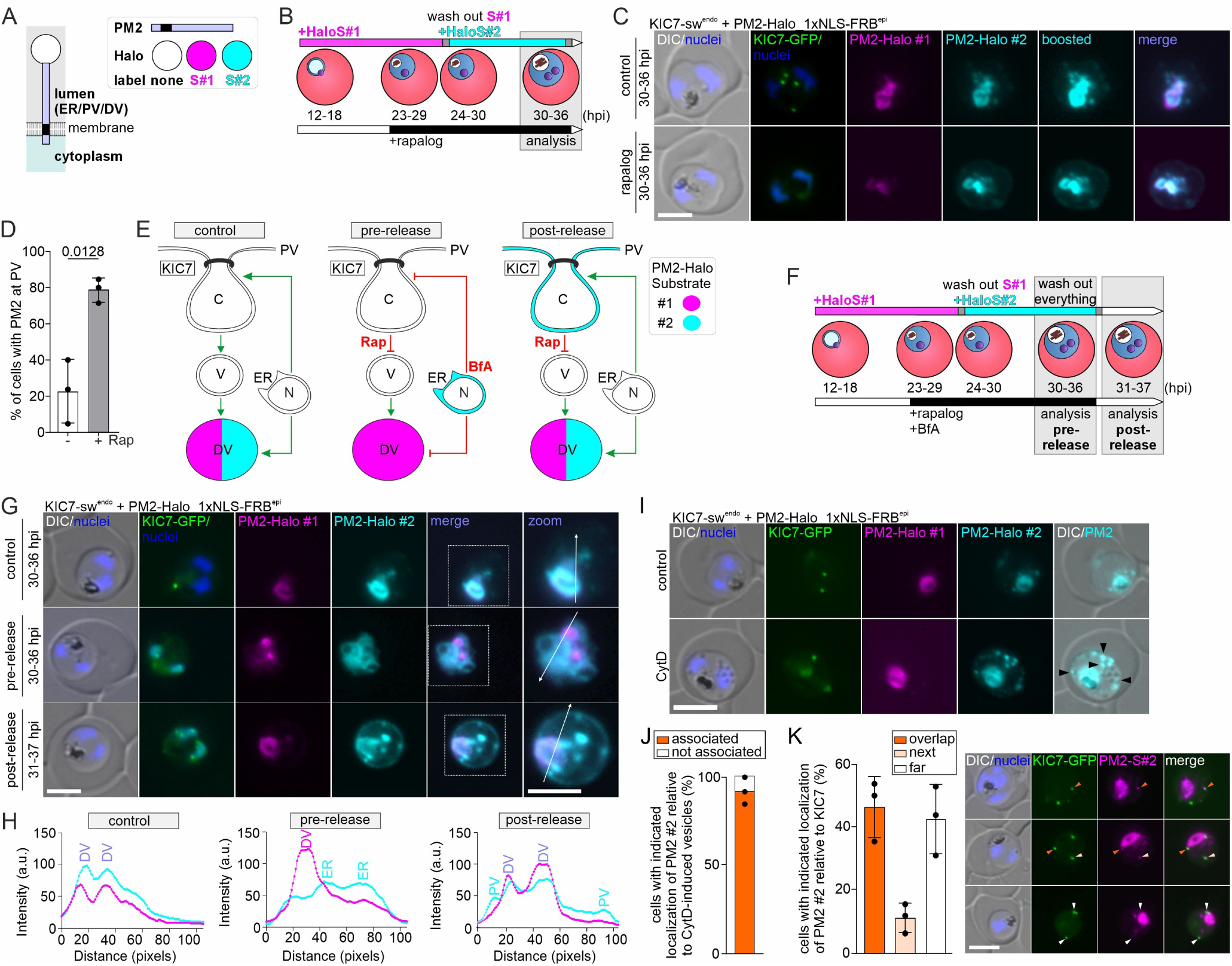
Plasmepsin Trafficking. **A)** Scheme showing PM2 topology and Halo labels. **B)** Experimental regimen for Halo substrate and rapalog addition used for (C). **C)** Live cell microscopy images of control and KS (rapalog) KIC7-sw^endo^ parasites episomally expressing PM2-Halo_1xNLS-FRB at 30-36 hpi following the experimental setup shown in (B). **D)** Quantification of percent of control (-Rap) and KIC7-KS (+Rap) parasites showing PM2 signal at the PV at 30-36 hpi following the experimental setup shown in (B). Black dots represent mean of one of three independent experiments with 21, 15, 21 (-Rap) and 20, 13, 14 (+Rap) parasites analyzed. Bars show mean and SD. P value, paired t-test of the means. **E, F)** Expected outcomes (E) and experimental regimen (F) for (G); C, cytostome; V, HbV; N, nucleus; ER, endoplasmatic reticulum; BfA, brefeldin A; Rap, rapalog, DV, digestive vacuole. Green arrows active pathways, red lines inhibited pathways. **G)** Live cell microscopy images of KIC7-swendo+PM2-Halo_1xNLS-FRBepi parasites at the indicated time points following the experimental setup shown in (F). Arrows indicate positions and directions of the intensity measurements corresponding to the intensity plots shown in (H). **H)** Exemplary intensity plots of the PMII signal (arbitrary units, a.u.) in the cells shown in (G). **I)** Live cell microscopy images of control and CytD-treated KIC7-sw^endo^+PM2-Halo_1xNLS-FRB^epi^ parasites following the experimental setup shown in (B): At 23-29 hpi, CytD was applied and cells were analyzed at 30-36 hpi. Arrowheads indicate association of PM2 with CytD-induced vesicles. **J)** Quantification of cells with indicated localization of PM2 #2 relative to CytD-induced vesicles in (I). **K)** Quantification of cells with indicated localization of PM2 #2 relative to KIC7 in KIC7-sw^endo^+PM2-Halo_1xNLS-FRB^epi^ control parasites. Parasites were assigned to the categories ‘overlap’, ‘next’, and ‘far’ if at least one PM2 focus per cell exhibited the respective localization pattern. Analysis based on three independent imaging sessions with 17, 14, 18 parasites analyzed. Bars show mean of the means per experiment and SD thereof. Nuclei were stained with Hoechst. Scale bars, 5µm. DIC, differential interference contrast; merge, merged channels of magenta and cyan or GFP and magenta (K).

However, there was still Halo substrate 2-labelled PM2 detectable in the DV in these experiments (Fig. 3C). In order to test whether this could be PM2 transported by an endocytosis-independent route, we reduced the potential leakiness of the Halo labelling assay by arresting newly synthesized and substrate 2-labelled PM2 in the secretory pathway through addition of BrefeldinA (Misumi et al., 1986) and at the same time blocking cytostome function (Fig. 3E (expected outcomes) and Fig. 3F (experimental scheme)). No newly labelled PM2 was detected in the DV before BrefeldinA was lifted, indicating efficient arrest of secretory trafficking (Fig. 3G,H, pre-release). Release of the BrefeldinA block resulted in appearance of substrate 2-lablled PM2 at the PPM and the DV (Fig. 3G,H, post-release). The PPM pool of PM2 confirmed the PPM-cytostome-endosomal transport route while the DV pool indicated either that the cytostome block was not absolute or that some of the PM2 also uses a direct Golgi to DV route. In conclusion, PM2 was confirmed to be a bona fide cargo of the endosomal transport pathway but there may also be a parallel direct Golgi to DV route.

Next, to test whether after passing the cytostome, PM2 is transported by HbVs, as suspected from our BioIDs, we again first saturated off all previously expressed PM2-Halo in the cells using Halo substrate 1, washed out substrate 1 and then labelled all newly expressed PM2-Halo with substrate 2 after addition of CytD to induce HbVs. While controls showed Halo1 and 2 signal predominantly in the DV, the cells grown in the presence of CytD showed Halo2-labelled (“newly made”) PM2 in foci associated with the induced HbVs in addition to the DV while the Halo1-labeled (“old”) PM2 was mostly found in the DV (Fig. 3I,J). These results indicate that PM2 is present at the HbVs, although not universally distributed around them (Fig. 3I). In a substantial proportion of control parasites, substrate 2-labelled PM2-Halo was also detected in foci fully or partially overlapping with the cytostome (as marked by KIC7) (Fig. 3J), indicating some accumulation at the cytostome before internalization.

Taken together, these data indicate that PM2 is transported from the ER to the PPM where it is internalized by the cytostome and transported by HbVs to the DV. Consistent with detection at cytostomal structures by EM (Klemba et al., 2004), we detected regular localisation close or overlapping with the cytostome in live control cells when tracking freshly produced PM2, and blocking cytostome function led to accumulation in the PPM. However, assuming our assay is not leaky, there may also be a direct ER to DV route of PM2 transport. These findings show that PM2 is a bona fide cargo of endocytosis that can be used to track this transport pathway. These findings also validate the identification of PM2 in the HbV surface BioID and by extension the other plasmepsins which likely are also cargos.

### PfTBC8 is needed for blood stage growth

Beside the cargo proteins enriched in the HbV BioIDs, a number of the hits had already experimental links to endocytosis in the parasite (VPS3 which is part of CORVET) (Mesén-Ramírez et al., 2025) or endosomal functions in model organisms (the Retromer subunit VPS35, FHIP1b/Hook) (Seaman, 2012; Xu et al., 2008). To find potentially new mechanistic endocytosis proteins, we selected one of the two TBC domain proteins (PF3D7_1114200, the one more enriched in the RBNS5L KS condition) for further analysis. TBC domain proteins typically act as GAPs for Rabs and are key regulators of vesicular trafficking (Homma et al., 2021). Sequence and AlphaFold (Abramson et al., 2024) structure comparisons confirmed presence of the TBC domain, which encompassed most of the PF3D7_1114200 sequence (Fig. 4A). A recent systematic analysis of TBC domain proteins in *T. gondii* listed PF3D7_1114200 as the malaria equivalent of TgTBC8. A comparison of the TBC domain proteins of different Apicomplexans supported this classification (Fig. S4). Consequently, we here refer to this protein as PfTBC8.

**Figure 4.**
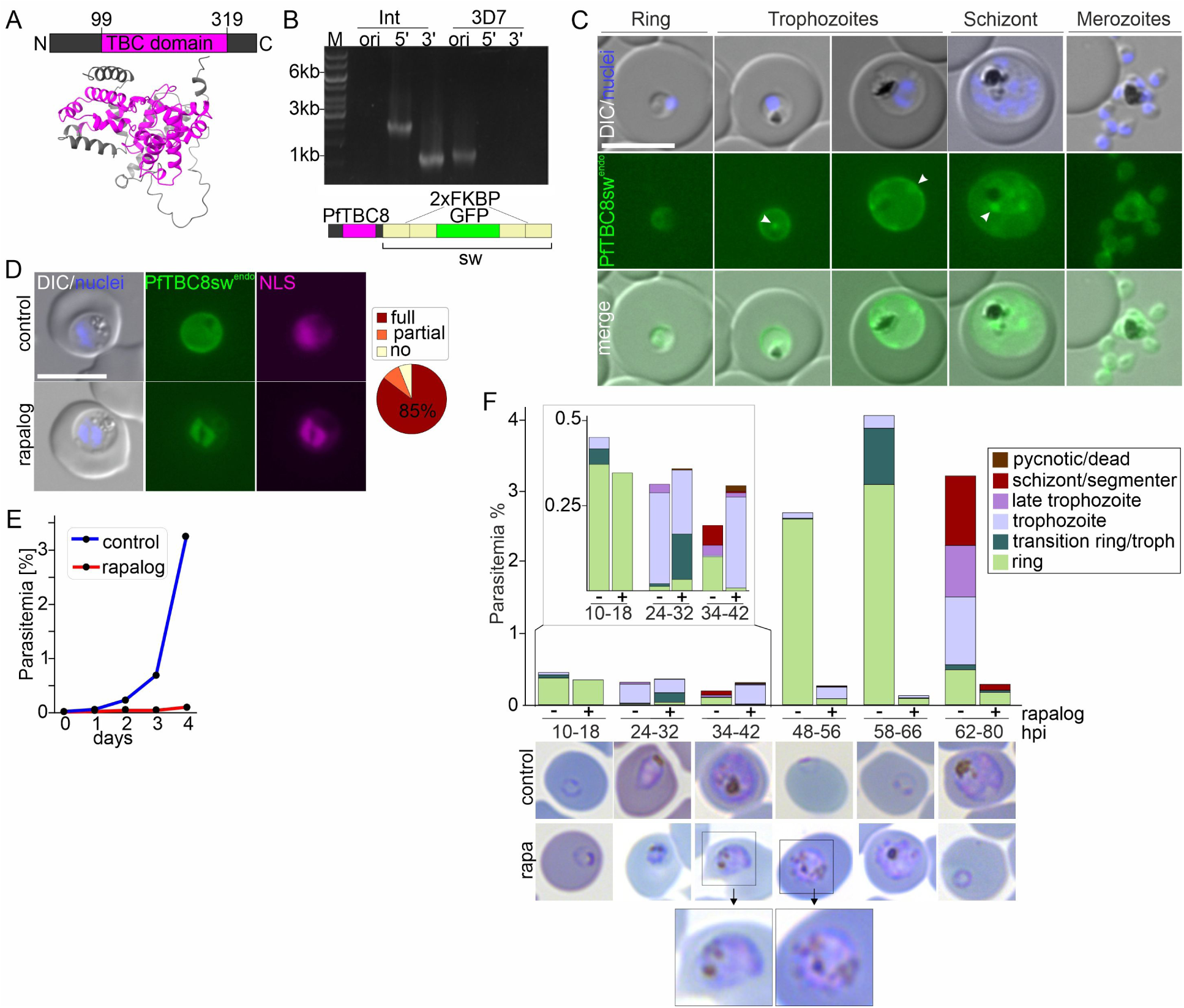
PfTBC8 is needed for blood stage growth. **A)** Domain organization and Alphafold2 predicted structure of PfTBC8. **B)** Agarose gels of PCR products from gDNA showing correct modification of the *P. falciparum* genome using primers across the 5’- (5’ int) 3’ - (3’ int) integration junction and absence of original locus (ori) in the integration parasites (Int) or 3D7 parental parasites (3D7) to obtain sw tagged PfTBC8 (see scheme below) expressed from the endogenous locus (PfTBC8-sw^endo^ parasites). M, marker (in kb). **C)** Representative live-cell fluorescence microscopy images of PfTBC8-sw^endo^ parasites at the indicated parasite stages (3 independent imaging sessions with at least 10 cells imaged). **D)** Representative live-cell fluorescence microscopy images of PfTBC8-sw^endo^ parasites expressing the 1xNLS mislocaliser (1xNLS-FRB-mCherry) for KS with rapalog and without (control) rapalog for one hour. Pie shows KS efficiency quantified as indicated (four independent experiments (rapalog: 14, 21, 25, and 22 cells, control 11, 12, 18, 12 cells). E) One of three representative growth curves (replicates in Fig. S5A) of PfTBC8-sw^endo^ KS parasites grown with rapalog (KS induction) or without (control) over 5 days (parasitemia measured by flow cytometry). **F)** Giemsa smears and quantification of stage progression and growth of PfTBC8-sw^endo^ KS (rapalog) compared to controls in synchronous parasites (KS induced in ring stages). Representative Giemsa smear images are shown. Black boxes, enlarged area shown below. For stage quantification at least 60 parasites or >15000 red blood cells per timepoint and condition were scored with the exception of the rapalog condition at 62-80 and 72-90 hpi because only very few parasites were left (where scoring stopped after 3000 RBCs). Two independent experiments (replicate in Fig. S5B). rapa, rapalog; hpi; hours post invasion.

In order to localize and functionally analyse PfTBC8, we used SLI to fuse it with the multipurpose sandwich (sw) tag (2xFKBP-GFP-2xFKBP) through modification of the genomic gene locus (Birnbaum et al., 2017) (Fig. 4B). Live cell microscopy with the resulting cell line (PfTBC8-sw^endo^) showed a GFP signal at the parasite plasma membrane with occasional circular or focal structures that often were also visible by DIC (Fig. 4C). No indication was found for a location at the IMC, the localization of TgTBC8 (Quan et al., 2024). However, while an IMC localization provides a typical pattern during schizont development, we can’t exclude the protein moves to such a localization in very late stage parasites where an IMC-localization resembles a PPM localization.

For functional analyses, the PfTBC8-sw^endo^ cell line was transfected with a plasmid mediating episomal expression of the 1xNLS mislocaliser (Birnbaum et al., 2017) to enable conditional removal of PfTBC8 to the nucleus by KS through addition of rapalog. KS efficiently mislocalised PfTBC8 to the nucleus 1 h after induction (Fig. 4D) and caused a severe growth defect compared to controls when parasite cultures were tracked over two developmental cycles (Fig. 4E, Fig. S5A). Induction of the KS in synchronous parasites and tracking of progression of parasites through blood stage development using Giemsa stained smears showed that the growth defect manifested in the trophozoite stage: while controls progressed through the cycle, the PfTBC8 KS parasites remained trophozoites and failed to reach the late trophozoite stage (Fig. 4F, Fig. S5B). Inspection of the Giemsa smears indicated that the stalled trophozoites frequently contained a comparably small amount of hemozoin in the DV and additional hemozoin crystals dispersed in the cytoplasm. Overall these findings show that PfTBC8 is a PPM protein important for the survival of asexual blood stage parasites with a function in trophozoites.

### PfTBC8 controls Rab11a

To gain more insight into the function of PfTBC8, we carried out DiQ-BioID (Birnbaum et al., 2020) to identify the proteins in its vicinity. For DiQ-BioID the biotinylising enzyme (here miniTurbo) is recruited to the target protein (PfTBC8) by conditional dimerisation using the FKBP-FRB system, taking advantage of the multipurpose sw tag on TBC8 (Fig. 4B). The DiQ-BioID approach permits removal of background biotinylation by quantitative mass spectrometry comparing biotinylated proteins when miniTurbo is dimerized to PfTBC8 (induced with rapalog) over control when it is not.

Consistent with the location of PfTBC8 at the PPM, many of the 49 significant DiQ-BioID hits (log2 fold enrichment >0.5, p>0,00316 (-log10 > 2,5)) were known or suspected PPM proteins (Fig. 5A, Table S2) such as e.g. PfFNT (Marchetti et al., 2015; Wu et al., 2015), PfNCR1 (Zhang et al., 2024) or ATP4 (Dyer et al., 1996; Rottmann et al., 2010). Very few typical vesicle trafficking proteins were in the hit list (Fig. 5A). This included Rab11a, the only enriched Rab which interestingly was also the best enriched Rab in the RBSN5 KS induced HbV BioID that had originally identified PfTBC8 (Fig. 2E,F). Given the expected role of TBC domain proteins in regulating Rabs, Rab11a may be the cognate Rab target of PfTBC8. To more directly test this, we assessed whether Rab11a responds to PfTBC8 inactivation by expressing it in the PfTBC8 KS parasite line. Under control conditions, episomally expressed mScarlet-Rab11a was detected around the nucleus, indicating an ER localization (Fig. 5B). However, when we induced the PfTBC8 KS, ∼60% of the cells additionally showed a signal of Rab11a at the PPM (Fig. 5B), the site where PfTBC8 normally would be located. In contrast, neither Rab6, a trans Golgi Rab also found at the ER (Cubillán-Marín et al., 2025), nor Rab5a, a Rab not involved in endocytosis (Farias et al., 2026; Sabitzki et al., 2024), were detected at the PPM after PfTBC8 KS and did not visibly alter their localization (Fig. 5C, Fig. S6). These findings indicate that inactivation of PfTBC8 leads to an accumulation of Rab11a at the PPM, providing evidence that it is the GAP inactivating Rab11a and failure to do so leaves Rab11a at the site where it carries out its function.

**Figure 5.**
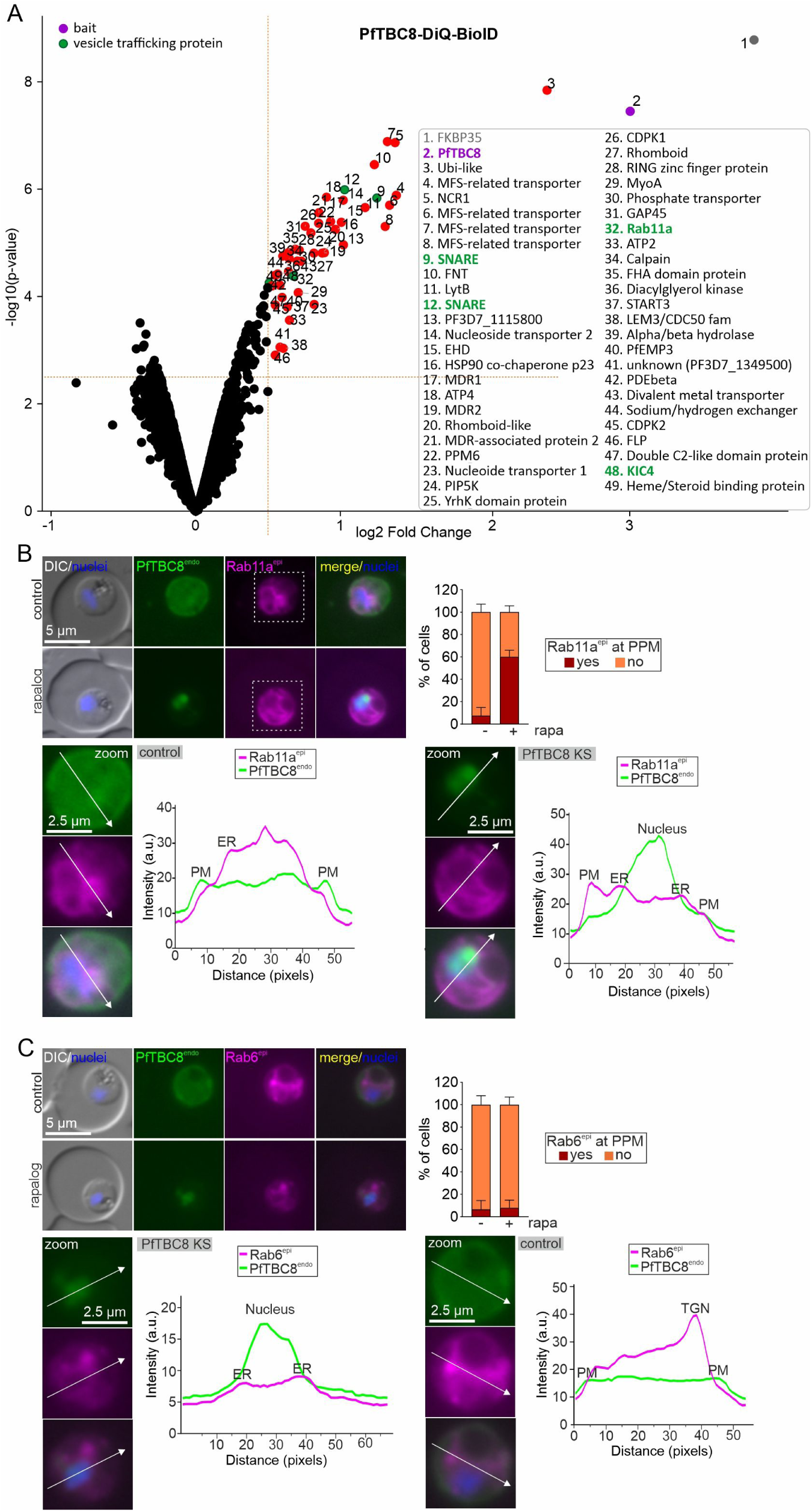
PfTBC8 proxiome contains Rab11a which is affected in localization after PfTBC8 KS. **A)** DiQ-BioID with endogenously tagged PfTBC8-sw (PfTBC8-sw^endo^) episomally co-expressing FRB-miniTurbo (cut offs (dashed red lines): log2-normalised ratios rapalog over control ≥0.5, p-values ≤ 0.003 (-log10 ≥2.5) of three independent experiments). **B-C)**, Representative live-cell fluorescence microscopy images of PfTBC8-sw^endo^ (green channel) episomally co-expressing Rab11a (B) (magenta channel) or Rab6 (C) upon PfTBC8-sw KS (rapalog) and under control conditions (control). Zoom shows enlarged area (last three images) of respective GFP and magenta channel or merge/nuclei image. White arrows indicate line used to calculate the intensity values shown in. Example intensity plots of the PfTBC8-sw^endo^ signal (green) and the Rab11a and Rab6i signal (magenta) (arbitrary units, a.u.) are shown. Bar graphs show quantification of number of cells (%) showing plasma membrane-associated Rab11a (B) or Rab6 (C) signal with rapalog (PfTBC8 KS) or not (control) (n = 3 independent replicates with 12, 12, 9 cells (rapalog) and 12, 12, 7 cells (control) in the Rab11a co-expressing parasites and 19, 25 and 25 cells (rapalog) and 13, 24 and 24 cells (control) in the Rab6 co-expressing parasites. Nuclei were stained with Hoechst, merge: overlay of green and magenta channel. Scalebar indicated. DIC, differential interference contrast; PM, Plasma membrane; ER, Endoplasmic reticulum; TGN; trans-Golgi.

### Rab11a is at HbVs

Given the connection of Rab11a and PfTBC8, and the enrichment of both of these proteins in the HbV BioIDs, we tested whether these proteins are located at HbVs. While PfTBC8 had been detected as a significant hit only in the RBSN5L KS-induced HbV BioID, we so far had failed to obtain a RBSN5L KS parasite line to assess the location of PfTBC8 under this HbV-induction condition. We therefore resorted to using CytD to induce HbVs. While we did observe cells with PfTBC8 surrounding vesicles obtained with that method (Fig. 6A), when we compared this to controls, the number of circular PfTBC8 structures was decreased (Fig. 6A). We conclude that there is no clear association of PfTBC8 with CytD-induced HbVs although we do not know whether the structures in controls are the same as those seen when HbVs are induced. However, even if different and bona fide HbVs, the proportion of cells with PfTBC8 at these vesicles was low (Fig. 6A).

**Figure 6.**
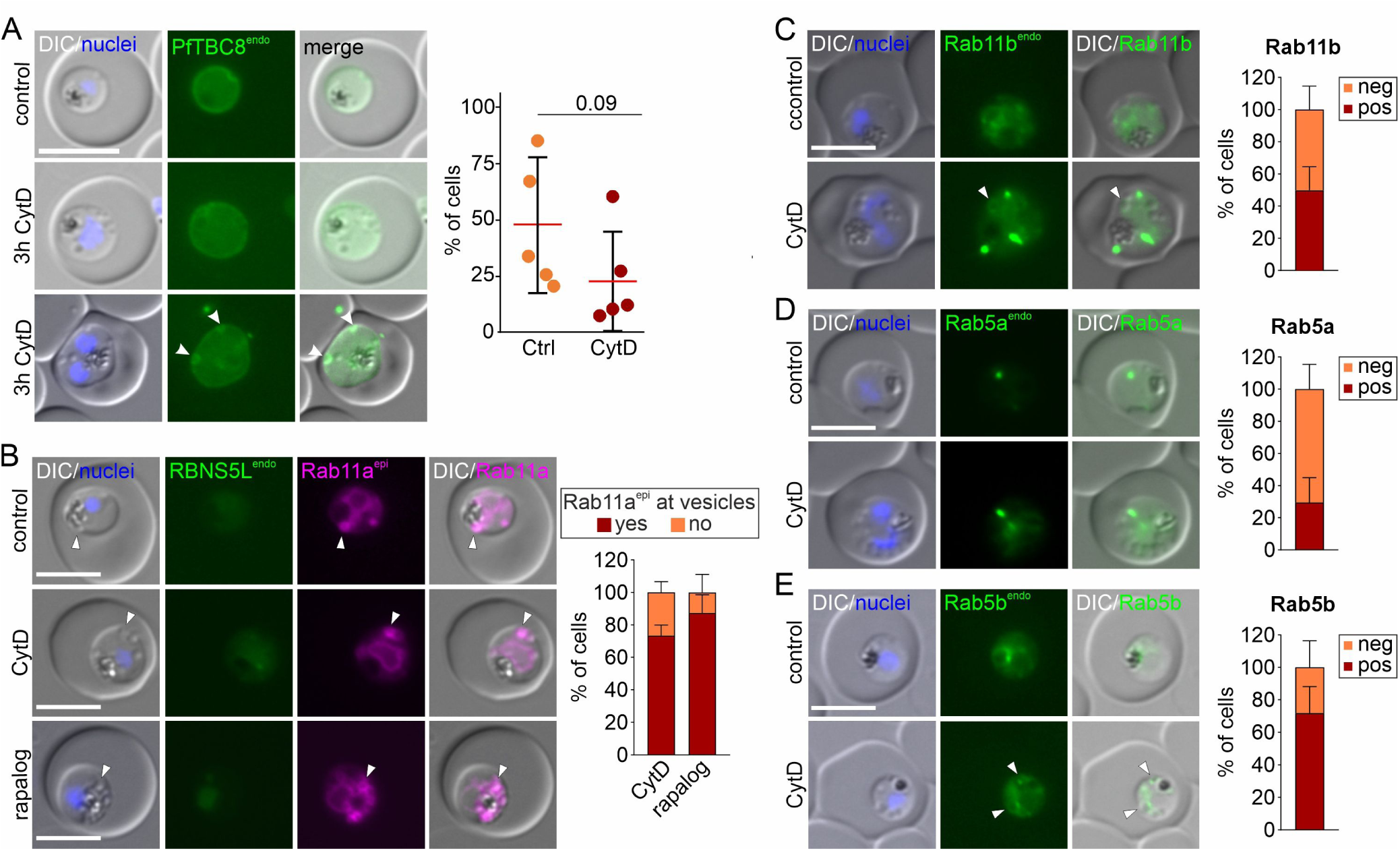
Testing association of PfTBC8 and selected Rabs with HbVs. **A)** Representative live-cell fluorescence microscopy images of PfTBC8-sw^endo^ parasites with CytD (3h CytD; two images, one showing a parasite with accumulation at vesicles and one without) and controls. Graph shows quantification of cells (%) with PfTBC8 signal at vesicles upon CytD addition or in controls (Ctrl). Bars show SD (black lines) and average (red line) (n=5 independent replicates (control: 3, 7, 4, 4 and 5 cells; CytD: 10, 5, 12, 14, and 10 cells). Merge, overlay of DIC and GFP. **C-D)** Association of the indicated Rab proteins CytD or RBNSL5 KS induced vesicles. Left, representative live cell images, right quantification of phenotype of n=3 independent experiments. Rab5b (10,16 and 9 cells), Rab11b (9,15 and 9 cells); Rab5a (12,15,12 cells). DIC, differential interference contrast; Nuclei stained with Hoechst. Scale bar 5 μm.

Next, we analysed Rab11a for which we obtained a RBSN5L KS line. Induction of HbVs by inactivation of RBSN5L resulted in ∼90% of cells showing an accumulation of Rab11a surrounding the induced HbVs (Fig. 6B, rapalog) and this was the case for ∼70% of cells when CytD was used to induce HbVs (Fig. 6B, CytD). This validated the detection of Rab11a in the HbV BioID (Fig. 2E, F).

We also tested Rab11b, Rab5a and Rab5b using CytD to induce HbVs. Rab11b, which had been more enriched in the CytD-induced BioID (Fig. 2E, F), was also detected at CytD-induced HbVs, but often this was faint and less clear than with Rab11a (Fig. 6C). Rab5a, not involved in endocytosis, showed little accumulation at CytD vesicles (Fig. 6D), while Rab5b, which has a role in endosomal transport, was detected at HbVs (Fig. 6E), as reported previously (Sabitzki et al., 2024). Overall, these findings indicate that Rab11a has a function at HbVs and potentially also its GAP PfTBC8, although we could here not establish an association of the latter protein at HbVs using CytD.

### PfTBC8 is needed for efficient HCC delivery to the DV

To assess a potential role of PfTBC8 in endocytosis, we carried out bloated FV assay (Jonscher et al., 2019). In this assay digestion of hemoglobin is inhibited with E64 (Rosenthal et al., 1988) which leads to bloating of the DV if endocytosis takes place. After conditional inactivation of PfTBC8 function by KS, the number of cells with a bloated DV was reduced to ∼ 50% compared to close to 100% in controls (Fig. 7A), indicating that PfTBC8 is needed for efficient endocytosis.

**Figure 7.**
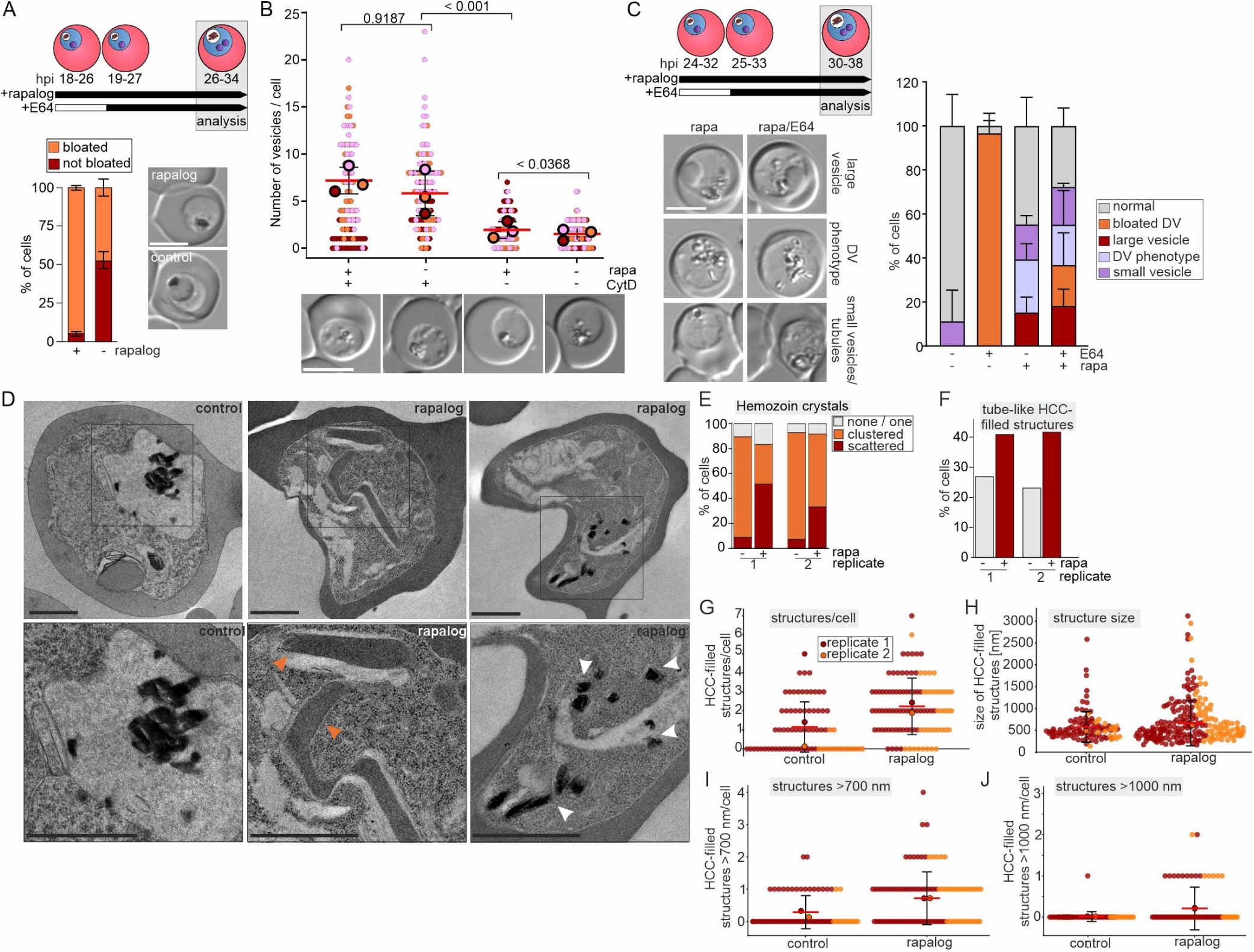
PfTBC8 is needed for efficient endocytosis. **A)** Bloated DV assay in PfTBC8 KS (rapalog) and control parasites grown in the presence of E64. Top shows experiment timing. Microscopy images show example DIC images. Graph shows number of cells with bloated DVs (n=3 independent experiments with 25, 23, 32 (control) and 30, 24, 30 (rapalog)). **B)** CytD endocytosis assay with PfTBC8 KS (rapalog) and control parasites. Superplot showing quantification of number of vesicles of before (-rapa) and after (+ rapa) KS induction and with or without CytD (+/- CytD) (n=3 independent experiments with - rapa/ - CytD: 31, 22, 27 cells; rapalog: 38, 35, 29 cells; CytD: 21, 30, 40 cells; rapalog and CytD: 40, 32, 34 cells. Small dots represent value from 1 parasite, large dots the mean of each independent experiment as per colour; two-tailed paired t test of the means, p-values indicated; mean (red); error bars (black) show SD. One representative DIC microscopy image for each condition is shown below the respective graph. **C)** Phenotypes scored in DIC Images in PfTBC8 KS and control parasites treated according to the indicated experimental scheme (n = 3 independent experiments with 21, 16, 11 (control), 39, 20, 10 (control with E64), 30, 17, 19 (rapalog) and 58, 35, and 19 cells (rapalog with E64); bars show SD). **D)** Representative electron microscopy images from two independent replicates of the PfTBC8 KS (rapalog) and control parasites. Black boxes, enlarged shown below. White arrows, scattered hemozoin crystals; orange arrows, elongated tube-like HCC-filled structures; Scale bars 500 nm. **E, F)** Bar chart showing percentage of cells with the indicated phenotypes. G -J) number per cell **G, I, J)** and size (diameter, H) of HCC-filled structures with the indicated specifications in PfTBC8 KS (rapalog) and control parasites based on the TEM analysis in (D). Replicates color-coded (n=2 independent experiments for control (57,14 cells) and rapalog (66, 36 cells)). Rapa, rapalog; HCC, Host cell cytosol; DV, digestive vacuole; hpi, hours post invasion. Rapa, rapalog; C, Cytostome; V, Vesicle, DV, digestive vacuole; ns, not significant. Scale bars; 5 μm.

So far, there were two endocytosis phenotypes described, an “early” phenotype stopping endocytosis before HbVs are formed (typically observed with cytostomal proteins (e.g. (Birnbaum et al., 2020)) or a “late” phenotype, which leads do the accumulation of HbVs (e.g. seen after CytD addition or inactivation of VPS45 or RBSN5L (Jonscher et al., 2019; Sabitzki et al., 2024; Smythe et al., 2007). As we did not detect the typical accumulation of vesicles in the cytoplasm after PfTBC8 KS (indicative of the late endocytosis phenotype), we tested whether PfTBC8 inactivation would reduce the number of CytD-induced HbVs (indicative of an early endocytosis phenotype as seen with KIC7 (Sabitzki et al., 2024)). However, PfTBC8 KS did not reduce the number of HbVs generated when CytD was added (Fig. 7B). Hence, despite the bloated DV assay phenotype, clearly showing reduced arrival of HCC in the DV, this could not be ascribed to the so far reported typical endocytosis phenotypes.

To obtain a better impression of the phenotype, we first assessed the appearance of the PfTBC8 KS cells in DIC. For this we focused on mid to late trophozoites (KS-induction at 24 - 32 hpi) where differences were more obvious than in younger stages. In contrast to controls, about 20% of the PfTBC8 KS cells contained a large vesicle like structure in the parasite cytoplasm, 20 – 40% of cells contained unfocussed or dispersed hemozoin crystals in the cytosol and about 10 – 20% of the cells contained small vesicle-like structures or tubules (Fig. 7C). The dispersed hemozoin crystals likely correspond to the dispersed hemozoin observed in the Giemsa stained parasites (Fig. 4E). We also included a condition where we added E64 to visualize whether the dispersed hemozoin arose from HCC digestion outside of the DV which would lead to bloating of extra DV regions. However, apart from the bloating of the DV observed in some cells, there was no apparent change in the frequency of the previously observed phenotypes which in total were detected roughly in 50% of the cells (Fig. 7C). Of note, with this later timing of the assay (24 – 32 hpi instead of 18 – 24 hpi for the bloated DV assay in Fig. 7A), even fewer cells showed a bloated DV after PfTBC8 KS, indicating the phenotype is more pronounced in later trophozoites and endocytosis more profoundly reduced (Fig. 7C).

To gain further insight into the PfTBC8 KS phenotype, we analysed the cells by transmission electron microscopy. The KS was induced in young trophozoites and the cells harvested in late trophozoites to cover the full phenotype. In contrast to controls, the PfTBC8 KS cells contained elongated hemoglobin filled tubular structures (Fig. 6D, orange arrows) and scattered hemozoin crystals (Fig. 6D, white arrows). Quantification of these phenotypes showed 30 to 50% of cells contained scattered hemozoin compared to less than 10% in controls (Fig. 6E) and that the number and size of structures filled with the density of HCC was more than in controls (Fig. 6F-J). We conclude that the internalized HCC accumulated in different structures and that likely the non-uniform nature of these structures prevented their detection in DIC in a similarly clear fashion to CytD or VPS45/RBSN5L/Rab5b-induced vesicles.

### Inactivation of PfTBC8 gradually impairs HbV intramembrane cargo protein trafficking

As HCC delivery was impaired after PfTBC8 KS but the phenotype had multiple manifestations of HCC accumulation in the parasite, we hypothesized that the phenotype could be due to a general deterioration of the endosomal transport pathway, rather than an interruption at a specific step. To test this, we assessed the functioning and condition of the endosomal transport pathway by analyzing cargo delivery of PM2-Halo after PfTBC8 KS. Again, we used the Halo tag to track newly synthesized PM2 (Fig. 8A). Halo substrate 1 was used to saturate PM2-Halo already present in the cell for 10 minutes, followed by adding rapalog to induce PfTBC8 KS. Halo substrate 2 was then added 5, 10, or 24 hours later to assess the fate of PM2-Halo synthesized after the KS was induced (Fig. 8A). In controls both Halo stained PM2 pools were detected in the DV (Fig. 8B-D). In contrast, in the PfTBC8 KS parasites the newly labelled PM2 (stained with Halo substrate 2) was detected outside of the DV in addition to some DV labelling (Fig. 8B-D). This phenotype was gradual: while 5 h after rapalog addition the new PM2 showed no significant difference in its cellular distribution compared to the control (Fig. 8B), increasingly more of the new pool of PM2 accumulated outside of the DV compared to control 10 h and 24 h after PfTBC8 KS (Fig. 8C-D), indicating a gradual impairment of PM2 delivery to the DV. This was not due to a loss of the DV, as the previously transported PM2-Halo (labelled with substrate 1) was still detectable in the areas overlapping with hemozoin (Fig. 8B-D). These findings are consistent with the idea that the endosomal trafficking route to the DV is gradually impaired in function when PfTBC8 is functionally inactivated.

**Figure 8.**
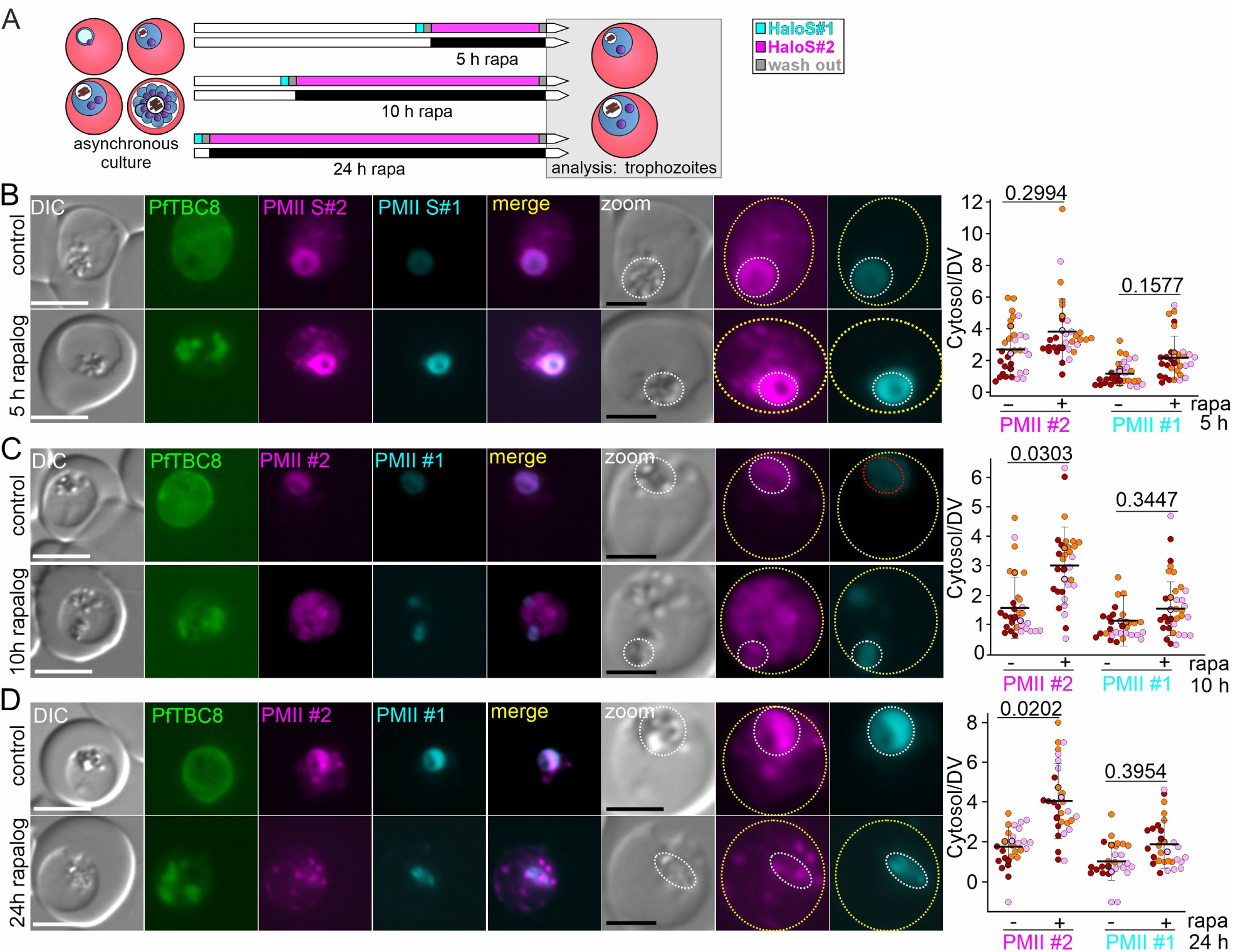
PMII trafficking after PfTBC8 KS is impaired. **A)** Schematic of the HaloTag-based pulse-chase labeling strategy used to monitor the trafficking of PMII upon PfTBC8 KS-induction. **B-D)** Representative live-cell fluorescence microscopy images showing PMII labeled with substrate#1 (PMII#1, cyan) and substrate#2 (PMII#2, magenta) upon PfTBC8 (green) KS-induction (rapalog) or under control conditions at 5 h (B), 10 h (C), and 24 h (D) with (rapalog) or without (control) rapalog. SuperPlots (Lord et al., 2020) show the Cytosol/DV ratio of the integrated density for each Halo substrate. Paired t-test of the mean, bars show SD and mean, p-value indicated (n=3 independent experiments (B: 10, 10, 10 cells for control and 10, 9 and 8 cells for rapalog; C: 10, 10, 10 cells for control and 10, 10 and 10 cells for rapalog; D: 7, 8, 10 cells for control and 9, 9 and 9 cells for rapalog). White scalebar: 5 μm, black scalebar 2.5 μm. Merge overlay of magenta (PMII#2, Halo2 substrate labelled) and cyan (PMII#1, Halo substrate 1 labelled) channel. Zoom shows enlarged area (last three images) of respective DIC, magenta or cyan channels. Yellow and red dashed line show parasite and DV outline, respectively. DIC, differential interference contrast; DV, digestive vacuole; RBC, red blood cell; rapa, rapalog

## Discussion

Endosomal transport in malaria parasites, particularly the pathway from the cytostome to the DV, is so far poorly defined (Spielmann et al., 2020). While it is now experimentally supported that the HbVs derive from the cytostome (Sabitzki et al., 2024), the nature of these structures is unclear. Based on size they could for instance be pinched off cytostomes, corresponding to oversized transport vesicles. However, based on precedents in better studied organisms, they also could correspond to endosomes which are more complex and versatile intermediary sorting stations that receive endocytic vesicles and mature while recycling processes remove components to be re-used and arrival of other components increase lytic capacity and reduce pH before they fuse with the lysosome (Huotari and Helenius, 2011).

Here we carried out a detailed morphological and surface proteome composition analysis of HbVs. Our CryoET analysis indicates that the accumulated HbVs all contain two membranes, but also showed that in about 20% of HbVs the inner membrane was ruptured. This might indicate a maturation process and upon fusion with the DV would lead to direct release of the HCC and broken membrane into the DV lumen. Previous work found that in V-type ATPase defective mutants intact HbVs accumulated in the DV which would support fusion of vesicles with intact inner membrane (Alder et al., 2023). One explanation for this and our observations is that the prolonged persistence of our HbVs might have led to the disruption of the inner membrane. Alternatively, it is possible that V-ATPase is already active in HbVs before fusion which also would explain both observations. Whether HbVs acidify has not been tested but – as for instance shown in our work here - they uniformly lack hemozoin and the appearance of their content does not noticeably change, indicating that also after inhibiting their fusion for many hours, HbVs do not display signs of profound digestion activity. Interestingly, we did observe 3 instances of HbVs fusing with the DV. This would further support direct fusion of the HbVs with the DV. It is possible that such a fusion event could be captured because fusion was impaired by the RBNS5L KS but due to this might also not progress in a physiological manner. While the steps between cytostome and HbVs remains enigmatic, the absence of densities in the HbV intermembrane space suggests an exclusion mechanism to avoid cytostomal internalization of PVM proteins or active removal prior to the HbV stage captured in this work.

Beside the rupture of the inner membrane, a further indicator of a maturation processes could be the detected intraluminal bodies. Interestingly, the CryoET revealed much more diverse structures than previous TEM of fixed samples (Lazarus et al., 2008; Jonscher et al., 2019; Sabitzki et al., 2024) which also matches recent SBF-SEM work analysing HbVs in an ATG18 mutant (Schmitz et al., 2026). The nature and relevance of these bodies therefore remains unclear.

The CryoET data also indicated lack of a protein coat on the HbVs as known from smaller transport vesicles in model organisms (Bonifacino and Glick, 2004). However, such coats are typically lost before fusion with the target membrane. Nevertheless, their large size and the findings in this work pointing to a maturation of HbVs, overall do not favour a model where HbVs are pure transport vesicles.

The BioID results further support some degree of maturation of HbVs. Firstly, the BioID detected the Retromer component VPS35, in other organisms involved in endosomal retrieval (Seaman, 2012). Secondly, it detected both Rab11 isoforms of the parasite, albeit each reached significance with only one of the HbV induction methods. Interestingly, each induction method also resulted in the detection of a different TBC domain protein. Our results with PfTBC8 and Rab11a indicate that PfTBC8 is a GAP for Rab11a (based on detection in the PfTBC8 BioID and changes in Rab11a location after PfTBC8 KS) and it is possible this is also the case for Rab11b and TBC7. However, whether a process akin to a Rab switch between Rab11a/TBC8 and Rab11b and potentially TBC7 occurs in HbVs, remains to be determined.

As GAPs are often key regulators of vesicular trafficking and PfTBC8 was one of the proteins from the HbV BioID without known function, we here chose it for functional analysis. Interestingly, conditional inactivation of PfTBC8 by KS showed a new endosomal transport phenotype, neither resembling HbV accumulation phenotypes of other endosomal transport proteins described nor the prevention of HbV found with cytostomal proteins (Spielmann et al., 2020). The non-uniform structures containing HCC after PfTBC8 KS might be the result of a global imbalance of the endosomal transport pathway which would be congruent with a recycling role, a canonical function of Rab11a in model organisms. As Rab11a was partially mislocalised after PfTBC8 KS, such a role is plausible. Work in *P. berghei* and in *T. gondii* has implicated Rab11a in dense granule transport, protein transport to the IMC, post Golgi secretory trafficking and transport to the PPM (Patil et al., 2020; Agop-Nersesian et al., 2010; Venugopal et al., 2020; McNamara et al., 2013). Interestingly, previous work studying TBC proteins in *T. gondii* detected TgTBC8 at the IMC (Quan et al., 2024). While we did not find any evidence for PfTBC8 at the IMC, the connection of Rab11a to functions of the IMC might be relevant should the connection between these two proteins detected here for *P. falciparum* be conserved in *T. gondii*. In conclusion, while our data shows that PfTBC8 is required for endosomal trafficking, the idea that this occurs through regulation of Rab11a and Rab11a-dependent recycling remains to be substantiated. It is also puzzling that we did not detect PfTBC8 at HbVs, although this might also be due to the induction method as this Rab was only detected in the RBSN5L KS-induced HbV, a condition we could not use to validate PfTBC8 HbV association. However, as Rab11a was detected at HbV, the functional connection of PfTBC8 with the HbV pathway is plausible.

A number of proteins involved in endosomal transport (defined by the appearance of HbVs in the parasite cytosol when these proteins are inactivated) are now known (Jonscher et al., 2019; Mukherjee et al., 2022; Sabitzki et al., 2024; Schmidt et al., 2023; Schmitz et al., 2026). Interestingly, few of these proteins were significantly enriched in our HbV surface BioID. One reason could be that these proteins are not located at the HbV but act indirectly, e.g. by functioning in a trans Golgi to HbV pathway that controls HbV maturation or mediating fusion at the DV, which might be the case for VPS45 (Cowles et al., 1994; Piper et al., 1994; Jonscher et al., 2019). It is also possible that our BioID is biased because it detects proteins only on the ∼70% of HbVs that contain PI(3)P which may represent a specific maturation state. However, it is important to note that PI(3)P is not only located at the DV but also co-localises with RBNS5L and Rab5b in DV proximal regions (Sabitzki et al., 2024) which may correspond to the DV proximal pool of VPS45 (Jonscher et al., 2019). Due to the subtraction of the HbV vs control condition in our BioID, it might be that generally PI(3)P-associated proteins are not enriched, because they are at such sites in both conditions. This might explain the absence of e.g. Rab5b which we here re-confirmed at the HbVs. It is therefore possible that our BioID preferentially selected for recycling components that disappear with further maturation of the HbVs or are not specifically at PI(3)P positive membranes. This might also be a reason for the strong enrichment of plasmepsins in our BioIDs. Under control conditions most plasmepsin molecules are in the DV, without pro-peptide reaching into the cytoplasm (Francis et al., 1997) and hence not available for biotinylation by PX-p40-miniTurbo. In contrast, in the HbV condition, plasmepsins still contain the pro-peptide part reaching into the cytoplasm and are available to be biotinylated on the HbV surface.

Using Halo-tracking of PM2 we provide strong support for the long-suspected trafficking route of DV proteases by endocytosis-mediated re-internalisation after default secretion to the parasite periphery (Klemba et al., 2004). We were able to detect and track PM2 through all stations of the pathway, the ER, the PPM, the cytostome, the HbVs and the DV. These findings indicate that endocytosis of the parasite has other cargos than the content of the host cell and PM2-Halo can be used as a marker for these trafficking steps. However, our work also provided indications that PM2 might reach the DV more directly. While it is possible this arises from residual endocytosis after KIC7 KS, the parasite might also posses a parallel transport pathway. Parallel pathways are also known to exist in vesicular trafficking in other organisms (Pfeffer, 2009; Cao et al., 2012). One option is that the direct PM2 transport route to the DV is the ancestral one and the high rate of endocytosis in malaria blood stages provided the possibility to increase capacity and tailor plasmepsin transport to hemoglobin uptake rates which might be advantageous given the function of these proteases in Hb degradation.

In summary we here report a new endosomal transport phenotype that may be the result of impaired recycling due to functional inactivation of PfTBC8. Furthermore, we identifed several more proteins in the HbV BioID that may be involved in endosomal transport processes. Taken together our results support a model where double membraned HbVs fuse with the DV, potentially after rupture of the inner membrane. This would be most reminiscent of the model proposed by Milani and colleagues (Milani et al., 2015) and include transport of parasite proteins as shown here for plasmepsins. We also detected the presence of canonical recycling components at the HbVs such as VPS35 and both Rab11 isoforms. The impact of PfTBC8 KS, likely via Rab11a, would be congruent with a recycling function. Together these findings indicate that components are removed from the HbV, overall supporting existence of a maturation pathway.

## Acknowledgments

This work was funded by the GRK2771 from the German Research Foundation (No. 1025453548970) to ALR, GF, TG and TS and grants to CMH from the G. Harold & Leila Y. Mathers Foundation and the Pew Biomedical Scholars Program. JD and TS acknowledge funding by the European Research Council (ERC) under the European Union’s Horizon 2020 research and innovation program (grant agreement number 101021493). We thank Jasmin Jansen, Frank Stein and Per Haberkant from the EMBL Proteomics Core Facility (EMBL Heidelberg) for the mass spectroscopy services and the FACS Core Facility (BNITM) for flow cytometry equipment and service.

## MATERIALS AND METHODS

### Cloning and plasmids used

Plasmids were generated using Gibson Assembly (Gibson et al., 2009) based on pSLI plasmids or modified pARL1 plasmids generated previously (Cubillán-Marín et al., 2025) (Supplemental file 1). Previously published cell lines used either directly or for co-transfections were RBNS5L-sw^endo^ and Rab5b-sw^endo^ (Sabitzki et al., 2024), GFP-2xFKBP-Rab5^endo^ (Birnbaum et al., 2017), and KIC7-sw^endo^ (Birnbaum et al., 2020).

### Parasite Culture transfection and selection linked integration

*P. falciparum* parasites were cultured at 37 °C in 5% O₂, 5% CO₂, and 90% N₂ in Albumax-supplemented culture medium containing human O⁺ erythrocytes (Universitätsklinikum Hamburg-Eppendorf (UKE), Hamburg, Germany). Cultures were grown in 2-, 5-, or 10-mL Petri dishes or in 50-mL culture flasks. For transfection, *P. falciparum* cultures (5-10% parasitemia) were enriched for schizont-stage parasites using 60% Percoll and transfected with 50 μg plasmid DNA using an Amaxa Nucleofector™ 2b (Lonza; program U-033) as described (Moon et al., 2013). Drugs for selection of transfected parasites and maintenance of transgenic parasites were 4 nM WR99210, 400 μg/mL G418, 2 μg/mL blasticidin S (BSD), or 0.9 μM DSM1, as appropriate. Parasite growth and parasitemia were routinely monitored by microscopic examination of Giemsa-stained blood smears.

SLI was performed as previously described (Birnbaum et al., 2017) to achieve genomic integration of episomal plasmids. Following the reappearance of episomal transfectants, parasite cultures were adjusted to a parasitemia of 4-10% and integrants selected with 400 μg/mL G418. For verification of correct integration, genomic DNA was isolated, and modification of the target locus verified by diagnostic PCR as described (Birnbaum et al., 2017).

### Light and Fluorescence Microscopy

Light and fluorescence microscopy were performed as previously described (Grüring and Spielmann, 2012) with minor modifications. Images of Giemsa-stained blood smears were generated with a Zeiss Axio Lab A1 light microscope equipped with a 100×/1.4 oil immersion objective and a Zeiss AxioCam ERC 5S camera. Images were acquired using Zen 2.3 software.

Live-cell fluorescence imaging was done with Hoechst 33342 (50 ng/mL; Invitrogen) or Halo substrate stained parasites as appropriate. For labeling of Halo-tagged proteins, the parasites were incubated with Halo substrates JF673-Halo or JF585 HaloTag (Cy5 channel) or JFX570-Halo or JFX650 HaloTag (Alexa Fluor 568 channel) at a final concentration of 0.5 nM for 10 min at 37 °C. Removal of excess dye prior to imaging or wash out for additional staining was done by three washing steps in RPMI medium.

Fluorescence microscopy was performed using a Zeiss Axio Imager M1 or M2 microscope equipped with an LQ-HXP 120 light source and Zeiss filter sets 44, 49, 64, and 50. Images were acquired using either 63× or 100× oil immersion objectives (NA 1.4) and a Hamamatsu Orca C4742-95 camera, controlled by AxioVision software (versions 4.7 and 4.8). Image processing, including brightness adjustments, channel merging, cropping, and figure assembly, was performed using Corel Photo-Paint (version 2019) and CorelDRAW (version 2019). Line intensity profiles were generated using ImageJ (Schneider et al., 2012).

### Flow Cytometry based growth assay and induction of KS

Asynchronous parasite cultures were adjusted to 0.05% parasitemia and split into a control and KS culture dish, KS induced by treatment with 250 nM rapalog and parasite proliferation was assessed every 24 h for five days as described (Birnbaum et al., 2017) based on a previously described flow cytometry method (Malleret et al., 2011). Briefly, 20 μL of parasite culture was incubated in 80 μL RPMI containing 0.5 mg/mL DHE and 0.45 mg/mL Hoechst 33342 for 20 min, staining quenched by addition of 400 μL RPMI supplemented with 0.003% glutaraldehyde, and samples analyzed using an LSR II flow cytometer.

### Bloated food vacuole assay (bloated DV assay) and synchronization of parasites for KS, inhibitor and staining assays

The bloated DV assay was essentially as described previously (Jonscher et al., 2019). Parasites were synchronized using a protocol of Percoll schizont purification followed by 5% sorbitol treatment as described (Grüring et al., 2011) with modifications. Schizont-stages were enriched using a 60% Percoll gradient incubated for 1 h at 37 °C with shaking (800 × g) in a mixture of 250 μL infected blood and 500 μL culture medium followed by an additional 7 h of culturing under standard conditions to permit parasites to invade. Synchronization using 5% sorbitol resulted in 0–8 h post invasion (hpi) rings that were cultured for a further 18 h to reach 18–24 hpi. Cultures were then split into two dishes, KS induced in one dish and E64 added to both conditions at a final concentration of 33 μM. After 8 h of incubation, differential interference contrast (DIC) microscopy images were acquired (operator blinded to the identity of the sample). For other synchronization regimens the timing was adapted to obtain the stages indicated in the respective figures, including addition of rapalog (250 nM), CytD (10 μM) and Halo dyes (0.5 nM).

### CytD assay to assess impact on endocytosis before HbV generation

CytD vesicle accumulation was done as previously described (Sabitzki et al., 2024) using PfTBC8-sw^endo^ parasites co-expressing the 1xNLS mislocaliser. The parasites were synchronized twice 8 h apart using 5% sorbitol to generate parasites of 10–18 hpi and then divided into four 2 mL culture dishes. PfTBC8 KS was induced in two dishes at 24–32 hpi by addition of 250 nM rapalog. One hour later, 10 μM CytD was added to one rapalog-treated and one untreated dish. After a further 5 h continued culture at 37°C, the parasites were imaged by DIC microscopy, and vesicle numbers were quantified (operator scoring the cells was blinded to the identity of the condition the cells originated from).

BioIDs

### Biotinylation of HbV surface proteins and preparation of extracts

For BioIDs, the parasites were grown in biotin-free RPMI-1640 medium (Biozol #USB-R9002-01). To generate the HbV surface proteome by BioID, RBNS5L^endo^ parasites episomally transfected with a plasmid expressing the P40x-miniTurbo construct (Ramón-Zamorano et al., 2026) and the 1xNLS mislocaliser (Fig. 2A) were synchronized by two sequential sorbitol treatments to obtain parasites with a 10–18 hpi stage window. At 22-30 hpi, cultures were divided into three experimental groups: rapalog-treated, cytochalasin D-treated, and untreated control. Biotin (50 μM) was added to all cultures at 27-35 hpi. Vesicle accumulation and the localization of the episomally expressed miniTurbo fusion proteins were verified by microscopy at 28-36 hpi. Parasites were harvested by centrifugation at 2,200 × g for 10 min at room temperature, washed twice with D-PBS and parasites released from their host cell using 0.03% (w/v) saponin. The parasites were washed 5 times with D-PBS or further until the supernatant was clear. The final parasite pellet was resuspended in lysis buffer (50mM Tris-HCl pH 7.5, 500 mM NaCl, 1% Triton-X-100) supplemented with 1 mM dithiothreitol (DTT), 2× protease inhibitor cocktail, and 1 mM phenylmethylsulfonyl fluoride (PMSF), after which SDS was added to a final concentration of 0.4%. Samples were then stored at −80°C for a minimum of 30 min or overnight before further processing.

Biotinylation and extracts for DiQ-BioID of the PfTBC8sw^endo^ parasites were performed as previously described (Birnbaum et al., 2020; Kimmel et al., 2023) but using a miniTurbo biotinylizer (Ramón-Zamorano et al., 2026). Samples were split into one condition with rapalog (biotinylizer recruited to PfTBC8) and one without and biotinylation carried out for 30 min by adding 50 μM biotin and extracts prepared as described for the HbV surface BioID.

### Preparation of Protease-Resistant Streptavidin Beads

Streptavidin Sepharose beads were chemically modified to enhance protease resistance as described by others (Rafiee et al., 2020). Briefly, the beads were washed twice with PBS containing 0.1% Tween 20 (PBS-T). For cyclohexanedione (CHD) treatment, the beads were resuspended in CHD solution prepared in PBS-T (pH 13) and incubated for 4 h at room temperature under continuous rotation. Following CHD modification, the beads were washed and subjected to reductive methylation by incubation in 4% formaldehyde and 0.2 M sodium cyanoborohydride prepared in PBS-T for 2 h at room temperature. The modified beads were then washed sequentially with 0.1 M Tris-HCl buffer (pH 7.5) and PBS-T before being resuspended in PBS-T. Finally, the protease-resistant beads were stored at 4 °C until further use.

### On-Bead Digestion

Beads were resuspended in elution buffer consisting of 2 M urea and 10 mM DTT in 100 mM Tris-HCl (pH 7.5) and incubated for 20 min at room temperature with shaking. Proteins were subsequently alkylated by adding iodoacetamide to a final concentration of 50 mM and incubating the samples for 10 min in the dark. 0.25 μg of Trypsin/LysC was then added, and samples were digested for 2 h at room temperature with shaking. Following digestion, the supernatant was collected, and the beads were subjected to a second elution step. The eluates were combined and supplemented with additional Trypsin/LysC (0.1 μg) for overnight digestion at room temperature. Digested samples were stored at -80 °C until shipment to the European Molecular Biology Laboratory (EMBL, Heidelberg) for further processing.

### Mass Spectrometry and Protein Enrichment Analysis

Peptide separation and MS spectra with an Orbitrap Fusion Lumos and data analysis using FragPipe (Kong et al., 2017) and a Uniprot *Plasmodium falciparum* isolate 3D7 fasta database were done as described (Guillén-Samander et al., 2026). The raw output files of FragPipe (protein.tsv files) were processed using the R programming language (ISBN 3-900051-07-0). Contaminants and reverse proteins were filtered out and only proteins that were quantified with at least 2 razor peptides (Razor.Peptides >= 2) were considered for the analysis. 1501 proteins passed the quality control filters. Log2 transformed raw TMT reporter ion intensities (’channel’ columns) were first cleaned for batch effects using the ‘removeBatchEffect’ function of the limma package (Ritchie et al., 2015) and further normalized using the ‘normalizeVSN’ function of the limma package (VSN - variance stabilization normalization - (Huber et al., 2002)). Proteins were tested for differential expression using a moderated t-test by applying the limma package (’lmFit’ and ‘eBayes’ functions). The replicate information was added as a factor in the design matrix given as an argument to the ‘lmFit’ function of limma. The t-value output of limma for certain statistical comparisons was analyzed with the ‘fdrtool’ function of the fdrtool packages (Strimmer, 2008) in order to extract p-values and false discovery rates (q-values were used). A protein was annotated as a hit with a false discovery rate (fdr) smaller 0.05 and an absolute fold-change of greater 1.5 and as a candidate with a fdr below 0.2 and an absolute fold-change of at least 1.

### Transmission Electron Microscopy (TEM)

*P. falciparum* parasites were synchronized by Percoll purification (60%) followed by sorbitol treatment 6 h later, yielding parasites between 0 and 6 h post-invasion (hpi). PfTBC8 KS was induced at 20–26 hpi. At 32–38 hpi, infected erythrocytes were enriched using 80% Percoll and fixed in 2.5% glutaraldehyde (Electron Microscopy Sciences) prepared in 50 mM cacodylate buffer (pH 7.2) for 30 min at room temperature. Samples were subsequently post-fixed with 2% osmium tetroxide (OsO₄; Electron Microscopy Sciences) in distilled water for 40 min on ice in the dark. Membranes were contrasted with 0.5% uranyl acetate (Agar Scientific) in distilled water for 30 min at room temperature. Samples were then dehydrated through a graded ethanol series (50– 100%) and embedded in epoxy resin (EPON; Carl Roth GmbH & Co. KG). Ultrathin sections (60 nm) were prepared using an Ultracut UC7 ultramicrotome (Leica Microsystems) and imaged on a Tecnai Spirit transmission electron microscope (FEI) equipped with a LaB₆ filament and operated at an accelerating voltage of 80 kV. Digital electron microscopy files (.emi/.ser) were converted to 8-bit TIFF images using the TIA Reader plugin for ImageJ.

### Phylogenetic tree generation

TBC homologues in reference genomes of *T. gondii, N. caninum, T. annulata, E. falciformis, B. microti, P. berghei, P. gallinaceum* and *P. falciparum* were identified using BLAST and HHpred. Protein sequences were retrieved from VEupathDB (release 67) (Amos et al., 2022), and a maximum-likelihood phylogenetic tree was generated in MEGA11 (Tamura et al., 2021) with 100 bootstrapping repeats from their alignment.

### Cryo-ET Methods

#### Parasite culture and synchronization for cryo-ET

RBNS5L-sw^endo^ KS parasites (Sabitzki et al., 2024) were cultured following a protocol adapted from (Duffy and Avery, 2018). Parasites were cultured in O+ or A+ erythrocytes (New York Blood Center), with continuous gentle shaking, at 37 °C under 5% O_2_, 5% CO_2_ and 90% N_2_ in complete RPMI (cRPMI): RPMI 1640 medium (Sigma, R6504-50L) supplemented with 0.05 mg ml−1 hypoxanthine (Sigma, H9377-100G), 0.06 mg ml−1 sodium hydroxide (Fisher Scientific, SS255-1), 0.8 mg l−1 thymidine (Sigma, T1895-1G), 0.04 mg ml−1 sodium pyruvate (Sigma, P5280-256), 2.25 mg ml−1 sodium bicarbonate (Sigma, S6014-500G), 5.9 mg ml−1 HEPES (Sigma, H4034-500G), 0.67 mg ml−1 glucose (Sigma, G7021-1KG), 0.01 mg ml−1 gentamycin (Sigma, G1271-100ml) and 0.5% Albumax II (Gibco, 11021-045), at a hematocrit of 2% at <10% parasitemia.

To enrich trophozoites, parasites were synchronized using a cushion of 65% Percoll (Cytiva, 17089109) gradients twice to target a 6-hour invasion window. Following a 24-hour recovery, cultures were treated with 250 nM rapamycin (Clontech, no. 635055) for 5 hours. After treatment, parasites (29-35 hpi) were pelleted at 800 × g for 3 min at room temperature and then enriched by gelatin flotation as described previously (Goodyer et al., 1994) in 0.7% gelatin at 37 °C for 1 h. The trophozoite-containing supernatant was collected, washed and resuspended in 1 ml of cRPMI.

### Grid freezing and CLEM

3.5 μL of iRBCs in cRPMI were applied to glow-discharged Quantifoil R2/2 Cu 200 grids, back-blotted with Whatman Grade 1 filter paper, and plunge frozen in liquid nitrogen-cooled liquid ethane using a manual plunger. After clipping, grids were assessed for ice thickness and cell density using a Leica EM Cryo CLEM.

### FIB milling

Lamellae were milled as previously described on an Aquilos cryo-FIB/SEM (Thermo Fisher Scientific) (Anton et al., 2025). Briefly, grids were sputtered with platinum metal and coated with an organometallic platinum layer using the gas injection system (GIS). Relief trenches were milled at 40° and subsequent lamella thinning was performed at 7°. Target lamellae sites were manually milled in parallel at 30 kV with steps of 0.5 nA, 0.3 nA, 0.1 nA, 50 pA, and 10 pA. Lamellae for cryo-ET and cryo-STET were polished to a target thickness of 150 nm and 400-800 nm, respectively.

### Cryo-ET data collection

Data collection was performed on a Titan Krios G3 (Thermo Fisher Scientific) operated at 300 kV and equipped with a K3 detector and BioQuantum energy filter (Gatan) with a slit width of 20 eV. Tilt series were acquired using SerialEM from -53° to 67° with a pre-tilt of 7° in 3° increments, using a dose-symmetric scheme at 53,000× magnification (0.8475 Å/pixel super-resolution), a target defocus of -3 μm and a total fluence of 120 e^-^/Å^2^. Individual tilts were recorded with constant beam intensity in 10 frames as LZW-compressed TIF format. Data collection statistics are listed in Table S3.

### Cryo-ET data processing

Tilt series motion correction, CTF estimation, and tilt stack creation were performed in Warp 1.0.9 (Tegunov and Cramer, 2019). Tilt series alignment and tomogram reconstruction were performed in AreTomo 1.3.4 with SART at bin 6 (10.17 Å/px) and inspected in IMOD 4.12.60 (Mastronarde and Held, 2017; Zheng et al., 2022). Tomograms were denoised and corrected for missing wedge using IsoNet (Liu et al., 2022). Membranes were segmented using MemBrain-Seg and cleaned in ChimeraX (Goddard et al., 2018; Lamm et al., 2024).

### Intermembrane distance determination

To quantify distance between adjacent membrane layers, we developed a custom MATLAB-based pairing algorithm. Coordinates points and surface normal vectors defining the membrane were generated from tomographic membrane segmentations using a membrane oversampling script by Pyle et al. with in-house modifications (Pyle et al., 2022). The algorithm utilized these coordinates and refined Euler angles to perform a neighbor search on 2D tomographic slices. For each particle on membrane, the surface normal vector, n, was extracted from the Euler angles. A search line, L, was projected from the particle center, (x0, y0), along the direction of the normal vector (Fig S1E, F).

The search was constrained to positive displacement from the particle coordinates along the normal vector to ensure directional specificity toward the anticipated neighboring layer, extending to a maximum search radius of 200 pixels (approximately 680 Å). A neighboring particle was successfully paired if it was located within a 2-pixel lateral tolerance of this projected line. To ensure unique 1:1 matching and avoid redundant measurements, both particles were removed from the search pool immediately upon pairing. Inter-membrane separation was defined as the 2D Euclidean distance between the centers of each matched pair, with mean values and standard deviations aggregated across all processed tomograms. The complete set of matched pairs underwent manual examination and selection. To ensure equal representation, the resulting measurements were averaged for each tomogram, preventing any individual vesicle from skewing the dataset.

### Vesicle diameter determination

Vesicle diameters were quantified using two approaches depending on their inclusion within the tomographic field of view. For vesicles fully captured within the tomograms, 3D diameters were measured from tomogram segmentations in UCSF Chimera by placing coordinate markers at opposing boundaries using the Volume Tracer and Structure Measurements tools. For larger vesicles that were only partially captured within a single tomogram, 2D diameters were manually measured from lower-resolution montage overviews using ImageJ.

### Statistical Analysis of CryoET data

Data were parsed and analyzed using custom Python scripts. To evaluate potential linear relationships between vesicle maximum diameter and intermembrane distance, Pearson correlation coefficients (r) were calculated. To compare vesicle max diameter and intermembrane distance across the four categorical feature groups, a One-Way Analysis of Variance (ANOVA) was performed, followed by Tukey’s HSD post-hoc tests for multiple comparisons. Statistical significance was defined as P < 0.05.

### Figures and plots

CryoET figures were generated in Adobe Illustrator and graphs were plotted in Python. ChimeraX was used to visualize segmentations. Central slice images were generated in IMOD. Lamella montages were compiled in Adobe Illustrator. All other figures were generated in Corel Draw v21. Fluorescence intensities were measured in Fiji ImageJ using the Plot Profile function. Images of individual parasite were processed using Corel Photo Paint v21, where cropping and adjustments of brightness and intensity were performed if required.

