## Supplementary material for "Characterization of Host Cell Cytosol-filled Vesicles in the Human Malaria Parasite *Plasmodium falciparum*": Supplmental figures

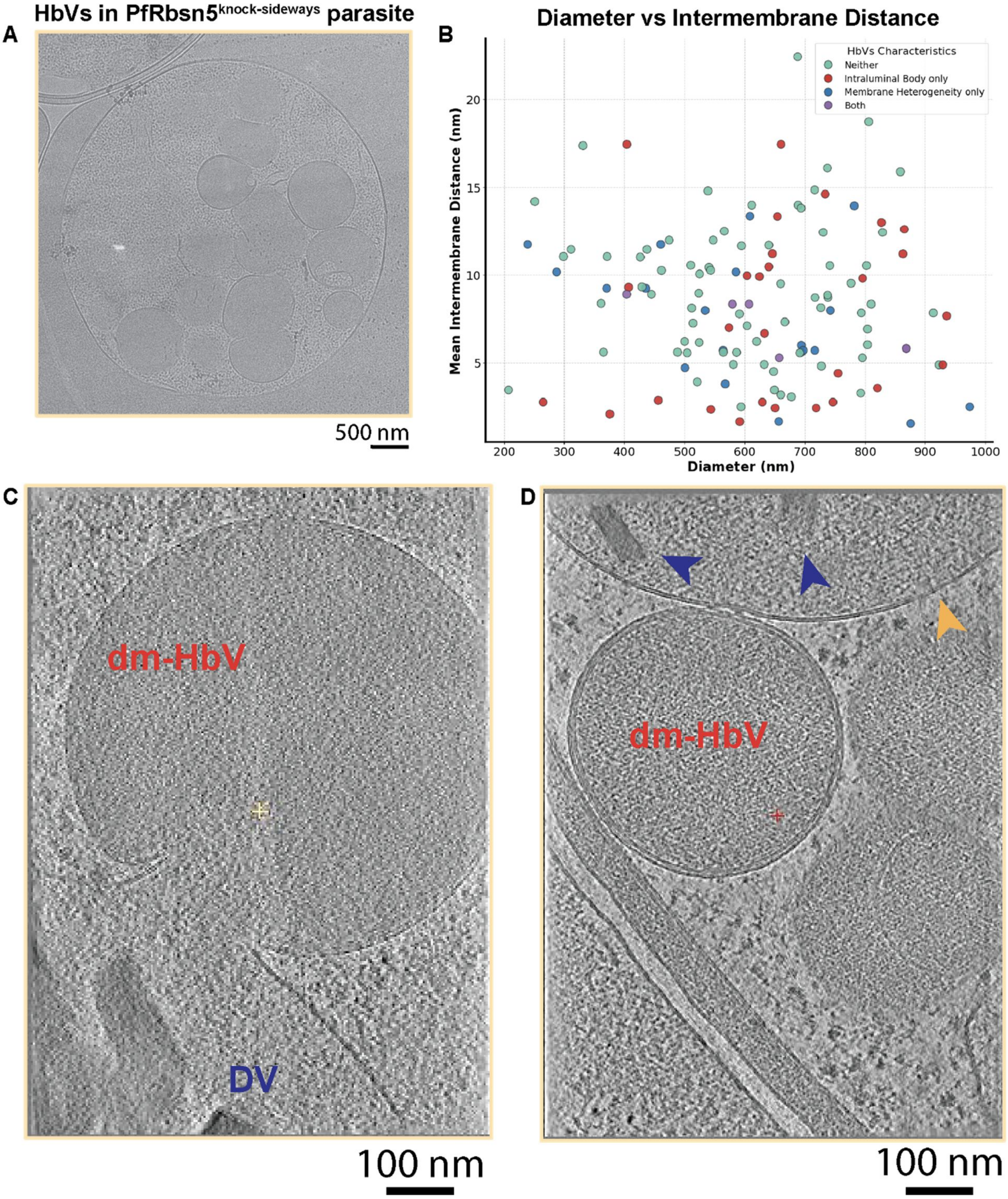


Figure S1 - **HbV characteristics and fusion with the DV. A**, Representative lamella overview of the region shown in Fig. 1A illustrating the accumulation of HbVs within the parasite. **B**, Scatter plot illustrating the relationship between vesicle diameter (nm) and mean intermembrane distance (nm). Individual data points represent single vesicles, color-coded by their structural classification: baseline HbVs (green), HbVs with intraluminal body only (red), HbVs with membrane heterogeneity only (blue), and HbVs with both features (purple). No significant correlation was observed between maximum vesicle diameter and mean intermembrane distance (Pearson $r = -0.074$, $p = 0.4093$). **C**, Averaged central slice from a tomogram showing double-membraned hemoglobin-filled vesicle (dm-HbV) in the process of fusing with the DV. **D**, Averaged central slice from a tomogram collected on another cell showing a single-membraned vesicle (yellow arrowhead) containing both host hemoglobin and hemozoin crystals.

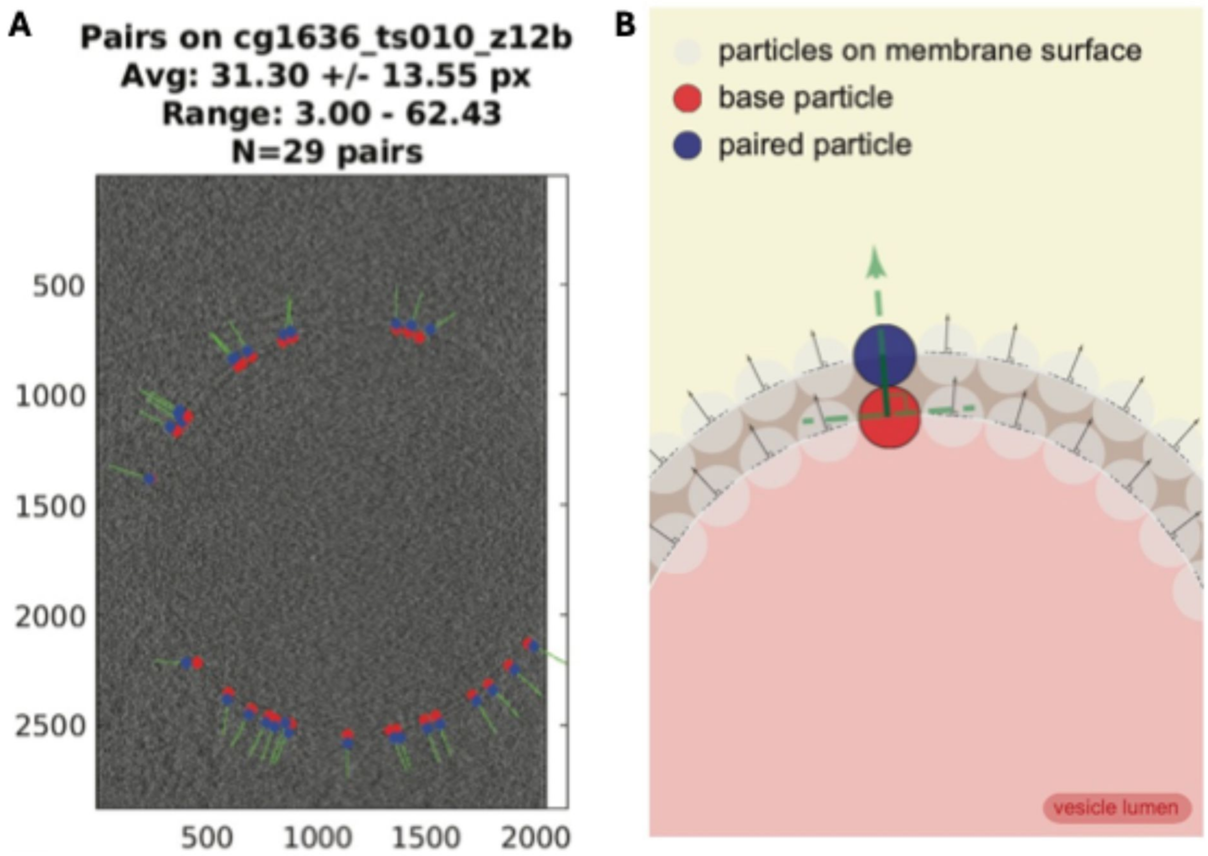


Figure S2 - **Intermembrane distance measurement method illustration A**, Representative micrograph showing the automated pairing of membrane points for analysis. **B**, Schematic defining the intermembrane distance determination strategy.

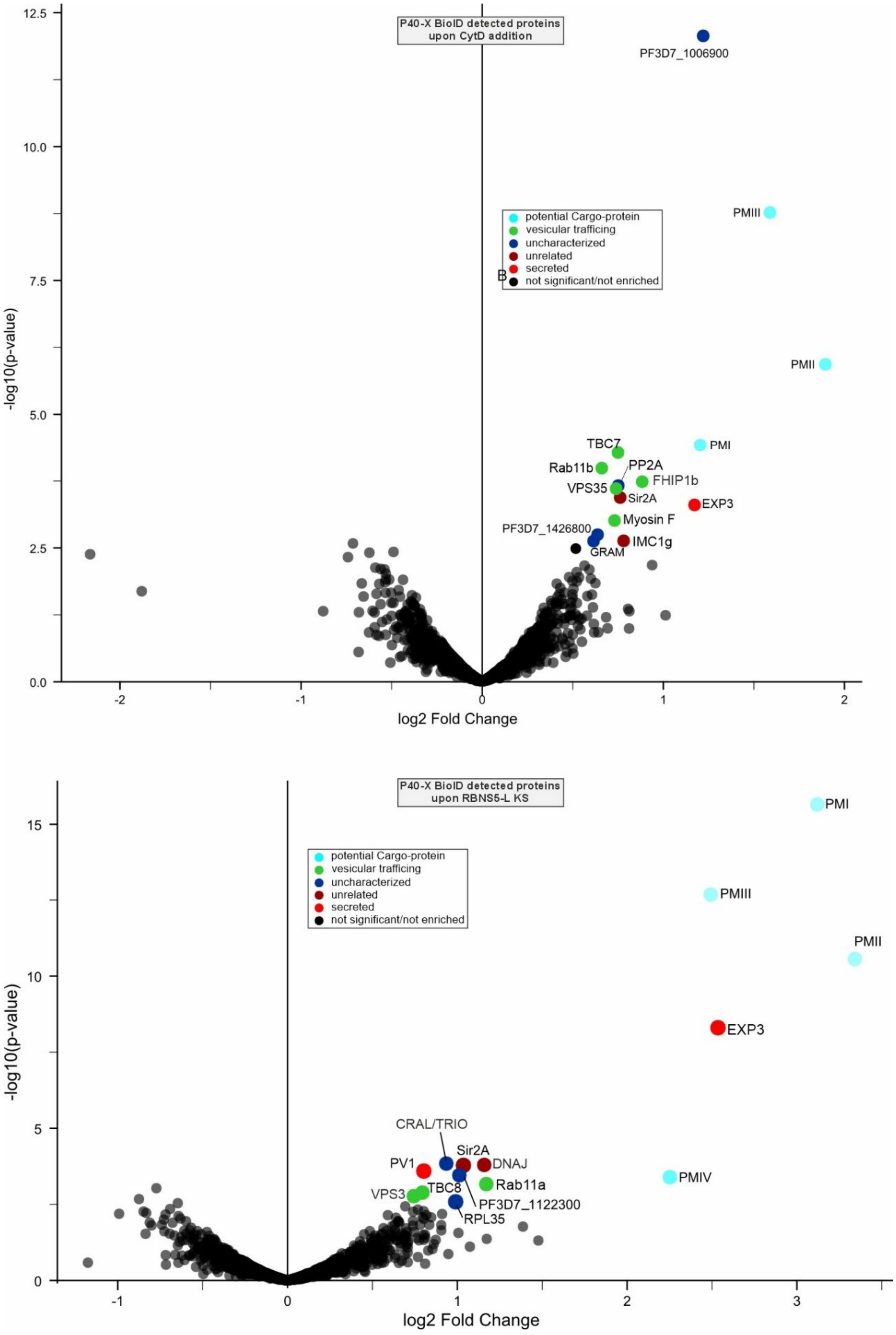


Figure S3 - **Full plots of the PI(3)P-based HbV surface proxiome.** Full plost of the results shown in Fig. 2E. Details as indicated in Figure 2.

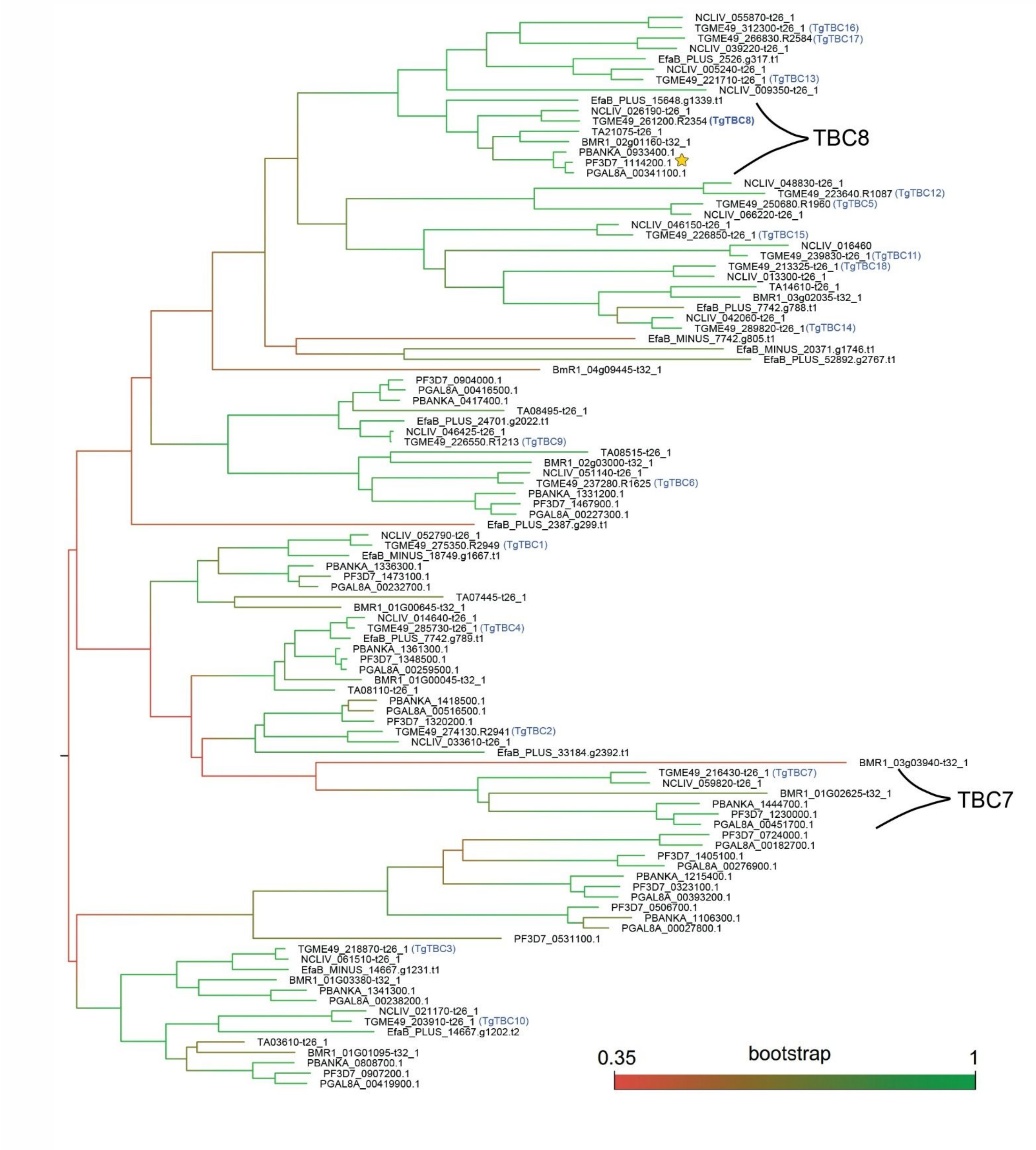


Figure S4 – **Phylogenetic tree of TBC domain proteins.** Phylogenetic tree of the TBC proteins found in chosen apicomplexan species using BLAST and HHpred. *T. gondii* annotations (PMID: 38713739) are indicated in blue next to its corresponding gene ID. Branches are colored by bootstrap value and the ones corresponding to TBC8 and TBC7 homologues, mentioned in this article, are indicated.

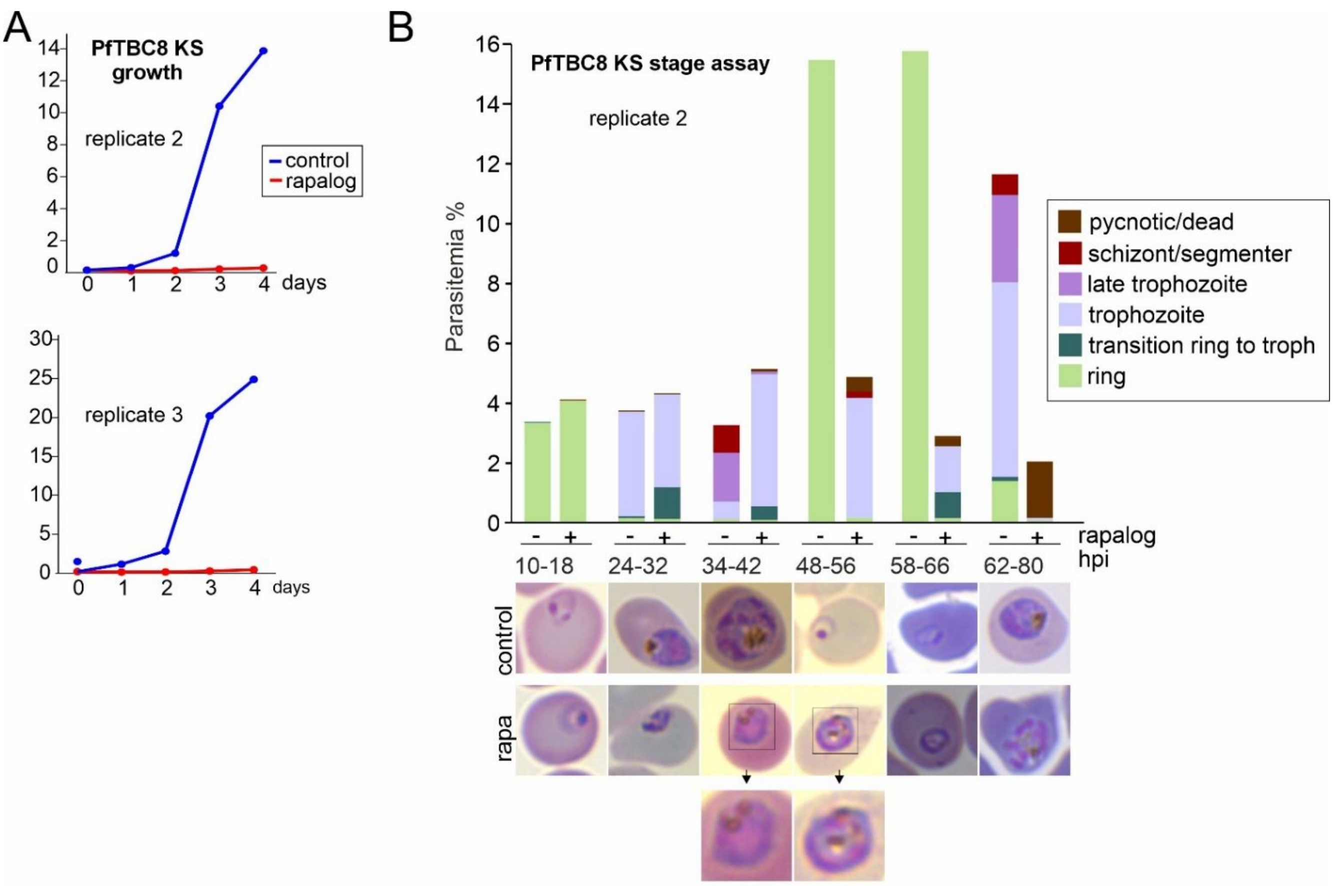


igure S5 – **Additional replicates of growth and stage assays of the PfTBC8 KS. A)** Independent replicates of growth curves shown in Figure 4E. **B)** Second independent replicate of the stage assay shown in Figure 4F. Details as indicated in Figure 4.

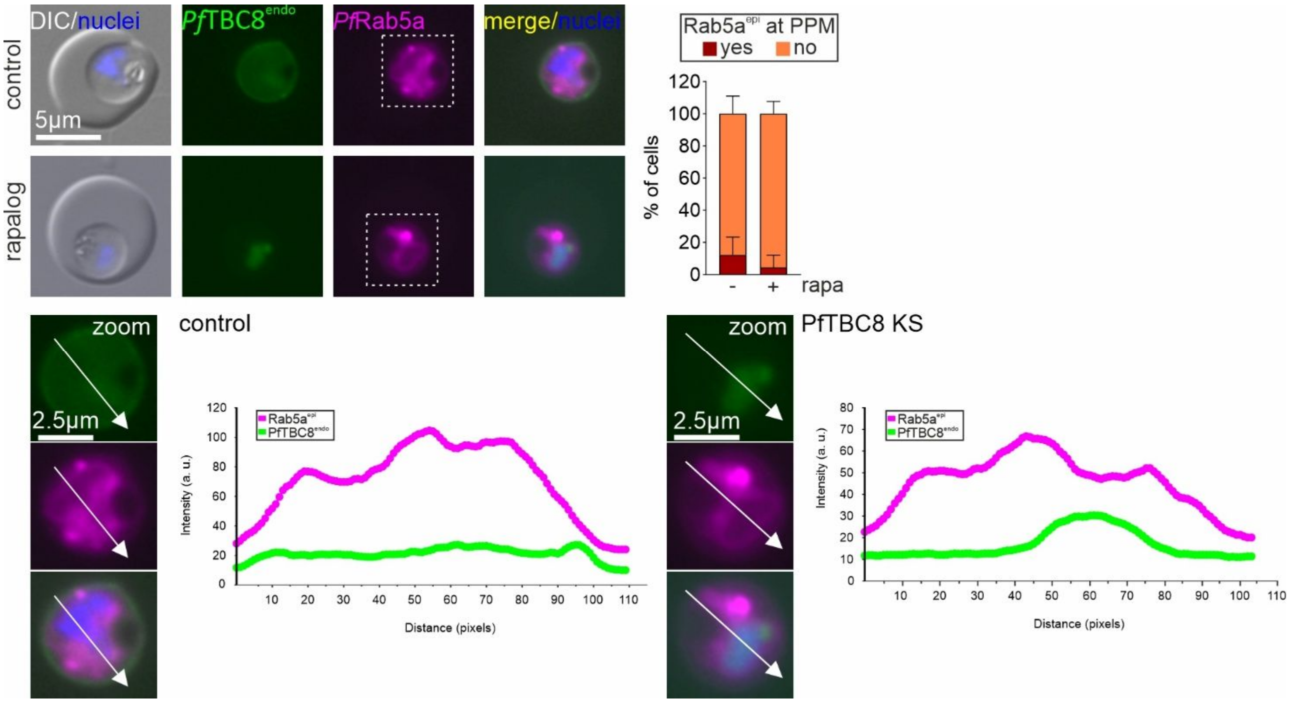


Figure S6 – **Rab5a does not change its localization upon PfTBC8 KS. A)** Representative live-cell fluorescence microscopy images of PfTBC8-sw$^{endo}$ (green channel) episomally co-expressing Rab5a (magenta channel) upon PfTBC8-sw KS (rapalog) and under control conditions (control). Zoom shows enlarged area (last three images) of respective GFP and magenta channel or merge/nuclei image. White arrows indicate line used to calculate the intensity values shown in. Example intensity plots of the PfTBC8-sw$^{endo}$ signal (green) and the Rab5a signal (magenta) (arbitrary units, a.u.) are shown. Bar graphs show quantification of number of cells (%) showing plasma membrane-associated Rab5a signal with rapalog (PfTBC8 KS) or not (control) (n = 3 independent replicates. Nuclei were stained with Hoechst, merge: overlay of green and magenta channel. Scalebar indicated. DIC, differential interference contrast.
