## supplemental File 1 for "Characterization of Host Cell Cytosol-filled Vesicles in the Human Malaria Parasite *Plasmodium falciparum*"

**PRIMERS**

**Primer used for cloning**

>PMII-Halo fw:
CCAGTTGCAACATTAGGTACCATGGCAGAAATTGGTACG

>PMII Halo rv:
GCTGCCATATCCCTCGACCCGGGTTAACCTGAAATTTCTAATGTAC

>NLS Rab5a epi fw: CCAGATTATGCACCAGTTGCAACATTAGGTACCATGGAAAAGAAAAGTAGTTATAAAACAGTTTTATTAGG

>Rab5a epi Rv: CGAACATTAAGCTGCCATATCCCTCGACCCGGGTTAACAACATCCTTTTTTTGAAAGTGTTTGATTATC

>P40-fw:
GTTTTTTTTAATTTCTTACATATAACTCGAGATGGCTGTGGCCCAGC

AGCTGCGGGCCGAGAGTGACTTTGAACAGC

>P40-rv:
CCGCCTCCGCTACCTCCGCCTCCACTACCTCCACCTCCGTTTAAAC

CGCGGAGTGCCTGGGGCACCTGCTCTGAGTCATAGGGCG

>miniturbo_fw:
CAGAGCAGGTGCCCCAGGCACTCCGCGGTTTAAACGGAGGTGGAG

GTAGTGGAGGCGGAGGTAGCGGAGGCGGTGGGAGTGGTGGAGGC

GGAAGTGGCGGTGGA

>miniturbo_rv:
GCAACTGGTGCATAATCTGGATTATCATATGGATAACTTGTCCTAGG

CTTTTCGGCAGACCGCAGACTGATTTCTCCGCCC

>L4_KpnI_rRab11a fw: CCAGATTATGCACCAGTTGCAACATTAGGTACCGCTATGAAGGAGGACTACTACGACTATTTATTCAAGATAG

>rRab11a_XmaI rv: CGAACATTAAGCTGCCATATCCCTCGACCCGGGCTAGCAGCACTTATTCTTCTTCTTTGTCTCATTC

>rRab11b AvrII fw: GGTGCTGGAGGTGCAGGTAGAcctaggAGCAACGAGGAGTATGACCACTTATACAAG

>rRab11b StuI rv: TATGCTATACGAAGTTATaggcctTCAGCAGCACTTAACCTTGTTCATGTTGTTGTCC

>HR Rab11b [PF3D7_1320600] NotI * fw:
GGGgcggccgcTAATATCGAAGTATAACTAGCGC

>HR Rab11b [PF3D7_1320600] PmeI rv: TATTAACTTCTGCTCgtttaaacCCTGCTAATGGAATGGTTCTCGTAGCAAATTCTACAC

**Primer used for integration check PCRs**

>PfTBC8 int fw:
ATATATATATATATATATTTCTTTCAG

>PfTBC8 int rv:
TCAATATAACAAATAAGGAAATGATAC

>GFP85 rv:
ACCTTCACCCTCTCCACTGAC

>pARLsense55:
GGAATTGTGAGCGGATAACAATTTCACACAGG

>Rab11b int fw:
*tcaag*ATATGGGATACTGCCG

>Rab11b int rv:
CTAACAACATTTTACTTTATTC

**PLASMIDS**

**>pARL_crt-Px-p40-miniTurbo-mScarlet_nmd3-NLS-FRBT2A-DHODH**

CGGCCGCTAACGTAACAGACTTAGGAGGAGATCTTAGTAGTTGAGTGATTCTATATACATATACATAAATAAATTACATAATATTATAAATTTTTTTTATATTATATTAGAATGTTTACTATATAAAAATAAATATTTTCTTATATATTTTTTTATTTTTTCATATGAAAAAAGAAATTTTTTATATATTTTTTTATTATATTTTATAAATAAAAAAGAACTTATACCTATTATTATTATTATATATAAAAAATATATATTTTTTTATAAATTCCTATTTTTCCGATTATTTATTTATTTTTTTTTTTTATTGTTTAAAAATATATAAAAAAAATTCTTATATATTTAATATTATAAGTCCATAAATATATATAATTTATATAATTTATATATTTCCATACTTTTTTTATTATATATCATGAGATAAAATAAATTTCATATAGAAATCGTATTTATTTATTAATTTGATATATATTAATAAATAAATATTAATAATAATATGTTATATATATATATATTTATTTAATTATTATATAGGAAACTATATATATATATATATATTATATTTTTTTTTTTGAAATATAATAATATATAATATTATTCCTAAAAATATCTATATTCTTATACCTGAACCCTTTTTTTTTTTTTTTTTTTTTTTTGACTTTCGATTGTTCATTCTGTTTATTGTATAAATATATAAATATATATATATTGTATATTTTATTACATATATTTTTTTTTTTTGAAAGTTAAGAATTTTAACTTAATAAAAAAAGTACATATATTTTTAATAAATGTCCTCCATTATATAAATTGTTATATAAAAGATTTTATATAATTTAAATAGAATTCATTTATAAACAAATTTGTTTATAAAAATATATTTATGTATATATAATATAAATATATATATTTATATATATATATATATATATATTACTATATATATTTTTTTTTTTTTTTCCTTTTTTTTACTTTCCCAAGTTGTACTGCTTCTAAGCTTTTTTAATAAACATATATAATTTGTACAAATATTTTAGATTATATACATGATGTATATTTGAATATATTTTCTATATATTTGTGGTTCCATTTTTGTATATTATATATAATATATTTATATATATATTGATATGTCAATATTTGTATAACACATGAAGTTTTTGTTTTTTTTTTTTTTTTTTTTTAATGGAGAATATTTAATAATATATGAAAAAAATTTTATATAAATATATATATATATATATATATATATATATATATATATATATATATGTATATATATATATTTATATATACATGTATGTTTTTTAAAAAGTTAAATAATTCTATAGATTATTTTCATTGTCTTCACATATATGACATAAATATTTTAAAATCGACATTCCGATATATTATATTTTTAGACTATAATATCCGTTAATAATAAATACACGCAGTCATATTATTTATTATACATTCATTTATTATTTTGTTTTTTTTAATTTCTTACATATAACTCGAGATGGCTGTGGCCCAGCAGCTGCGGGCCGAGAGTGACTTTGAACAGCTTCCGGATGATGTTGCCATCTCGGCCAACATTGCTGACATCGAGGAGAAGAGAGGCTTCACCAGCCACTTTGTTTTCGTCATCGAGGTGAAGACAAAAGGAGGATCCAAGTACCTCATCTACCGCCGCTACCGCCAGTTCCATGCTTTGCAGAGCAAGCTGGAGGAGCGCTTCGGGCCAGACAGCAAGAGCAGTGCCCTGGCCTGTACCCTGCCCACACTCCCAGCCAAAGTCTACGTGGGTGTGAAACAGGAGATCGCCGAGATGCGGATACCTGCCCTCAACGCCTACATGAAGAGCCTGCTCAGCCTGCCGGTCTGGGTGCTGATGGATGAGGACGTCCGGATCTTCTTTTACCAGTCGCCCTATGACTCAGAGCAGGTGCCCCAGGCACTCCGCGGTTTAAACGGAGGTGGAGGTAGTGGAGGCGGAGGTAGCGGAGGCGGTGGGAGTGGTGGAGGCGGAAGTGGCGGTGGAGGTAGCGGAGGTGGAGGTAGTGGAGGCGGAGGTAGCacgcgtatcccgctgctgaacgctaaacagattctgggacagctggacggcgggagcgtggcagtcctgcctgtggtcgactccaccaatcagtacctgctggatcgaatcggcgagctgaagagtggggatgcttgcattgcagaatatcagcaggcagggagaggaagcagagggaggaaatggttctctccttttggagctaacctgtacctgagtatgttttggcgcctgaagcggggaccagcagcaatcggcctgggcccggtcatcggaattgtcatggcagaagcgctgcgaaagctgggagcagacaaggtgcgagtcaaatggcccaatgacctgtatctgcaggatagaaagctggcaggcatcctggtggagctggccggaataacaggcgatgctgcacagatcgtcattggcgccgggattaacgtggctatgaggcgcgtggaggaaagcgtggtcaatcagggctggatcacactgcaggaagcagggattaacctggacaggaatactctggccgctatgctgatccgagagctgcgggcagccctggaactgttcgagcaggaaggcctggctccatatctgtcacggtgggagaagctggataacttcatcaatagacccgtgaagctgatcattggggacaaagagattttcgggattagccgggggattgataaacagggagccctgctgctggaacaggacggagttatcaaaccctggatgggcggagaaatcagtctgcggtctgccgaaaagCCTAGGACAAGTTATCCATATGATAATCCAGATTATGCACCAGTTGCAACATTAGGTACCATGGTGAGTAAGGGTGAGGCAGTGATTAAGGAGTTTATGCGTTTTAAGGTGCACATGGAGGGTAGTATGAACGGTCACGAGTTTGAGATTGAGGGTGAGGGTGAGGGTCGTCCATACGAGGGTACACAGACAGCAAAGCTTAAGGTGACAAAGGGTGGTCCACTTCCATTTAGTTGGGATATTCTTAGTCCACAGTTTATGTACGGTAGTCGTGCATTTACAAAGCACCCAGCAGATATTCCAGATTACTACAAGCAGAGTTTTCCAGAGGGTTTTAAGTGGGAGCGTGTGATGAACTTTGAGGATGGTGGTGCAGTGACAGTGACACAGGATACAAGTCTTGAGGATGGTACACTTATTTACAAGGTGAAGCTTCGTGGTACAAACTTTCCACCAGATGGTCCAGTGATGCAGAAGAAGACAATGGGTTGGGAGGCAAGTACAGAGCGTCTTTACCCAGAGGATGGTGTGCTTAAGGGTGATATTAAGATGGCACTTCGTCTTAAGGATGGTGGTCGTTACCTTGCAGATTTTAAGACAACATACAAGGCAAAGAAGCCAGTGCAGATGCCAGGTGCATACAACGTGGATCGTAAGCTTGATATTACAAGTCACAACGAGGATTACACAGTGGTGGAGCAGTACGAGCGTAGTGAGGGTCGTCACAGTACAGGTGGTATGGATGAGCTTTACAAGTAACCCGGGTCGAGGGATATGGCAGCTTAATGTTCGTTTTTCTTATTTATATATTTATACCAATTGATTGTATTTATAACTGTAAAAATGTGTATGTTGTGTGCATATTTTTTTTTGTGCATGCACATGCATGTAAATAGCTAAAATTATGAACATTTTATTTTTTGTTCAGAAAAAAAAAACTTTACACACATAAAATGGCTAGTATGAATAGCCATATTTTATATAAATTAAATCCTATGAATTTATGACCATATTAAAAATTTAGATATTTATGGAACATAATATGTTTGAAACAATAAGACAAAATTATTATTATTATTATTATTTTTACTGTTATAATTATGTGTCTCCTTCAATGATTCATAAATAGTTGGACTTGATTTTTAAAATGTTTATAATATGATTAGCATAGTTAAATAAAAAAAGTTGAAAAATTAAAAAAAAACATATAAACACAAATGATGGTTTTTCCTTCAATTTCGATATCAATTTATAGAAACAAAATATATACTTGTATAATTTTATTTTTTTATATAAATCATTACATATATAATTATACAATATTTTTTCTAAGAGATAATTATATATTAATATATATAAAAAAAGGTGTTTTTTTTTTTTTTTTTTATTTTTATTTTTATTTTATGGTAATATTTTATTTTCCTTATTTTATAAATTATATTAGTTTATATGTGATTAATTTTATATATTATCAATTTATATATTTTTAAATGCTTACTTAATTATCTTTTTTTTTTTTTTTTTTTTTTTTTCCCCTCTTTTTATATTAATTTATTTTTGAAAAAATTGATATATATATAGTTGAAATATAAATTTCAAAAAAAATGATCACAAAATATACACTTAAATATAGGTACAATAAAAAAAAAAAATAAAAATATAATTACAAGATAATATTTTTTCCTGCTATCAAATTTTTATATATTCTCCTCAAAGAAAAAATATAAATAAATGAAGTAAATTAAAAAAAAAATTTTCTTTTTCTTCTTCTTTTGTAATTCCTTATTTATACATATTTTACTATATTTTCATAAAAATAAATTGTCATATTATATAAATATATATACCAAACCATAATTATATAGCCCTCAACATATTTTTATGATGTTTTTTCTTTTTAAATGTGATACGTAATTAAATATAATAATATATATTAAATATTATATTTTGTAATATTTACTTTCATGAGGTTTAATAAATTATAAAGAGAGAATAAAAAAAAAAAAAAAAAAAATTTATCCATATACAAATTATTAATTTATTTTTATTTTTTATTTCCCTTTGTATATATTATAAAAAAATAATCTACATAATTTTATATGATGATAATTACAATATAATATTTATAATATATATTATTTGTTAAGAGAAAAAAAAAATAAATATATACCTTCTTTTTGAGAATTGAATAAATTGTTTAATATATATATATATATATATAAATATATTATACATTTATGGTGAAAAAAAATGTTGTATTTAATTAGATTTAATATATATATATAAAAAATAGCGTATTTAAAATAATATATATATATATATATTATTATTATTACAAAATGACGGATATTATAAAAGTATATATCTATATATATGTATATATATATAATATTATTTTACTATATATATATAATATATAAATGTAAATGCATATAGTATCTATGTATATTATATATATATATAATATTAATTACATATCTAAGTTTTCTTTTCTTTTCTTTTTTTTTTTTTTTTTTTTTATTTTTTATAGAGAGCCCGTTATATATTATATATTAAAATTTTTTATAATAGCATTTATATACAATATTTTATTAACTAAAAGAAAAAAAAAAAAAAAAAAAAAAAAAAAACGAGGAAATTTATATTTCTTTAACAACATTTTAATTAAAATCATGATAATACAATTTCAAATCATTTTGTAATTATATAAATAAATATATATATATATATATATATATATATGTACTTTTAAATTAGGAATATTCTCATTTATAAATATATCTTATTTTTTAAATTGGTATAAAAAAAAAAAAAAAAATAAGAAACCGTTGATTAAATAATACATATATAATATAAATATATTTTATAAATATATATTTATATATATATATATATATATTTATAACGTATATCATTTTAAAGATAAACTAGTATGGCACCAAAAAAAAAAAGAAAAGTTACGCGTGATCCAACAAGAAGTGCAAATAGTGGAGCAGGAGCAGGAGCAGGAGCAATATTAAGTAGAGCTAGCATGGCTTCTAGAATCCTCTGGCATGAGATGTGGCATGAAGGCCTGGAAGAGGCATCTCGTTTGTACTTTGGGGAAAGGAACGTGAAAGGCATGTTTGAGGTGCTGGAGCCCTTGCATGCTATGATGGAACGGGGCCCCCAGACTCTGAAGGAAACATCCTTTAATCAGGCCTATGGTCGAGATTTAATGGAGGCCCAAGAGTGGTGCAGGAAGTACATGAAATCAGGGAATGTCAAGGACCTCCTCCAAGCCTGGGACCTCTATTATCATGTGTTCCGACGAATCTCAAAGGGTGAGGGTCGTGGTTCACTTCTTACTTGCGGTGACGTTGAGGAGAACCCTGGTCCTGTCGACATGACAGCCAGTTTAACTACCAAGTTCTTGAACAATACCTATGAAAACCCATTTATGAATGCATCCGGTGTTCATTGCATGACTACACAAGAATTAGATGAATTAGCAAACTCTAAAGCTGGCGCATTCATTACAAAGAGTGCTACAACCTTAGAAAGAGAAGGTAACCCTGAACCACGTTACATTTCTGTCCCTCTAGGCAGTATCAACTCCATGGGTTTACCAAACGAAGGTATCGACTACTATTTGTCCTATGTATTAAACCGTCAAAAGAATTATCCTGATGCACCTGCTATTTTCTTCTCAGTTGCTGGTATGAGCATTGATGAAAATTTAAATTTGTTGAGGAAAATCCAAGATAGCGAATTCAACGGTATTACCGAGTTAAACTTGTCTTGTCCTAATGTGCCTGGGAAACCACAAGTTGCTTATGACTTTGACTTGACAAAGGAAACCTTGGAAAAGGTTTTTGCCTTTTTCAAAAAACCTCTTGGTGTCAAGTTGCCTCCTTATTTTGATTTTGCCCATTTTGATATCATGGCAAAAATATTGAACGAGTTCCCATTAGCTTATGTCAACTCTATCAATAGTATAGGAAATGGTCTTTTCATTGATGTGGAGAAGGAGAGTGTAGTAGTGAAGCCAAAGAATGGTTTCGGGGGTATTGGAGGTGAATATGTTAAGCCAACCGCGCTCGCCAATGTTCGTGCATTTTACACTCGTTTGAGACCTGAAATCAAAGTTATCGGTACAGGTGGAATTAAGTCCGGTAAGGATGCATTTGAACATCTTCTATGTGGTGCCTCTATGCTACAGATTGGTACAGAATTACAAAAAGAGGGCGTCAAGATTTTTGAACGTATCGAAAAAGAATTAAAAGACATAATGGAAGCTAAGGGTTATACATCCATAGATCAGTTCCGTGGGAAGTTGAACAGCATTTAAAAGCTTATTTAATAATAGATTAAAAATATTATAAAAATAAAAACATAAACACAGAAATTACAAAAAAAATACATATGAATTTTTTTTTTGTAATCTTCCTTATAAATATAGAATAATGAATCATATAAAACATATCATTATTCATTTATTTACATTTAAAATTATTGTTTCAGTATCTTTAATTTATTATGTATATATAAAAATAACTTACAATTTTATTAATAAACAATATATGTTTATTAATTCATGTTTTGTAATTTATGGGATAGCGATTTTTTTTACTGTCTGTATTTTTCTTTTTTAATTATGTTTTAATTGTATTTTATTTTTATTATTGTTCTTTTTATAGTATTATTTTAAAACAAAATGTATTTTCTAAGAACTTATAATAATAATAAATATAAATTTTAATAAAAATTATATTTATCTTTTACAATATGAACATAAAGTACAACATTAATATATAGCTTTTAATATTTTTATTCCTAATCATGTAAATCTTAAATTTTTCTTTTTAAACATATGTTAAATATTTATTTCTCATTATATATAAGAACATATTTATTAAATCTAGAATTCTATAGTGAGTCGTATTACAATTCACTGGCCGTCGTTTTACAACGTCGTGACTGGGAAAACCCTGGCGTTACCCAACTTAATCGCCTTGCAGCACATCCCCCTTTCGCCAGCTGGCGTAATAGCGAAGAGGCCCGCACCGATCGCCCTTCCCAACAGTTGCGCAGCCTGAATGGCGAATGGCGCCTGATGCGGTATTTTCTCCTTACGCATCTGTGCGGTATTTCACACCGCATATGGTGCACTCTCAGTACAATCTGCTCTGATGCCGCATAGTTAAGCCAGCCCCGACACCCGCCAACACCCGCTGACGCGCCCTGACGGGCTTGTCTGCTCCCGGCATCCGCTTACAGACAAGCTGTGACCGTCTCCGGGAGCTGCATGTGTCAGAGGTTTTCACCGTCATCACCGAAACGCGCGAGACGAAAGGGCCTCGTGATACGCCTATTTTTATAGGTTAATGTCATGATAATAATGGTTTCTTAGACGTCAGGTGGCACTTTTCGGGGAAATGTGCGCGGAACCCCTATTTGTTTATTTTTCTAAATACATTCAAATATGTATCCGCTCATGAGACAATAACCCTGATAAATGCTTCAATAATATTGAAAAAGGAAGAGTATGAGTATTCAACATTTCCGTGTCGCCCTTATTCCCTTTTTTGCGGCATTTTGCCTTCCTGTTTTTGCTCACCCAGAAACGCTGGTGAAAGTAAAAGATGCTGAAGATCAGTTGGGTGCACGAGTGGGTTACATCGAACTGGATCTCAACAGCGGTAAGATCCTTGAGAGTTTTCGCCCCGAAGAACGTTTTCCAATGATGAGCACTTTTAAAGTTCTGCTATGTGGCGCGGTATTATCCCGTATTGACGCCGGGCAAGAGCAACTCGGTCGCCGCATACACTATTCTCAGAATGACTTGGTTGAGTACTCACCAGTCACAGAAAAGCATCTTACGGATGGCATGACAGTAAGAGAATTATGCAGTGCTGCCATAACCATGAGTGATAACACTGCGGCCAACTTACTTCTGACAACGATCGGAGGACCGAAGGAGCTAACCGCTTTTTTGCACAACATGGGGGATCATGTAACTCGCCTTGATCGTTGGGAACCGGAGCTGAATGAAGCCATACCAAACGACGAGCGTGACACCACGATGCCTGTAGCAATGCCAACAACGTTGCGCAAACTATTAACTGGCGAACTACTTACTCTAGCTTCCCGGCAACAATTAATAGACTGGATGGAGGCGGATAAAGTTGCAGGACCACTTCTGCGCTCGGCCCTTCCGGCTGGCTGGTTTATTGCTGATAAATCTGGAGCCGGTGAGCGTGGGTCTCGCGGTATCATTGCAGCACTGGGGCCAGATGGTAAGCCCTCCCGTATCGTAGTTATCTACACGACGGGGAGTCAGGCAACTATGGATGAACGAAATAGACAGATCGCTGAGATAGGTGCCTCACTGATTAAGCATTGGTAACTGTCAGACCAAGTTTACTCATATATACTTTAGATTGATTTAAAACTTCATTTTTAATTTAAAAGGATCTAGGTGAAGATCCTTTTTGATAATCTCATGACCAAAATCCCTTAACGTGAGTTTTCGTTCCACTGAGCGTCAGACCCCGTAGAAAAGATCAAAGGATCTTCTTGAGATCCTTTTTTTCTGCGCGTAATCTGCTGCTTGCAAACAAAAAAACCACCGCTACCAGCGGTGGTTTGTTTGCCGGATCAAGAGCTACCAACTCTTTTTCCGAAGGTAACTGGCTTCAGCAGAGCGCAGATACCAAATACTGTCCTTCTAGTGTAGCCGTAGTTAGGCCACCACTTCAAGAACTCTGTAGCACCGCCTACATACCTCGCTCTGCTAATCCTGTTACCAGTGGCTGCTGCCAGTGGCGATAAGTCGTGTCTTACCGGGTTGGACTCAAGACGATAGTTACCGGATAAGGCGCAGCGGTCGGGCTGAACGGGGGGTTCGTGCACACAGCCCAGCTTGGAGCGAACGACCTACACCGAACTGAGATACCTACAGCGTGAGCTATGAGAAAGCGCCACGCTTCCCGAAGGGAGAAAGGCGGACAGGTATCCGGTAAGCGGCAGGGTCGGAACAGGAGAGCGCACGAGGGAGCTTCCAGGGGGAAACGCCTGGTATCTTTATAGTCCTGTCGGGTTTCGCCACCTCTGACTTGAGCGTCGATTTTTGTGATGCTCGTCAGGGGGGCGGAGCCTATCGAAAAACGCCAGCAACGCGGCCTTTTTACGGTTCCTGGCCTTTTGCTGGCCTTTTGCTCACATGTTCTTTCCTGCGTTATCCCCTGATTCTGTGGATAACCGTATTACCGCCTTTGAGTGAGCTGATACCGCTCGCCGCAGCCGAACGACCGAGCGCAGCGAGTCAGTGAGCGAGGAAGCGGAAGAGCGCCCAATACGCAAACCGCCTCTCCCCGCGCGTTGGCCGATTCATTAATGCAGCTGGCACGACAGGTTTCCCGACTGGAAAGCGGGCAGTGAGCGCAACGCAATTAATGTGAGTTAGCTCACTCATTAGGCACCCCAGGCTTTACACTTTATGCTTCCGGCTCGTATGTTGTGTGGAATTGTGAGCGGATAACAATTTCACACAGGAAACAGCTATGACCATGATTACGCCAAGCTATTTAGGTGACACTATAGAATACTC

**>crt´PMII-Halo_nmd3´1xNLS-FRB-T2A-yDHODH (colour code:** Plasmepsin II, FRB, Halo)

CGGCCGCTAACGTAACAGACTTAGGAGGAGATCTTAGTAGTTGAGTGATTCTATATACATATACATAAATAAATTACATAATATTATAAATTTTTTTTATATTATATTAGAATGTTTACTATATAAAAATAAATATTTTCTTATATATTTTTTTATTTTTTCATATGAAAAAAGAAATTTTTTATATATTTTTTTATTATATTTTATAAATAAAAAAGAACTTATACCTATTATTATTATTATATATAAAAAATATATATTTTTTTATAAATTCCTATTTTTCCGATTATTTATTTATTTTTTTTTTTTATTGTTTAAAAATATATAAAAAAAATTCTTATATATTTAATATTATAAGTCCATAAATATATATAATTTATATAATTTATATATTTCCATACTTTTTTTATTATATATCATGAGATAAAATAAATTTCATATAGAAATCGTATTTATTTATTAATTTGATATATATTAATAAATAAATATTAATAATAATATGTTATATATATATATATTTATTTAATTATTATATAGGAAACTATATATATATATATATATTATATTTTTTTTTTTGAAATATAATAATATATAATATTATTCCTAAAAATATCTATATTCTTATACCTGAACCCTTTTTTTTTTTTTTTTTTTTTTTTGACTTTCGATTGTTCATTCTGTTTATTGTATAAATATATAAATATATATATATTGTATATTTTATTACATATATTTTTTTTTTTTGAAAGTTAAGAATTTTAACTTAATAAAAAAAGTACATATATTTTTAATAAATGTCCTCCATTATATAAATTGTTATATAAAAGATTTTATATAATTTAAATAGAATTCATTTATAAACAAATTTGTTTATAAAAATATATTTATGTATATATAATATAAATATATATATTTATATATATATATATATATATATTACTATATATATTTTTTTTTTTTTTTCCTTTTTTTTACTTTCCCAAGTTGTACTGCTTCTAAGCTTTTTTAATAAACATATATAATTTGTACAAATATTTTAGATTATATACATGATGTATATTTGAATATATTTTCTATATATTTGTGGTTCCATTTTTGTATATTATATATAATATATTTATATATATATTGATATGTCAATATTTGTATAACACATGAAGTTTTTGTTTTTTTTTTTTTTTTTTTTTAATGGAGAATATTTAATAATATATGAAAAAAATTTTATATAAATATATATATATATATATATATATATATATATATATATATATATATGTATATATATATATTTATATATACATGTATGTTTTTTAAAAAGTTAAATAATTCTATAGATTATTTTCATTGTCTTCACATATATGACATAAATATTTTAAAATCGACATTCCGATATATTATATTTTTAGACTATAATATCCGTTAATAATAAATACACGCAGTCATATTATTTATTATACATTCATTTATTATTTTGTTTTTTTTAATTTCTTACATATAACTCGAGATGGATATTACAGTAAGAGAACATGATTTTAAACATGGCTTTATCAAAAGCAATTCAACATTTGATGGATTAAACATTGACAATTCAAAGAACAAAAAAAAAATACAGAAAGGATTTCAAATACTATATGTACTTCTCTTTTGTAGTGTAATGTGTGGTTTATTTTATTATGTGTATGAAAATGTATGGCTTCAAAGAGATAATGAAATGAATGAAATTTTAAAAAATTCCGAACATTTAACTATTGGATTTAAAGTTGAAAATGCACATGATAGAATTTTGAAAACTATAAAAACACATAAATTAAAAAATTACATTAAAGAATCTGTCAATTTTCTTAATTCAGGACTTACTAAAACAAATTATTTAGGTAGTTCAAATGATAATATCGAATTAGTAGATTTCCAAAATATAATGTTTTATGGTGATGCAGAAGTTGGAGATAACCAACAACCATTTACATTTATTCTTGATACAGGATCTGCTAATTTATGGGTCCCAAGTGTTAAATGTACAACTGCAGGATGTTTAACTAAACATCTATATGATTCATCTAAATCAAGAACATATGAAAAAGATGGAACCAAAGTAGAAATGAATTATGTGTCAGGAACTGTTAGTGGATTTTTCAGTAAAGATTTAGTAACTGTTGGTAATTTATCTCTTCCATATAAATTTATTGAAGTAATAGATACTAATGGATTCGAACCAACTTATACTGCTTCAACATTTGATGGTATCCTTGGTTTAGGATGGAAAGATTTATCAATAGGTTCAGTAGATCCAATTGTTGTTGAATTAAAAAACCAAAACAAAATTGAAAATGCTCTTTTCACCTTTTACTTACCTGTACATGATAAACATACAGGATTCTTAACCATTGGTGGTATTGAAGAAAGATTTTATGAAGGACCACTAACTTACGAAAAATTAAACCACGATTTATATTGGCAAATAACTTTAGATGCACACGTTGGAAATATAATGTTAGAAAAAGCAAACTGTATTGTAGATAGTGGTACTAGTGCCATTACTGTACCAACTGACTTTTTAAATAAAATGTTGCAGAATTTAGATGTTATCAAAGTCCCATTCTTACCTTTCTATGTAACTCTTTGTAACAACAGCAAATTACCAACTTTTGAATTTACCTCAGAAAATGGTAAATACACATTAGAACCTGAATACTACCTTCAACACATAGAAGATGTTGGTCCAGGATTATGTATGCTTAATATCATAGGATTAGATTTTCCAGTACCAACCTTTATTCTAGGTGACCCATTCATGAGAAAATATTTTACCGTCTTTGATTATGATAATCAGAGTGTTGGTATTGCTCTTGCTAAAAAGAATTTACCTAGGACAAGTTATCCATATGATAATCCAGATTATGCACCAGTTGCAACATTAGGTACCATGGCAGAAATTGGTACGGGTTTTCCATTTGATCCTCATTATGTGGAGGTGCTGGGGGAAAGGATGCATTATGTTGACGTAGGACCAAGAGATGGTACTCCAGTGTTATTTTTGCATGGAAACCCAACCTCGAGTTATGTATGGAGAAATATAATTCCACATGTAGCACCAACACATAGATGTATAGCTCCTGATTTAATTGGTATGGGAAAAAGTGATAAACCTGACTTAGGATATTTTTTTGATGATCATGTCCGTTTTATGGATGCTTTCATTGAAGCTCTTGGCCTTGAAGAAGTAGTATTAGTTATACATGATTGGGGATCCGCCTTAGGATTTCATTGGGCCAAGAGGAATCCTGAAAGAGTAAAAGGAATAGCATTCATGGAATTCATACGACCAATCCCCACATGGGATGAATGGCCAGAATTTGCACGCGAAACATTTCAAGCTTTTAGAACTACAGATGTTGGTAGAAAATTAATAATAGATCAAAATGTATTTATAGAAGGAACTTTACCTATGGGTGTTGTAAGGCCGTTAACAGAAGTTGAAATGGACCACTACCGTGAACCTTTTTTAAATCCAGTAGATAGAGAGCCCTTATGGAGATTTCCTAATGAATTACCTATTGCAGGTGAACCCGCGAATATTGTTGCTTTAGTAGAAGAATATATGGATTGGTTACATCAGTCTCCTGTTCCTAAACTTCTATTTTGGGGTACACCTGGAGTTCTAATACCACCAGCTGAAGCAGCAAGATTAGCAAAATCATTACCAAATTGTAAAGCTGTTGATATAGGTCCTGGGTTGAATTTATTACAAGAAGATAATCCAGATTTGATTGGATCTGAGATAGCTAGATGGCTAAGTACATTAGAAATTTCAGGTTAACCCGGGTCGAGGGATATGGCAGCTTAATGTTCGTTTTTCTTATTTATATATTTATACCAATTGATTGTATTTATAACTGTAAAAATGTGTATGTTGTGTGCATATTTTTTTTTGTGCATGCACATGCATGTAAATAGCTAAAATTATGAACATTTTATTTTTTGTTCAGAAAAAAAAAACTTTACACACATAAAATGGCTAGTATGAATAGCCATATTTTATATAAATTAAATCCTATGAATTTATGACCATATTAAAAATTTAGATATTTATGGAACATAATATGTTTGAAACAATAAGACAAAATTATTATTATTATTATTATTTTTACTGTTATAATTATGTGTCTCCTTCAATGATTCATAAATAGTTGGACTTGATTTTTAAAATGTTTATAATATGATTAGCATAGTTAAATAAAAAAAGTTGAAAAATTAAAAAAAAACATATAAACACAAATGATGGTTTTTCCTTCAATTTCGATATCAATTTATAGAAACAAAATATATACTTGTATAATTTTATTTTTTTATATAAATCATTACATATATAATTATACAATATTTTTTCTAAGAGATAATTATATATTAATATATATAAAAAAAGGTGTTTTTTTTTTTTTTTTTTATTTTTATTTTTATTTTATGGTAATATTTTATTTTCCTTATTTTATAAATTATATTAGTTTATATGTGATTAATTTTATATATTATCAATTTATATATTTTTAAATGCTTACTTAATTATCTTTTTTTTTTTTTTTTTTTTTTTTTCCCCTCTTTTTATATTAATTTATTTTTGAAAAAATTGATATATATATAGTTGAAATATAAATTTCAAAAAAAATGATCACAAAATATACACTTAAATATAGGTACAATAAAAAAAAAAAATAAAAATATAATTACAAGATAATATTTTTTCCTGCTATCAAATTTTTATATATTCTCCTCAAAGAAAAAATATAAATAAATGAAGTAAATTAAAAAAAAAATTTTCTTTTTCTTCTTCTTTTGTAATTCCTTATTTATACATATTTTACTATATTTTCATAAAAATAAATTGTCATATTATATAAATATATATACCAAACCATAATTATATAGCCCTCAACATATTTTTATGATGTTTTTTCTTTTTAAATGTGATACGTAATTAAATATAATAATATATATTAAATATTATATTTTGTAATATTTACTTTCATGAGGTTTAATAAATTATAAAGAGAGAATAAAAAAAAAAAAAAAAAAAATTTATCCATATACAAATTATTAATTTATTTTTATTTTTTATTTCCCTTTGTATATATTATAAAAAAATAATCTACATAATTTTATATGATGATAATTACAATATAATATTTATAATATATATTATTTGTTAAGAGAAAAAAAAAATAAATATATACCTTCTTTTTGAGAATTGAATAAATTGTTTAATATATATATATATATATATAAATATATTATACATTTATGGTGAAAAAAAATGTTGTATTTAATTAGATTTAATATATATATATAAAAAATAGCGTATTTAAAATAATATATATATATATATATTATTATTATTACAAAATGACGGATATTATAAAAGTATATATCTATATATATGTATATATATATAATATTATTTTACTATATATATATAATATATAAATGTAAATGCATATAGTATCTATGTATATTATATATATATATAATATTAATTACATATCTAAGTTTTCTTTTCTTTTCTTTTTTTTTTTTTTTTTTTTTATTTTTTATAGAGAGCCCGTTATATATTATATATTAAAATTTTTTATAATAGCATTTATATACAATATTTTATTAACTAAAAGAAAAAAAAAAAAAAAAAAAAAAAAAAAACGAGGAAATTTATATTTCTTTAACAACATTTTAATTAAAATCATGATAATACAATTTCAAATCATTTTGTAATTATATAAATAAATATATATATATATATATATATATATATGTACTTTTAAATTAGGAATATTCTCATTTATAAATATATCTTATTTTTTAAATTGGTATAAAAAAAAAAAAAAAAATAAGAAACCGTTGATTAAATAATACATATATAATATAAATATATTTTATAAATATATATTTATATATATATATATATATATTTATAACGTATATCATTTTAAAGATAAACTAGTATGGCACCAAAAAAAAAAAGAAAAGTTACGCGTGATCCAACAAGAAGTGCAAATAGTGGAGCAGGAGCAGGAGCAGGAGCAATATTAAGTAGAGCTAGCATGGCTTCTAGAATCCTCTGGCATGAGATGTGGCATGAAGGCCTGGAAGAGGCATCTCGTTTGTACTTTGGGGAAAGGAACGTGAAAGGCATGTTTGAGGTGCTGGAGCCCTTGCATGCTATGATGGAACGGGGCCCCCAGACTCTGAAGGAAACATCCTTTAATCAGGCCTATGGTCGAGATTTAATGGAGGCCCAAGAGTGGTGCAGGAAGTACATGAAATCAGGGAATGTCAAGGACCTCCTCCAAGCCTGGGACCTCTATTATCATGTGTTCCGACGAATCTCAAAGGGTGAGGGTCGTGGTTCACTTCTTACTTGCGGTGACGTTGAGGAGAACCCTGGTCCTGTCGACATGACAGCCAGTTTAACTACCAAGTTCTTGAACAATACCTATGAAAACCCATTTATGAATGCATCCGGTGTTCATTGCATGACTACACAAGAATTAGATGAATTAGCAAACTCTAAAGCTGGCGCATTCATTACAAAGAGTGCTACAACCTTAGAAAGAGAAGGTAACCCTGAACCACGTTACATTTCTGTCCCTCTAGGCAGTATCAACTCCATGGGTTTACCAAACGAAGGTATCGACTACTATTTGTCCTATGTATTAAACCGTCAAAAGAATTATCCTGATGCACCTGCTATTTTCTTCTCAGTTGCTGGTATGAGCATTGATGAAAATTTAAATTTGTTGAGGAAAATCCAAGATAGCGAATTCAACGGTATTACCGAGTTAAACTTGTCTTGTCCTAATGTGCCTGGGAAACCACAAGTTGCTTATGACTTTGACTTGACAAAGGAAACCTTGGAAAAGGTTTTTGCCTTTTTCAAAAAACCTCTTGGTGTCAAGTTGCCTCCTTATTTTGATTTTGCCCATTTTGATATCATGGCAAAAATATTGAACGAGTTCCCATTAGCTTATGTCAACTCTATCAATAGTATAGGAAATGGTCTTTTCATTGATGTGGAGAAGGAGAGTGTAGTAGTGAAGCCAAAGAATGGTTTCGGGGGTATTGGAGGTGAATATGTTAAGCCAACCGCGCTCGCCAATGTTCGTGCATTTTACACTCGTTTGAGACCTGAAATCAAAGTTATCGGTACAGGTGGAATTAAGTCCGGTAAGGATGCATTTGAACATCTTCTATGTGGTGCCTCTATGCTACAGATTGGTACAGAATTACAAAAAGAGGGCGTCAAGATTTTTGAACGTATCGAAAAAGAATTAAAAGACATAATGGAAGCTAAGGGTTATACATCCATAGATCAGTTCCGTGGGAAGTTGAACAGCATTTAAAAGCTTATTTAATAATAGATTAAAAATATTATAAAAATAAAAACATAAACACAGAAATTACAAAAAAAATACATATGAATTTTTTTTTTGTAATCTTCCTTATAAATATAGAATAATGAATCATATAAAACATATCATTATTCATTTATTTACATTTAAAATTATTGTTTCAGTATCTTTAATTTATTATGTATATATAAAAATAACTTACAATTTTATTAATAAACAATATATGTTTATTAATTCATGTTTTGTAATTTATGGGATAGCGATTTTTTTTACTGTCTGTATTTTTCTTTTTTAATTATGTTTTAATTGTATTTTATTTTTATTATTGTTCTTTTTATAGTATTATTTTAAAACAAAATGTATTTTCTAAGAACTTATAATAATAATAAATATAAATTTTAATAAAAATTATATTTATCTTTTACAATATGAACATAAAGTACAACATTAATATATAGCTTTTAATATTTTTATTCCTAATCATGTAAATCTTAAATTTTTCTTTTTAAACATATGTTAAATATTTATTTCTCATTATATATAAGAACATATTTATTAAATCTAGAATTCTATAGTGAGTCGTATTACAATTCACTGGCCGTCGTTTTACAACGTCGTGACTGGGAAAACCCTGGCGTTACCCAACTTAATCGCCTTGCAGCACATCCCCCTTTCGCCAGCTGGCGTAATAGCGAAGAGGCCCGCACCGATCGCCCTTCCCAACAGTTGCGCAGCCTGAATGGCGAATGGCGCCTGATGCGGTATTTTCTCCTTACGCATCTGTGCGGTATTTCACACCGCATATGGTGCACTCTCAGTACAATCTGCTCTGATGCCGCATAGTTAAGCCAGCCCCGACACCCGCCAACACCCGCTGACGCGCCCTGACGGGCTTGTCTGCTCCCGGCATCCGCTTACAGACAAGCTGTGACCGTCTCCGGGAGCTGCATGTGTCAGAGGTTTTCACCGTCATCACCGAAACGCGCGAGACGAAAGGGCCTCGTGATACGCCTATTTTTATAGGTTAATGTCATGATAATAATGGTTTCTTAGACGTCAGGTGGCACTTTTCGGGGAAATGTGCGCGGAACCCCTATTTGTTTATTTTTCTAAATACATTCAAATATGTATCCGCTCATGAGACAATAACCCTGATAAATGCTTCAATAATATTGAAAAAGGAAGAGTATGAGTATTCAACATTTCCGTGTCGCCCTTATTCCCTTTTTTGCGGCATTTTGCCTTCCTGTTTTTGCTCACCCAGAAACGCTGGTGAAAGTAAAAGATGCTGAAGATCAGTTGGGTGCACGAGTGGGTTACATCGAACTGGATCTCAACAGCGGTAAGATCCTTGAGAGTTTTCGCCCCGAAGAACGTTTTCCAATGATGAGCACTTTTAAAGTTCTGCTATGTGGCGCGGTATTATCCCGTATTGACGCCGGGCAAGAGCAACTCGGTCGCCGCATACACTATTCTCAGAATGACTTGGTTGAGTACTCACCAGTCACAGAAAAGCATCTTACGGATGGCATGACAGTAAGAGAATTATGCAGTGCTGCCATAACCATGAGTGATAACACTGCGGCCAACTTACTTCTGACAACGATCGGAGGACCGAAGGAGCTAACCGCTTTTTTGCACAACATGGGGGATCATGTAACTCGCCTTGATCGTTGGGAACCGGAGCTGAATGAAGCCATACCAAACGACGAGCGTGACACCACGATGCCTGTAGCAATGCCAACAACGTTGCGCAAACTATTAACTGGCGAACTACTTACTCTAGCTTCCCGGCAACAATTAATAGACTGGATGGAGGCGGATAAAGTTGCAGGACCACTTCTGCGCTCGGCCCTTCCGGCTGGCTGGTTTATTGCTGATAAATCTGGAGCCGGTGAGCGTGGGTCTCGCGGTATCATTGCAGCACTGGGGCCAGATGGTAAGCCCTCCCGTATCGTAGTTATCTACACGACGGGGAGTCAGGCAACTATGGATGAACGAAATAGACAGATCGCTGAGATAGGTGCCTCACTGATTAAGCATTGGTAACTGTCAGACCAAGTTTACTCATATATACTTTAGATTGATTTAAAACTTCATTTTTAATTTAAAAGGATCTAGGTGAAGATCCTTTTTGATAATCTCATGACCAAAATCCCTTAACGTGAGTTTTCGTTCCACTGAGCGTCAGACCCCGTAGAAAAGATCAAAGGATCTTCTTGAGATCCTTTTTTTCTGCGCGTAATCTGCTGCTTGCAAACAAAAAAACCACCGCTACCAGCGGTGGTTTGTTTGCCGGATCAAGAGCTACCAACTCTTTTTCCGAAGGTAACTGGCTTCAGCAGAGCGCAGATACCAAATACTGTCCTTCTAGTGTAGCCGTAGTTAGGCCACCACTTCAAGAACTCTGTAGCACCGCCTACATACCTCGCTCTGCTAATCCTGTTACCAGTGGCTGCTGCCAGTGGCGATAAGTCGTGTCTTACCGGGTTGGACTCAAGACGATAGTTACCGGATAAGGCGCAGCGGTCGGGCTGAACGGGGGGTTCGTGCACACAGCCCAGCTTGGAGCGAACGACCTACACCGAACTGAGATACCTACAGCGTGAGCTATGAGAAAGCGCCACGCTTCCCGAAGGGAGAAAGGCGGACAGGTATCCGGTAAGCGGCAGGGTCGGAACAGGAGAGCGCACGAGGGAGCTTCCAGGGGGAAACGCCTGGTATCTTTATAGTCCTGTCGGGTTTCGCCACCTCTGACTTGAGCGTCGATTTTTGTGATGCTCGTCAGGGGGGCGGAGCCTATCGAAAAACGCCAGCAACGCGGCCTTTTTACGGTTCCTGGCCTTTTGCTGGCCTTTTGCTCACATGTTCTTTCCTGCGTTATCCCCTGATTCTGTGGATAACCGTATTACCGCCTTTGAGTGAGCTGATACCGCTCGCCGCAGCCGAACGACCGAGCGCAGCGAGTCAGTGAGCGAGGAAGCGGAAGAGCGCCCAATACGCAAACCGCCTCTCCCCGCGCGTTGGCCGATTCATTAATGCAGCTGGCACGACAGGTTTCCCGACTGGAAAGCGGGCAGTGAGCGCAACGCAATTAATGTGAGTTAGCTCACTCATTAGGCACCCCAGGCTTTACACTTTATGCTTCCGGCTCGTATGTTGTGTGGAATTGTGAGCGGATAACAATTTCACACAGGAAACAGCTATGACCATGATTACGCCAAGCTATTTAGGTGACACTATAGAATACTC

**>crt-mScarlet-Rab11a_nmd3-1xNLS-FRB-T2A DHODH**

crt promotor mScarlet linker Rab11a recodonized (no introns) nmd3 promotor NLS·FRB T2A (skip peptide) yDHODH (DSM1) *amp* promotor+Ampiclin resitance replication origin

cggccgctaacgtaacagacttaggaggagatcttagtagttgagtgattctatatacatatacataaataaattacataatattataaattttttttatattatattagaatgtttactatataaaaataaatattttcttatatatttttttattttttcatatgaaaaaagaaattttttatatatttttttattatattttataaataaaaaagaacttatacctattattattattatatataaaaaatatatatttttttataaattcctatttttccgattatttatttattttttttttttattgtttaaaaatatataaaaaaaattcttatatatttaatattataagtccataaatatatataatttatataatttatatatttccatactttttttattatatatcatgagataaaataaatttcatatagaaatcgtatttatttattaatttgatatatattaataaataaatattaataataatatgttatatatatatatatttatttaattattatataggaaactatatatatatatatatattatatttttttttttgaaatataataatatataatattattcctaaaaatatctatattcttatacctgaacccttttttttttttttttttttttttgactttcgattgttcattctgtttattgtataaatatataaatatatatatattgtatattttattacatatattttttttttttgaaagttaagaattttaacttaataaaaaaagtacatatatttttaataaatgtcctccattatataaattgttatataaaagattttatataatttaaatagaattcatttataaacaaatttgtttataaaaatatatttatgtatatataatataaatatatatatttatatatatatatatatatatattactatatatatttttttttttttttccttttttttactttcccaagttgtactgcttctaagcttttttaataaacatatataatttgtacaaatattttagattatatacatgatgtatatttgaatatattttctatatatttgtggttccatttttgtatattatatataatatatttatatatatattgatatgtcaatatttgtataacacatgaagtttttgtttttttttttttttttttttaatggagaatatttaataatatatgaaaaaaattttatataaatatatatatatatatatatatatatatatatatatatatatatatgtatatatatatatttatatatacatgtatgttttttaaaaagttaaataattctatagattattttcattgtcttcacatatatgacataaatattttaaaatcgacattccgatatattatatttttagactataatatccgttaataataaatacacgcagtcatattatttattatacattcatttattattttgttttttttaatttcttacatataactcgagATGGTGAGTAAGGGTGAGGCAGTGATTAAGGAGTTTATGCGTTTTAAGGTGCACATGGAGGGTAGTATGAACGGTCACGAGTTTGAGATTGAGGGTGAGGGTGAGGGTCGTCCATACGAGGGTACACAGACAGCAAAGCTTAAGGTGACAAAGGGTGGTCCACTTCCATTTAGTTGGGATATTCTTAGTCCACAGTTTATGTACGGTAGTCGTGCATTTACAAAGCACCCAGCAGATATTCCAGATTACTACAAGCAGAGTTTTCCAGAGGGTTTTAAGTGGGAGCGTGTGATGAACTTTGAGGATGGTGGTGCAGTGACAGTGACACAGGATACAAGTCTTGAGGATGGTACACTTATTTACAAGGTGAAGCTTCGTGGTACAAACTTTCCACCAGATGGTCCAGTGATGCAGAAGAAGACAATGGGTTGGGAGGCAAGTACAGAGCGTCTTTACCCAGAGGATGGTGTGCTTAAGGGTGATATTAAGATGGCACTTCGTCTTAAGGATGGTGGTCGTTACCTTGCAGATTTTAAGACAACATACAAGGCAAAGAAGCCAGTGCAGATGCCAGGTGCATACAACGTGGATCGTAAGCTTGATATTACAAGTCACAACGAGGATTACACAGTGGTGGAGCAGTACGAGCGTAGTGAGGGTCGTCACAGTACAGGTGGTATGGATGAGCTTTACAAGCCTAGGACAAGTTATCCATATGATAATCCAGATTATGCACCAGTTGCAACATTAGGTACCGCTATGAAGGAGGACTACTACGACTATTTATTCAAGATAGTATTAATAGGTGACAGTGGTGTTGGTAAGAGTAATTTGTTAAGTCGTTTCACTCGTGATGAGTTCAACCTTGAGTCAAAATCAACAATTGGAGTTGAGTTCGCAACAAAGTCAATACAGCTTAAGAACAACAAGATAATTAAGGCACAGATTTGGGACACTGCTGGACAGGAGAGGTACCGTGCTATAACAAGCGCATACTACAGGGGTGCTGTAGGAGCACTTCTTGTTTACGACATTACTAAGAAGAACAGTTTCGAGAACATAGAGAAGTGGCTTAAGGAGCTTCGTGACAACGCTGACAGTAACATAGTTATACTTCTTGTAGGAAACAAGTCAGACCTTAAACACCTTAGAGTAATAAACGACAACGACGCAACTCAGTACGCAAAGAAGGAGAAATTAGCTTTTATTGAGACTAGCGCATTAGAAGCTACAAACGTTGAATTGGCATTCCACCAGCTTCTTAACGAGATATACAACGTTCGTCAGAAGAAGCAGGCAACTAAGAACGACGACAACCTTAGTATTCAGCCACGTGGAAAGAAGATAAACGTTGACGACGACAACGACAAGAATGAGACAAAGAAGAAGAATAAGTGCTGCTAGcccgggtcgagggatatggcagcttaatgttcgtttttcttatttatatatttataccaattgattgtatttataactgtaaaaatgtgtatgttgtgtgcatatttttttttgtgcatgcacatgcatgtaaatagctaaaattatgaacattttattttttgttcagaaaaaaaaaactttacacacataaaatggctagtatgaatagccatattttatataaattaaatcctatgaatttatgaccatattaaaaatttagatatttatggaacataatatgtttgaaacaataagacaaaattattattattattattatttttactgttataattatgtgtctccttcaatgattcataaatagttggacttgatttttaaaatgtttataatatgattagcatagttaaataaaaaaagttgaaaaattaaaaaaaaacatataaacacaaatgatggtttttccttcaatttcgatatcaatttatagaaacaaaatatatacttgtataattttatttttttatataaatcattacatatataattatacaatattttttctaagagataattatatattaatatatataaaaaaaggtgttttttttttttttttttatttttatttttattttatggtaatattttattttccttattttataaattatattagtttatatgtgattaattttatatattatcaatttatatatttttaaatgcttacttaattatctttttttttttttttttttttttttcccctctttttatattaatttatttttgaaaaaattgatatatatatagttgaaatataaatttcaaaaaaaatgatcacaaaatatacacttaaatataggtacaataaaaaaaaaaaataaaaatataattacaagataatattttttcctgctatcaaatttttatatattctcctcaaagaaaaaatataaataaatgaagtaaattaaaaaaaaaattttctttttcttcttcttttgtaattccttatttatacatattttactatattttcataaaaataaattgtcatattatataaatatatataccaaaccataattatatagccctcaacatatttttatgatgttttttctttttaaatgtgatacgtaattaaatataataatatatattaaatattatattttgtaatatttactttcatgaggtttaataaattataaagagagaataaaaaaaaaaaaaaaaaaaatttatccatatacaaattattaatttatttttattttttatttccctttgtatatattataaaaaaataatctacataattttatatgatgataattacaatataatatttataatatatattatttgttaagagaaaaaaaaaataaatatataccttctttttgagaattgaataaattgtttaatatatatatatatatatataaatatattatacatttatggtgaaaaaaaatgttgtatttaattagatttaatatatatatataaaaaatagcgtatttaaaataatatatatatatatatattattattattacaaaatgacggatattataaaagtatatatctatatatatgtatatatatataatattattttactatatatatataatatataaatgtaaatgcatatagtatctatgtatattatatatatatataatattaattacatatctaagttttcttttcttttctttttttttttttttttttttattttttatagagagcccgttatatattatatattaaaattttttataatagcatttatatacaatattttattaactaaaagaaaaaaaaaaaaaaaaaaaaaaaaaaaacgaggaaatttatatttctttaacaacattttaattaaaatcatgataatacaatttcaaatcattttgtaattatataaataaatatatatatatatatatatatatatatgtacttttaaattaggaatattctcatttataaatatatcttattttttaaattggtataaaaaaaaaaaaaaaaataagaaaccgttgattaaataatacatatataatataaatatattttataaatatatatttatatatatatatatatatatttataacgtatatcattttaaagataaactagtATGGCACCAAAAAAAAAAAGAAAAGTTACGCGTGATCCAACAAGAAGTGCAAATAGTGGAGCAGGAGCAGGAGCAGGAGCAATATTAAGTAGAGCTAGCATGGCTTCTAGAATCCTCTGGCATGAGATGTGGCATGAAGGCCTGGAAGAGGCATCTCGTTTGTACTTTGGGGAAAGGAACGTGAAAGGCATGTTTGAGGTGCTGGAGCCCTTGCATGCTATGATGGAACGGGGCCCCCAGACTCTGAAGGAAACATCCTTTAATCAGGCCTATGGTCGAGATTTAATGGAGGCCCAAGAGTGGTGCAGGAAGTACATGAAATCAGGGAATGTCAAGGACCTCCTCCAAGCCTGGGACCTCTATTATCATGTGTTCCGACGAATCTCAAAGGGTGAGGGTCGTGGTTCACTTCTTACTTGCGGTGACGTTGAGGAGAACCCTGGTCCTGTCGACATGACAGCCAGTTTAACTACCAAGTTCTTGAACAATACCTATGAAAACCCATTTATGAATGCATCCGGTGTTCATTGCATGACTACACAAGAATTAGATGAATTAGCAAACTCTAAAGCTGGCGCATTCATTACAAAGAGTGCTACAACCTTAGAAAGAGAAGGTAACCCTGAACCACGTTACATTTCTGTCCCTCTAGGCAGTATCAACTCCATGGGTTTACCAAACGAAGGTATCGACTACTATTTGTCCTATGTATTAAACCGTCAAAAGAATTATCCTGATGCACCTGCTATTTTCTTCTCAGTTGCTGGTATGAGCATTGATGAAAATTTAAATTTGTTGAGGAAAATCCAAGATAGCGAATTCAACGGTATTACCGAGTTAAACTTGTCTTGTCCTAATGTGCCTGGGAAACCACAAGTTGCTTATGACTTTGACTTGACAAAGGAAACCTTGGAAAAGGTTTTTGCCTTTTTCAAAAAACCTCTTGGTGTCAAGTTGCCTCCTTATTTTGATTTTGCCCATTTTGATATCATGGCAAAAATATTGAACGAGTTCCCATTAGCTTATGTCAACTCTATCAATAGTATAGGAAATGGTCTTTTCATTGATGTGGAGAAGGAGAGTGTAGTAGTGAAGCCAAAGAATGGTTTCGGGGGTATTGGAGGTGAATATGTTAAGCCAACCGCGCTCGCCAATGTTCGTGCATTTTACACTCGTTTGAGACCTGAAATCAAAGTTATCGGTACAGGTGGAATTAAGTCCGGTAAGGATGCATTTGAACATCTTCTATGTGGTGCCTCTATGCTACAGATTGGTACAGAATTACAAAAAGAGGGCGTCAAGATTTTTGAACGTATCGAAAAAGAATTAAAAGACATAATGGAAGCTAAGGGTTATACATCCATAGATCAGTTCCGTGGGAAGTTGAACAGCATTTAAaagcttatttaataatagattaaaaatattataaaaataaaaacataaacacagaaattacaaaaaaaatacatatgaattttttttttgtaatcttccttataaatatagaataatgaatcatataaaacatatcattattcatttatttacatttaaaattattgtttcagtatctttaatttattatgtatatataaaaataacttacaattttattaataaacaatatatgtttattaattcatgttttgtaatttatgggatagcgattttttttactgtctgtatttttcttttttaattatgttttaattgtattttatttttattattgttctttttatagtattattttaaaacaaaatgtattttctaagaacttataataataataaatataaattttaataaaaattatatttatcttttacaatatgaacataaagtacaacattaatatatagcttttaatatttttattcctaatcatgtaaatcttaaatttttctttttaaacatatgttaaatatttatttctcattatatataagaacatatttattaaatctagaattctatagtgagtcgtattacaattcactggccgtcgttttacaacgtcgtgactgggaaaaccctggcgttacccaacttaatcgccttgcagcacatccccctttcgccagctggcgtaatagcgaagaggcccgcaccgatcgcccttcccaacagttgcgcagcctgaatggcgaatggcgcctgatgcggtattttctccttacgcatctgtgcggtatttcacaccgcatatggtgcactctcagtacaatctgctctgatgccgcatagttaagccagccccgacacccgccaacacccgctgacgcgccctgacgggcttgtctgctcccggcatccgcttacagacaagctgtgaccgtctccgggagctgcatgtgtcagaggttttcaccgtcatcaccgaaacgcgcgagacgaaagggcctcgtgatacgcctatttttataggttaatgtcatgataataatggtttcttagacgtcaggtggcacttttcggggaaatgtgcgcggaacccctatttgtttatttttctaaatacattcaaatatgtatccgctcatgagacaataaccctgataaatgcttcaataatattgaaaaaggaagagtATGAGTATTCAACATTTCCGTGTCGCCCTTATTCCCTTTTTTGCGGCATTTTGCCTTCCTGTTTTTGCTCACCCAGAAACGCTGGTGAAAGTAAAAGATGCTGAAGATCAGTTGGGTGCACGAGTGGGTTACATCGAACTGGATCTCAACAGCGGTAAGATCCTTGAGAGTTTTCGCCCCGAAGAACGTTTTCCAATGATGAGCACTTTTAAAGTTCTGCTATGTGGCGCGGTATTATCCCGTATTGACGCCGGGCAAGAGCAACTCGGTCGCCGCATACACTATTCTCAGAATGACTTGGTTGAGTACTCACCAGTCACAGAAAAGCATCTTACGGATGGCATGACAGTAAGAGAATTATGCAGTGCTGCCATAACCATGAGTGATAACACTGCGGCCAACTTACTTCTGACAACGATCGGAGGACCGAAGGAGCTAACCGCTTTTTTGCACAACATGGGGGATCATGTAACTCGCCTTGATCGTTGGGAACCGGAGCTGAATGAAGCCATACCAAACGACGAGCGTGACACCACGATGCCTGTAGCAATGCCAACAACGTTGCGCAAACTATTAACTGGCGAACTACTTACTCTAGCTTCCCGGCAACAATTAATAGACTGGATGGAGGCGGATAAAGTTGCAGGACCACTTCTGCGCTCGGCCCTTCCGGCTGGCTGGTTTATTGCTGATAAATCTGGAGCCGGTGAGCGTGGGTCTCGCGGTATCATTGCAGCACTGGGGCCAGATGGTAAGCCCTCCCGTATCGTAGTTATCTACACGACGGGGAGTCAGGCAACTATGGATGAACGAAATAGACAGATCGCTGAGATAGGTGCCTCACTGATTAAGCATTGGTAActgtcagaccaagtttactcatatatactttagattgatttaaaacttcatttttaatttaaaaggatctaggtgaagatcctttttgataatctcatgaccaaaatcccttaacgtgagttttcgttccactgagcgtcagaccccgtagaaaagatcaaaggatcttcttgagatcctttttttctgcgcgtaatctgctgcttgcaaacaaaaaaaccaccgctaccagcggtggtttgtttgccggatcaagagctaccaactctttttccgaaggtaactggcttcagcagagcgcagataccaaatactgtccttctagtgtagccgtagttaggccaccacttcaagaactctgtagcaccgcctacatacctcgctctgctaatcctgttaccagtggctgctgccagtggcgataagtcgtgtcttaccgggttggactcaagacgatagttaccggataaggcgcagcggtcgggctgaacggggggttcgtgcacacagcccagcttggagcgaacgacctacaccgaactgagatacctacagcgtgagctatgagaaagcgccacgcttcccgaagggagaaaggcggacaggtatccggtaagcggcagggtcggaacaggagagcgcacgagggagcttccagggggaaacgcctggtatctttatagtcctgtcgggtttcgccacctctgacttgagcgtcgatttttgtgatgctcgtcaggggggcggagcctatcgaaaaacgccagcaacgcggcctttttacggttcctggccttttgctggccttttgctcacatgttctttcctgcgttatcccctgattctgtggataaccgtattaccgcctttgagtgagctgataccgctcgccgcagccgaacgaccgagcgcagcgagtcagtgagcgaggaagcggaagagcgcccaatacgcaaaccgcctctccccgcgcgttggccgattcattaatgcagctggcacgacaggtttcccgactggaaagcgggcagtgagcgcaacgcaattaatgtgagttagctcactcattaggcaccccaggctttacactttatgcttccggctcgtatgttgtgtggaattgtgagcggataacaatttcacacaggaaacagctatgaccatgattacgccaagctatttaggtgacactatagaatactc

**>crt-mScarlet-Rab5a(recod)_nmd3-1xNLS-FRB-T2A**

CGGCCGCTAACGTAACAGACTTAGGAGGAGATCTTAGTAGTTGAGTGATTCTATATACATATACATAAATAAATTACATAATATTATAAATTTTTTTTATATTATATTAGAATGTTTACTATATAAAAATAAATATTTTCTTATATATTTTTTTATTTTTTCATATGAAAAAAGAAATTTTTTATATATTTTTTTATTATATTTTATAAATAAAAAAGAACTTATACCTATTATTATTATTATATATAAAAAATATATATTTTTTTATAAATTCCTATTTTTCCGATTATTTATTTATTTTTTTTTTTTATTGTTTAAAAATATATAAAAAAAATTCTTATATATTTAATATTATAAGTCCATAAATATATATAATTTATATAATTTATATATTTCCATACTTTTTTTATTATATATCATGAGATAAAATAAATTTCATATAGAAATCGTATTTATTTATTAATTTGATATATATTAATAAATAAATATTAATAATAATATGTTATATATATATATATTTATTTAATTATTATATAGGAAACTATATATATATATATATATTATATTTTTTTTTTTGAAATATAATAATATATAATATTATTCCTAAAAATATCTATATTCTTATACCTGAACCCTTTTTTTTTTTTTTTTTTTTTTTTGACTTTCGATTGTTCATTCTGTTTATTGTATAAATATATAAATATATATATATTGTATATTTTATTACATATATTTTTTTTTTTTGAAAGTTAAGAATTTTAACTTAATAAAAAAAGTACATATATTTTTAATAAATGTCCTCCATTATATAAATTGTTATATAAAAGATTTTATATAATTTAAATAGAATTCATTTATAAACAAATTTGTTTATAAAAATATATTTATGTATATATAATATAAATATATATATTTATATATATATATATATATATATTACTATATATATTTTTTTTTTTTTTTCCTTTTTTTTACTTTCCCAAGTTGTACTGCTTCTAAGCTTTTTTAATAAACATATATAATTTGTACAAATATTTTAGATTATATACATGATGTATATTTGAATATATTTTCTATATATTTGTGGTTCCATTTTTGTATATTATATATAATATATTTATATATATATTGATATGTCAATATTTGTATAACACATGAAGTTTTTGTTTTTTTTTTTTTTTTTTTTTAATGGAGAATATTTAATAATATATGAAAAAAATTTTATATAAATATATATATATATATATATATATATATATATATATATATATATATGTATATATATATATTTATATATACATGTATGTTTTTTAAAAAGTTAAATAATTCTATAGATTATTTTCATTGTCTTCACATATATGACATAAATATTTTAAAATCGACATTCCGATATATTATATTTTTAGACTATAATATCCGTTAATAATAAATACACGCAGTCATATTATTTATTATACATTCATTTATTATTTTGTTTTTTTTAATTTCTTACATATAACTCGAGATGGTGAGTAAGGGTGAGGCAGTGATTAAGGAGTTTATGCGTTTTAAGGTGCACATGGAGGGTAGTATGAACGGTCACGAGTTTGAGATTGAGGGTGAGGGTGAGGGTCGTCCATACGAGGGTACACAGACAGCAAAGCTTAAGGTGACAAAGGGTGGTCCACTTCCATTTAGTTGGGATATTCTTAGTCCACAGTTTATGTACGGTAGTCGTGCATTTACAAAGCACCCAGCAGATATTCCAGATTACTACAAGCAGAGTTTTCCAGAGGGTTTTAAGTGGGAGCGTGTGATGAACTTTGAGGATGGTGGTGCAGTGACAGTGACACAGGATACAAGTCTTGAGGATGGTACACTTATTTACAAGGTGAAGCTTCGTGGTACAAACTTTCCACCAGATGGTCCAGTGATGCAGAAGAAGACAATGGGTTGGGAGGCAAGTACAGAGCGTCTTTACCCAGAGGATGGTGTGCTTAAGGGTGATATTAAGATGGCACTTCGTCTTAAGGATGGTGGTCGTTACCTTGCAGATTTTAAGACAACATACAAGGCAAAGAAGCCAGTGCAGATGCCAGGTGCATACAACGTGGATCGTAAGCTTGATATTACAAGTCACAACGAGGATTACACAGTGGTGGAGCAGTACGAGCGTAGTGAGGGTCGTCACAGTACAGGTGGTATGGATGAGCTTTACAAGCCTAGGACAAGTTATCCATATGATAATCCAGATTATGCACCAGTTGCAACATTAGGTACCATGGAAAAGAAAAGTAGTTATAAAACAGTTTTATTAGGAGAATCATCAGTAGGAAAATCAAGTATAGTTTTACGATTAACAAAAGATACTTTTCATGAGAATACTAACACAACAATAGGTGCATCTTTTTGTACATATGTAGTAAATTTAAATGATATAAATATAAAGAATAATAGTAATAATGAAAAAAATAATAATATAAATTCAATAAATGATGATAATAATGTTATTATAACAAATCAACATAATAATTATAATGAAAACTTATGTAATATAAAATTTGATATATGGGATACAGCTGGACAAGAAAGATATGCAAGTATTGTCCCACTATATTATAGAGGTGCTACTTGTGCTATAGTAGTTTTTGATATTAGCAATTCAAATACTTTAGATCGAGCTAAGACATGGGTTAATCAATTAAAAATTAGTAGTAACTATATTATTATTTTAGTTGCTAATAAAATAGATAAAAACAAATTCCAAGTTGATATATTAGAAGTACAAAAATATGCTCAAGACAATAATTTACTTTTTATTCAAACAAGTGCCAAAACTGGAACAAACATAAAAAATATATTCTACATGCTTGCTGAAGAAATTTATAAAAATATTATAAATAATAATAATACCTCAAAAAATAAAACAGTTAATAAAAACTTAATAAATCTAGATAATCAAACACTTTCAAAAAAAGGATGTTGTTAACCCGGGTCGAGGGATATGGCAGCTTAATGTTCGTTTTTCTTATTTATATATTTATACCAATTGATTGTATTTATAACTGTAAAAATGTGTATGTTGTGTGCATATTTTTTTTTGTGCATGCACATGCATGTAAATAGCTAAAATTATGAACATTTTATTTTTTGTTCAGAAAAAAAAAACTTTACACACATAAAATGGCTAGTATGAATAGCCATATTTTATATAAATTAAATCCTATGAATTTATGACCATATTAAAAATTTAGATATTTATGGAACATAATATGTTTGAAACAATAAGACAAAATTATTATTATTATTATTATTTTTACTGTTATAATTATGTGTCTCCTTCAATGATTCATAAATAGTTGGACTTGATTTTTAAAATGTTTATAATATGATTAGCATAGTTAAATAAAAAAAGTTGAAAAATTAAAAAAAAACATATAAACACAAATGATGGTTTTTCCTTCAATTTCGATATCAATTTATAGAAACAAAATATATACTTGTATAATTTTATTTTTTTATATAAATCATTACATATATAATTATACAATATTTTTTCTAAGAGATAATTATATATTAATATATATAAAAAAAGGTGTTTTTTTTTTTTTTTTTTATTTTTATTTTTATTTTATGGTAATATTTTATTTTCCTTATTTTATAAATTATATTAGTTTATATGTGATTAATTTTATATATTATCAATTTATATATTTTTAAATGCTTACTTAATTATCTTTTTTTTTTTTTTTTTTTTTTTTTCCCCTCTTTTTATATTAATTTATTTTTGAAAAAATTGATATATATATAGTTGAAATATAAATTTCAAAAAAAATGATCACAAAATATACACTTAAATATAGGTACAATAAAAAAAAAAAATAAAAATATAATTACAAGATAATATTTTTTCCTGCTATCAAATTTTTATATATTCTCCTCAAAGAAAAAATATAAATAAATGAAGTAAATTAAAAAAAAAATTTTCTTTTTCTTCTTCTTTTGTAATTCCTTATTTATACATATTTTACTATATTTTCATAAAAATAAATTGTCATATTATATAAATATATATACCAAACCATAATTATATAGCCCTCAACATATTTTTATGATGTTTTTTCTTTTTAAATGTGATACGTAATTAAATATAATAATATATATTAAATATTATATTTTGTAATATTTACTTTCATGAGGTTTAATAAATTATAAAGAGAGAATAAAAAAAAAAAAAAAAAAAATTTATCCATATACAAATTATTAATTTATTTTTATTTTTTATTTCCCTTTGTATATATTATAAAAAAATAATCTACATAATTTTATATGATGATAATTACAATATAATATTTATAATATATATTATTTGTTAAGAGAAAAAAAAAATAAATATATACCTTCTTTTTGAGAATTGAATAAATTGTTTAATATATATATATATATATATAAATATATTATACATTTATGGTGAAAAAAAATGTTGTATTTAATTAGATTTAATATATATATATAAAAAATAGCGTATTTAAAATAATATATATATATATATATTATTATTATTACAAAATGACGGATATTATAAAAGTATATATCTATATATATGTATATATATATAATATTATTTTACTATATATATATAATATATAAATGTAAATGCATATAGTATCTATGTATATTATATATATATATAATATTAATTACATATCTAAGTTTTCTTTTCTTTTCTTTTTTTTTTTTTTTTTTTTTATTTTTTATAGAGAGCCCGTTATATATTATATATTAAAATTTTTTATAATAGCATTTATATACAATATTTTATTAACTAAAAGAAAAAAAAAAAAAAAAAAAAAAAAAAAACGAGGAAATTTATATTTCTTTAACAACATTTTAATTAAAATCATGATAATACAATTTCAAATCATTTTGTAATTATATAAATAAATATATATATATATATATATATATATATGTACTTTTAAATTAGGAATATTCTCATTTATAAATATATCTTATTTTTTAAATTGGTATAAAAAAAAAAAAAAAAATAAGAAACCGTTGATTAAATAATACATATATAATATAAATATATTTTATAAATATATATTTATATATATATATATATATATTTATAACGTATATCATTTTAAAGATAAACTAGTATGGCACCAAAAAAAAAAAGAAAAGTTACGCGTGATCCAACAAGAAGTGCAAATAGTGGAGCAGGAGCAGGAGCAGGAGCAATATTAAGTAGAGCTAGCATGGCTTCTAGAATCCTCTGGCATGAGATGTGGCATGAAGGCCTGGAAGAGGCATCTCGTTTGTACTTTGGGGAAAGGAACGTGAAAGGCATGTTTGAGGTGCTGGAGCCCTTGCATGCTATGATGGAACGGGGCCCCCAGACTCTGAAGGAAACATCCTTTAATCAGGCCTATGGTCGAGATTTAATGGAGGCCCAAGAGTGGTGCAGGAAGTACATGAAATCAGGGAATGTCAAGGACCTCCTCCAAGCCTGGGACCTCTATTATCATGTGTTCCGACGAATCTCAAAGGGTGAGGGTCGTGGTTCACTTCTTACTTGCGGTGACGTTGAGGAGAACCCTGGTCCTGTCGACATGACAGCCAGTTTAACTACCAAGTTCTTGAACAATACCTATGAAAACCCATTTATGAATGCATCCGGTGTTCATTGCATGACTACACAAGAATTAGATGAATTAGCAAACTCTAAAGCTGGCGCATTCATTACAAAGAGTGCTACAACCTTAGAAAGAGAAGGTAACCCTGAACCACGTTACATTTCTGTCCCTCTAGGCAGTATCAACTCCATGGGTTTACCAAACGAAGGTATCGACTACTATTTGTCCTATGTATTAAACCGTCAAAAGAATTATCCTGATGCACCTGCTATTTTCTTCTCAGTTGCTGGTATGAGCATTGATGAAAATTTAAATTTGTTGAGGAAAATCCAAGATAGCGAATTCAACGGTATTACCGAGTTAAACTTGTCTTGTCCTAATGTGCCTGGGAAACCACAAGTTGCTTATGACTTTGACTTGACAAAGGAAACCTTGGAAAAGGTTTTTGCCTTTTTCAAAAAACCTCTTGGTGTCAAGTTGCCTCCTTATTTTGATTTTGCCCATTTTGATATCATGGCAAAAATATTGAACGAGTTCCCATTAGCTTATGTCAACTCTATCAATAGTATAGGAAATGGTCTTTTCATTGATGTGGAGAAGGAGAGTGTAGTAGTGAAGCCAAAGAATGGTTTCGGGGGTATTGGAGGTGAATATGTTAAGCCAACCGCGCTCGCCAATGTTCGTGCATTTTACACTCGTTTGAGACCTGAAATCAAAGTTATCGGTACAGGTGGAATTAAGTCCGGTAAGGATGCATTTGAACATCTTCTATGTGGTGCCTCTATGCTACAGATTGGTACAGAATTACAAAAAGAGGGCGTCAAGATTTTTGAACGTATCGAAAAAGAATTAAAAGACATAATGGAAGCTAAGGGTTATACATCCATAGATCAGTTCCGTGGGAAGTTGAACAGCATTTAAAAGCTTATTTAATAATAGATTAAAAATATTATAAAAATAAAAACATAAACACAGAAATTACAAAAAAAATACATATGAATTTTTTTTTTGTAATCTTCCTTATAAATATAGAATAATGAATCATATAAAACATATCATTATTCATTTATTTACATTTAAAATTATTGTTTCAGTATCTTTAATTTATTATGTATATATAAAAATAACTTACAATTTTATTAATAAACAATATATGTTTATTAATTCATGTTTTGTAATTTATGGGATAGCGATTTTTTTTACTGTCTGTATTTTTCTTTTTTAATTATGTTTTAATTGTATTTTATTTTTATTATTGTTCTTTTTATAGTATTATTTTAAAACAAAATGTATTTTCTAAGAACTTATAATAATAATAAATATAAATTTTAATAAAAATTATATTTATCTTTTACAATATGAACATAAAGTACAACATTAATATATAGCTTTTAATATTTTTATTCCTAATCATGTAAATCTTAAATTTTTCTTTTTAAACATATGTTAAATATTTATTTCTCATTATATATAAGAACATATTTATTAAATCTAGAATTCTATAGTGAGTCGTATTACAATTCACTGGCCGTCGTTTTACAACGTCGTGACTGGGAAAACCCTGGCGTTACCCAACTTAATCGCCTTGCAGCACATCCCCCTTTCGCCAGCTGGCGTAATAGCGAAGAGGCCCGCACCGATCGCCCTTCCCAACAGTTGCGCAGCCTGAATGGCGAATGGCGCCTGATGCGGTATTTTCTCCTTACGCATCTGTGCGGTATTTCACACCGCATATGGTGCACTCTCAGTACAATCTGCTCTGATGCCGCATAGTTAAGCCAGCCCCGACACCCGCCAACACCCGCTGACGCGCCCTGACGGGCTTGTCTGCTCCCGGCATCCGCTTACAGACAAGCTGTGACCGTCTCCGGGAGCTGCATGTGTCAGAGGTTTTCACCGTCATCACCGAAACGCGCGAGACGAAAGGGCCTCGTGATACGCCTATTTTTATAGGTTAATGTCATGATAATAATGGTTTCTTAGACGTCAGGTGGCACTTTTCGGGGAAATGTGCGCGGAACCCCTATTTGTTTATTTTTCTAAATACATTCAAATATGTATCCGCTCATGAGACAATAACCCTGATAAATGCTTCAATAATATTGAAAAAGGAAGAGTATGAGTATTCAACATTTCCGTGTCGCCCTTATTCCCTTTTTTGCGGCATTTTGCCTTCCTGTTTTTGCTCACCCAGAAACGCTGGTGAAAGTAAAAGATGCTGAAGATCAGTTGGGTGCACGAGTGGGTTACATCGAACTGGATCTCAACAGCGGTAAGATCCTTGAGAGTTTTCGCCCCGAAGAACGTTTTCCAATGATGAGCACTTTTAAAGTTCTGCTATGTGGCGCGGTATTATCCCGTATTGACGCCGGGCAAGAGCAACTCGGTCGCCGCATACACTATTCTCAGAATGACTTGGTTGAGTACTCACCAGTCACAGAAAAGCATCTTACGGATGGCATGACAGTAAGAGAATTATGCAGTGCTGCCATAACCATGAGTGATAACACTGCGGCCAACTTACTTCTGACAACGATCGGAGGACCGAAGGAGCTAACCGCTTTTTTGCACAACATGGGGGATCATGTAACTCGCCTTGATCGTTGGGAACCGGAGCTGAATGAAGCCATACCAAACGACGAGCGTGACACCACGATGCCTGTAGCAATGCCAACAACGTTGCGCAAACTATTAACTGGCGAACTACTTACTCTAGCTTCCCGGCAACAATTAATAGACTGGATGGAGGCGGATAAAGTTGCAGGACCACTTCTGCGCTCGGCCCTTCCGGCTGGCTGGTTTATTGCTGATAAATCTGGAGCCGGTGAGCGTGGGTCTCGCGGTATCATTGCAGCACTGGGGCCAGATGGTAAGCCCTCCCGTATCGTAGTTATCTACACGACGGGGAGTCAGGCAACTATGGATGAACGAAATAGACAGATCGCTGAGATAGGTGCCTCACTGATTAAGCATTGGTAACTGTCAGACCAAGTTTACTCATATATACTTTAGATTGATTTAAAACTTCATTTTTAATTTAAAAGGATCTAGGTGAAGATCCTTTTTGATAATCTCATGACCAAAATCCCTTAACGTGAGTTTTCGTTCCACTGAGCGTCAGACCCCGTAGAAAAGATCAAAGGATCTTCTTGAGATCCTTTTTTTCTGCGCGTAATCTGCTGCTTGCAAACAAAAAAACCACCGCTACCAGCGGTGGTTTGTTTGCCGGATCAAGAGCTACCAACTCTTTTTCCGAAGGTAACTGGCTTCAGCAGAGCGCAGATACCAAATACTGTCCTTCTAGTGTAGCCGTAGTTAGGCCACCACTTCAAGAACTCTGTAGCACCGCCTACATACCTCGCTCTGCTAATCCTGTTACCAGTGGCTGCTGCCAGTGGCGATAAGTCGTGTCTTACCGGGTTGGACTCAAGACGATAGTTACCGGATAAGGCGCAGCGGTCGGGCTGAACGGGGGGTTCGTGCACACAGCCCAGCTTGGAGCGAACGACCTACACCGAACTGAGATACCTACAGCGTGAGCTATGAGAAAGCGCCACGCTTCCCGAAGGGAGAAAGGCGGACAGGTATCCGGTAAGCGGCAGGGTCGGAACAGGAGAGCGCACGAGGGAGCTTCCAGGGGGAAACGCCTGGTATCTTTATAGTCCTGTCGGGTTTCGCCACCTCTGACTTGAGCGTCGATTTTTGTGATGCTCGTCAGGGGGGCGGAGCCTATCGAAAAACGCCAGCAACGCGGCCTTTTTACGGTTCCTGGCCTTTTGCTGGCCTTTTGCTCACATGTTCTTTCCTGCGTTATCCCCTGATTCTGTGGATAACCGTATTACCGCCTTTGAGTGAGCTGATACCGCTCGCCGCAGCCGAACGACCGAGCGCAGCGAGTCAGTGAGCGAGGAAGCGGAAGAGCGCCCAATACGCAAACCGCCTCTCCCCGCGCGTTGGCCGATTCATTAATGCAGCTGGCACGACAGGTTTCCCGACTGGAAAGCGGGCAGTGAGCGCAACGCAATTAATGTGAGTTAGCTCACTCATTAGGCACCCCAGGCTTTACACTTTATGCTTCCGGCTCGTATGTTGTGTGGAATTGTGAGCGGATAACAATTTCACACAGGAAACAGCTATGACCATGATTACGCCAAGCTATTTAGGTGACACTATAGAATACTC

>pSLI yDHODH-T2A-2xFKBP-GFP-2xFKBP-POI^loxP^ / ^Pfcam^hDHFR

Homology Region Rab11b (gDNA)) STOP codon 2xMyc LoxP FKBP35 Linker GFP T2A (skip peptide) yDHODH (DSM1) recodonized full length Rab11b Pfcam promotor hDHFR (WR) *hsp86 3’* *amp* promotor+Ampiclin resitance replication origin

taatgtgagttagctcactcattaggcaccccaggctttacactttatgcttccggctcgtatgttgtgtggaattgtgagcggataacaatttcacacaggaaacagctatgaccatgattacgccaagctatttaggtgacactatagaatactcgcggccgcTAATATCGAAGTATAACTAGCGCACATTATAGGAGAAGCGCAGGGGCTATATTAGTATATGATATAACTAAGAAGAAAACATTTTTAAGTATATCAAAATGGTTAGAAGAAATAAGACAAAATGCAGATAAGGATATTGTCATTATGCTTGTTGGAAATAAGGTAGATTTAACTGAAGAAGACGAAACTAAAAGGAAGgtaagaaaataaaaagatatgcatatgtacatgtacatatttttttttattattattatgattttatttttttttttttttatttttttttttttttatttttttttggtgaagGTGACATATGAACAAGGCGCAAACTTTGCAAGAGAAAATAATTTATTCTTTGCAGAGGCCTCCGCTGTATCTAAATTAAATGTAAAACATATATTTGAAAATTTATTACAAGAAATATATAATAACAGATTAAAAAATAATAATCGAAGCTTTAGTAATAGAAGTGTAGCTACGTGCGAAAGCGCAATACAACTCACCAAAGCCAGGAGTGTCATAAAATTAAATGAGGTATATGATAATCAGAGTGAAGATGGTTTAAACGAGCAGAAGTTAATATCAGAAGAGGATTTGGGTGAACAAAAACTCATAAGCGAAGAAGATTTA ATAACTTCGTATAGCATACATTATACGAAGTTATTCCGGAGAAGGAAGAGGAAGTTTATTAACATGTGGAGATGTAGAAGAAAATCCAGGACCAATGACAGCCAGTTTAACTACCAAGTTCTTGAACAATACCTATGAAAACCCATTTATGAATGCATCCGGTGTTCATTGCATGACTACACAAGAATTAGATGAATTAGCAAACTCTAAAGCTGGCGCATTCATTACAAAGAGTGCTACAACCTTAGAAAGAGAAGGTAACCCTGAACCACGTTACATTTCTGTCCCTCTAGGCAGTATCAACTCCATGGGTTTACCAAACGAAGGTATCGACTACTATTTGTCCTATGTATTAAACCGTCAAAAGAATTATCCTGATGCACCTGCTATTTTCTTCTCAGTTGCTGGTATGAGCATTGATGAAAATTTAAATTTGTTGAGGAAAATCCAAGATAGCGAATTCAACGGTATTACCGAGTTAAACTTGTCTTGTCCTAATGTGCCTGGGAAACCACAAGTTGCTTATGACTTTGACTTGACAAAGGAAACCTTGGAAAAGGTTTTTGCCTTTTTCAAAAAACCTCTTGGTGTCAAGTTGCCTCCTTATTTTGATTTTGCCCATTTTGATATCATGGCAAAAATATTGAACGAGTTCCCATTAGCTTATGTCAACTCTATCAATAGTATAGGAAATGGTCTTTTCATTGATGTGGAGAAGGAGAGTGTAGTAGTGAAGCCAAAGAATGGTTTCGGGGGTATTGGAGGTGAATATGTTAAGCCAACCGCGCTCGCCAATGTTCGTGCATTTTACACTCGTTTGAGACCTGAAATCAAAGTTATCGGTACAGGTGGAATTAAGTCCGGTAAGGATGCATTTGAACATCTTCTATGTGGTGCCTCTATGCTACAGATTGGTACAGAATTACAAAAAGAGGGCGTCAAGATTTTTGAACGTATCGAAAAAGAATTAAAAGACATAATGGAAGCTAAGGGTTATACATCCATAGATCAGTTCCGTGGGAAGTTGAACAGCATTGGTGAAGGTAGAGGTTCTTTGTTGACTTGTGGTGATGTTGAAGAAAATCCAGGTCCAGCTAGCCGTGGTGTTCAGGTCGAGACTATTAGCCCTGGAGATGGACGCACGTTTCCTAAGCGTGGACAGACATGCGTAGTTCACTACACAGGTATGTTGGAGGACGGTAAAAAGTTCGACAGCTCACGCGACCGCAATAAACCTTTCAAGTTTATGCTTGGCAAGCAGGAGGTTATTCGTGGATGGGAGGAGGGTGTAGCACAGATGTCTGTTGGACAGCGTGCTAAGTTGACAATTTCACCTGACTATGCTTATGGCGCTACGGGCCATCCCGGGATCATTCCGCCACATGCGACTCTGGTATTCGACGTTGAATTATTAAAGTTAGAGACAGCTAGAGGGGCCGCTGCAGGTGCTGGTGGAGCTGGAAGACGTGGAGTACAAGTAGAGACTATCTCTCCAGGTGACGGTCGCACTTTCCCAAAGCGTGGCCAAACCTGTGTTGTACATTACACTGGTATGCTGGAGGATGGGAAAAAGTTCGATTCCAGTCGCGACCGTAACAAACCGTTCAAATTCATGTTGGGAAAGCAGGAAGTGATCCGCGGGTGGGAGGAAGGCGTGGCGCAAATGAGCGTCGGTCAGCGGGCTAAATTGACCATTTCCCCTGACTACGCGTATGGGGCTACTGGGCACCCAGGGATTATTCCGCCTCACGCTACACTTGTGTTTGATGTCGAACTTTTGAAACTGGAAACTGCCAGGGGAGCAGCCGCAGGAGCAGGGGGGGCAGGAAGGGTCGACATGAGTAAAGGAGAAGAACTTTTCACTGGAGTTGTCCCAATTCTTGTTGAATTAGATGGTGATGTTAATGGGCACAAATTTTCTGTCAGTGGAGAGGGTGAAGGTGATGCAACATACGGAAAACTTACCCTTAAATTTATTTGCACTACTGGAAAACTACCAGTTCCATGGCCAACACTTGTCACTACTTTCGCGTATGGTCTTCAATGCTTTGCGAGATACCCAGATCATATGAAACAGCATGACTTTTTCAAGAGTGCCATGCCCGAAGGTTATGTACAGGAAAGAACTATATTTTTCAAAGATGACGGGAACTACAAGACACGTGCTGAAGTCAAGTTTGAAGGTGATACCCTTGTTAATAGAATCGAGTTAAAAGGTATTGATTTTAAAGAAGATGGAAACATTCTTGGACACAAATTGGAATACAACTATAACTCACACAATGTATACATCATGGCAGACAAACAAAAGAATGGAATCAAAGTTAACTTCAAAATTAGACACAACATTGAAGATGGAAGCGTTCAACTAGCAGACCATTATCAACAAAATACTCCAATTGGCGATGGCCCTGTCCTTTTACCAGACAACCATTACCTGTCCACACAATCTGCCCTTTCGAAAGATCCCAACGAAAAGAGAGACCACATGGTCCTTCTTGAGTTTGTAACAGCTGCTGGGATTACACATGGCATGGATGAGCTCTACAAACCTAGCTCAGGATTGAGATCAAGATCTGCTGCTGCTGGTGCTGGTGGTGCTGCTAGAGCTGCTCTGCAGAGAGGAGTACAAGTTGAAACAATATCACCAGGAGATGGTCGTACATTTCCAAAAAGAGGTCAAACTTGTGTTGTACATTATACTGGAATGCTTGAAGATGGAAAGAAATTTGATTCATCTCGTGATAGAAATAAACCATTTAAATTTATGCTAGGTAAACAAGAAGTAATACGAGGTTGGGAAGAAGGAGTTGCTCAAATGAGTGTAGGTCAAAGAGCAAAACTTACTATATCTCCAGATTATGCTTATGGTGCAACTGGACATCCAGGTATAATTCCACCTCATGCAACTCTTGTATTTGATGTGGAGCTTCTAAAACTAGAAACTAGAGGTGTTCAGGTTGAAACAATTTCACCTGGAGATGGCAGAACCTTTCCTAAAAGAGGACAGACTTGCGTAGTTCATTATACAGGCATGCTAGAGGATGGTAAGAAATTTGATTCTAGTCGAGATAGAAATAAGCCATTCAAGTTTATGCTAGGTAAACAGGAAGTAATAAGAGGTTGGGAAGAGGGTGTAGCACAGATGTCAGTTGGACAAAGAGCAAAGTTAACAATATCACCAGATTATGCATACGGTGCAACAGGCCATCCTGGCATCATCCCTCCACATGCAACTTTAGTATTCGACGTTGAATTGTTAAAGTTAGAGACAACGCGTGCTAGAGGTGCTGCTGCTGGTGCTGGAGGTGCAGGTAGA

CCTAGGAGCAACGAGGAGTATGACCACTTATACAAGATTATACTTGTTGGAGATGCTACTGTTGGAAAAACTCACCTTTTGAGTCGTTACATTCGTGGTAGCTTACCATCAGTTGCTAAGGCAACAATAGGAGTTGAGTTCGCAACACGTACAATACCTCTTGCTGTTGGAGGAACAGTTAAGGCACAAATTTGGGACACAGCAGGACAGGAGAGGTACAGGTCAATTACATCAGCTCACTACCGTCGTTCAGCTGGAGCAATTCTTGTTTACGACATTACAAAAAAAAAGACTTTCCTTTCAATTAGTAAGTGGCTTGAGGAGATTCGTCAGAACGCTGACAAAGACATAGTTATAATGTTAGTAGGTAACAAAGTTGACCTTACAGAGGAGGATGAGACAAAGCGTAAAGTTACTTACGAGCAGGGAGCTAATTTCGCTCGTGAGAACAACCTTTTTTTCGCTGAAGCAAGTGCAGTTAGCAAGCTTAACGTTAAGCACATTTTCGAGAACCTTCTTCAGGAGATTTACAACAATCGTCTTAAGAACAACAACAGGTCATTCTCAAACCGTTCAGTTGCAACATGTGAGTCAGCTATTCAGTTAACAAAGGCACGTTCAGTTATTAAGCTTAACGAAGTTTACGACAACCAATCAGAGGACAACAACATGAACAAGGTTAAGTGCTGCTGAAGGCCTataacttcgtatagcatacattatacgaagttattatgactcgagggatatggcagcttaatgttcgtttttcttatttatatatttataccaattgattgtatttataactgtaaaaatgtgtatgttgtgtgcatatttttttttgtgcatgcacatgcatgtaaatagctaaaattatgaacattttattttttgttcagaaaaaaaaaactttacacacataaaatggctagtatgaatagccatattttatataaattaaatcctatgaatttatgaccatattaaaaatttagatatttatggaacataatatgtttgaaacaataagacaaaattattattattattattatttttactgttataattatgtgtctccttcaatgattcataaatagttggacttgatttttaaaatgtttataatatgattagcatagttaaataaaaaaagttgaaaaattaaaaaaaaacatataaacacaaatgatggtttttccttcaatttcgatatcaatttatagaaacaaaatatatacttgtataattttatttttttatataaatcattacatatataattatacaatattttttctaagagataattatatattaatatatataaaaaaaggtgttttttttttttttttttatttttatttttattttatggtaatattttattttccttattttataaattatattagtttatatgtgattaattttatatattatcaatttatatatttttaaatgcttacttaattatctttttttttttttttttttttttttcccctctttttatattaatttatttttgaaaaaattgatatatatatatatatataatatatatatatacatgtagtagtattaaacaatgtataatatatataaataatatatttatatatttcatttcaattttaattttttttggttttttttttttttctttttgtcatatttaaaaaaaattatattcatataagttatgcattttttataaacattattcaatatatgtataatataatatatatatatatattaatgtattattccaatgtgcatgataaaagaaaaaaataatatttataaaaaaaaagaaaaataaaacaaaaaaagaaaaaaaaaaaaaaaaaaaaaaaaatacaaaaataaataatataatttataattatatattcttgtcacaataaaaatatatatatatatatatatatttataatatgtatattttaaactagaaaaggaataactaatattttatttattatcattcaagatttatattttataataataaatacctaatagaaatatatcaggatccATGCATGGTTCGCTAAACTGCATCGTCGCTGTGTCCCAGAACATGGGCATCGGCAAGAACGGGGACTACCCCTGGCCACCGCTCAGGAACGAATTTAGATATTTCCAGAGAATGACCACAACCTCTTCAGTAGAAGGTAAACAGAATCTGGTGATTATGGGTAAGAAGACCTGGTTCTCCATTCCTGAGAAGAATCGACCTTTAAAGGGTAGAATTAATTTAGTTCTCAGCAGAGAACTCAAGGAACCTCCACAAGGAGCTCATTTTCTTTCCAGAAGTCTAGATGATGCCTTAAAACTTACTGAACAACCAGAATTAGCAAATAAAGTAGACATGGTCTGGATAGTTGGTGGCAGTTCTGTTTATAAGGAAGCCATGAATCACCCAGGCCATCTTAAACTATTTGTGACAAGGATCATGCAAGACTTTGAAAGTGACACGTTTTTTCCAGAAATTGATTTGGAGAAATATAAACTTCTGCCAGAATACCCAGGTGTTCTCTCTGATGTCCAGGAGGAGAAAGGCATTAAGTACAAATTTGAAGTATATGAGAAGAATGATTAA*gcttatttaataatagattaaaaatattataaaaataaaaacataaacacagaaattacaaaaaaaatacatatgaattttttttttgtaatcttccttataaatatagaataatgaatcatataaaacatatcattattcatttatttacatttaaaattattgtttcagtatctttaatttattatgtatatataaaaataacttacaattttattaataaacaatatatgtttattaattcatgttttgtaatttatgggatagcgattttttttactgtctgtatttttcttttttaattatgttttaattgtattttatttttattattgttctttttatagtattattttaaaacaaaatgtattttctaagaacttataataataataaatataaattttaataaaaattatatttatcttttacaatatgaacataaagtacaacattaatatatagcttttaatatttttattcctaatcatgtaaatcttaaatttttctttttaaacatatgttaaatatttatttctcattatatataagaacatatttattaaatctagaatt*ctatagtgagtcgtattacaattcactggccgtcgttttacaacgtcgtgactgggaaaaccctggcgttacccaacttaatcgccttgcagcacatccccctttcgccagctggcgtaatagcgaagaggcccgcaccgatcgcccttcccaacagttgcgcagcctgaatggcgaatggcgcctgatgcggtattttctccttacgcatctgtgcggtatttcacaccgcatatggtgcactctcagtacaatctgctctgatgccgcatagttaagccagccccgacacccgccaacacccgctgacgcgccctgacgggcttgtctgctcccggcatccgcttacagacaagctgtgaccgtctccgggagctgcatgtgtcagaggttttcaccgtcatcaccgaaacgcgcgagacgaaagggcctcgtgatacgcctatttttataggttaatgtcatgataataatggtttcttagacgtcaggtggcacttttcggggaaatgtg

cgcggaacccctatttgtttatttttctaaatacattcaaatatgtatccgctcatgagacaataaccctgataaatgcttcaataatattgaaaaaggaagagt

ATGAGTATTCAACATTTCCGTGTCGCCCTTATTCCCTTTTTTGCGGCATTTTGCCTTCCTGTTTTTGCTCACCCAGAAACGCTGGTGAAAGTAAAAGATGCTGAAGATCAGTTGGGTGCACGAGTGGGTTACATCGAACTGGATCTCAACAGCGGTAAGATCCTTGAGAGTTTTCGCCCCGAAGAACGTTTTCCAATGATGAGCACTTTTAAAGTTCTGCTATGTGGCGCGGTATTATCCCGTATTGACGCCGGGCAAGAGCAACTCGGTCGCCGCATACACTATTCTCAGAATGACTTGGTTGAGTACTCACCAGTCACAGAAAAGCATCTTACGGATGGCATGACAGTAAGAGAATTATGCAGTGCTGCCATAACCATGAGTGATAACACTGCGGCCAACTTACTTCTGACAACGATCGGAGGACCGAAGGAGCTAACCGCTTTTTTGCACAACATGGGGGATCATGTAACTCGCCTTGATCGTTGGGAACCGGAGCTGAATGAAGCCATACCAAACGACGAGCGTGACACCACGATGCCTGTAGCAATGCCAACAACGTTGCGCAAACTATTAACTGGCGAACTACTTACTCTAGCTTCCCGGCAACAATTAATAGACTGGATGGAGGCGGATAAAGTTGCAGGACCACTTCTGCGCTCGGCCCTTCCGGCTGGCTGGTTTATTGCTGATAAATCTGGAGCCGGTGAGCGTGGGTCTCGCGGTATCATTGCAGCACTGGGGCCAGATGGTAAGCCCTCCCGTATCGTAGTTATCTACACGACGGGGAGTCAGGCAACTATGGATGAACGAAATAGACAGATCGCTGAGATAGGTGCCTCACTGATTAAGCATTGGTAActgtcagaccaagtttactcatatatactttagattgatttaaaacttcatttttaatttaaaaggatctaggtgaagatcctttttgataatctcatgaccaaaatcccttaacgtgagttttcgttccactgagcgtcagaccccgtagaaaagatcaaaggatcttcttgagatcctttttttctgcgcgtaatctgctgcttgcaaacaaaaaaaccaccgctaccagcggtggtttgtttgccggatcaagagctaccaactctttttccgaaggtaactggcttcagcagagcgcagataccaaatactgtccttctagtgtagccgtagttaggccaccacttcaagaactctgtagcaccgcctacatacctcgctctgctaatcctgttaccagtggctgctgccagtggcgataagtcgtgtcttaccgggttggactcaagacgatagttaccggataaggcgcagcggtcgggctgaacggggggttcgtgcacacagcccagcttggagcgaacgacctacaccgaactgagatacctacagcgtgagctatgagaaagcgccacgcttcccgaagggagaaaggcggacaggtatccggtaagcggcagggtcggaacaggagagcgcacgagggagcttccagggggaaacgcctggtatctttatagtcctgtcgggtttcgccacctctgacttgagcgtcgatttttgtgatgctcgtcaggggggcggagcctatcgaaaaacgccagcaacgcggcctttttacggttcctggccttttgctggccttttgctcacatgttctttcctgcgttatcccctgattctgtggataaccgtattaccgcctttgagtgagctgataccgctcgccgcagccgaacgaccgagcgcagcgagtcagtgagcgaggaagcggaagagcgcccaatacgcaaaccgcctctccccgcgcgttggccgattcattaatgcagctggcacgacaggtttcccgactggaaagcgggcagtgagcgcaacgcaat
