## Supplementary material for "Characterization of Host Cell Cytosol-filled Vesicles in the Human Malaria Parasite *Plasmodium falciparum*": Table S3

**Table S3. Cryo-ET data collection**

|  | RBSN5 rapalog-treated | RBSN5 rapalog-treated (cryo-STET) | RBSN5 control |
| --- | --- | --- | --- |
| **Data collection and processing** |  |  |  |
| Magnification | 53,000 | 28,000 | 53,000 |
| Voltage (kV) | 300 | 300 | 300 |
| Total electron exposure (e–/Å^2^) | 120 | 180 | 120 |
| Defocus target (μm) | -3 | 0 | -3 |
| Pixel size (Å) | 1.695 | 10.84 | 1.695 |
| Tilt series range (°) | -53 to 67 | -60 to 60 | -53 to 67 |
| Starting tilt (°) | 7 | 0 | 7 |
| Tilt increment (°) | 3 | 2 | 3 |
| Tilt scheme | Dose-symmetric | Dose-symmetric | Dose-symmetric |
